# Autoantigen TRIM21 (Ro52) assembles pro-inflammatory immune complexes following lytic cell death

**DOI:** 10.1101/2024.09.06.611470

**Authors:** Esther L. Jones, Benjamin Demarco, Madelon M.E de Jong, Han Cai, Sarah Hill, Ryan E. Glass, Gemma Harris, Saba Nayar, Benjamin A. Fisher, Audrey Gérard, Jelena S. Bezbradica, Lynn B. Dustin

**Author notes:** Department of Medicine, University of Basel; Basel, Switzerland (Benjamin Demarco).

## Abstract

Sjögren’s disease (SjD) causes localised and systemic inflammation due to autoantibody production against intracellular proteins, such as TRIM21/Ro52. TRIM21 is an E3 ubiquitin ligase which binds antibody Fc domains on opsonised pathogens, which have escaped extracellular immunity and entered cytosols; TRIM21 ubiquitinates these, driving their proteasomal degradation. How and why TRIM21 becomes an autoantigen remains unclear. We show that TRIM21 is released upon lytic cell death (pyroptosis/necroptosis) but not apoptosis. Released TRIM21 binds circulating antibody Fc domains, and forms large immune complexes (ICs). These are further enhanced with TRIM21/Ro52 seropositive SjD plasma antibodies, where interactions are mediated via both Fc and F(ab’)_2_ domains. TRIM21-ICs are taken up by macrophages, which in high interferon environments drive pro-inflammatory responses, antigen presentation, and inflammatory and metabolic transcriptional changes. Whilst many cytosolic proteins are released by dead cells, due to its high affinity for antibodies, TRIM21 can generate large ICs. This may perpetuate inflammation and antigen presentation, causing TRIM21 to be highly autoimmunogenic.

**One Sentence Summary:** How the intracellular protein TRIM21 becomes an autoantigen.

## INTRODUCTION

Sjögren’s disease (SjD) is an autoimmune disease with a predicted global prevalence of 60 patients/100,000 population (*1*), and affecting an estimated 2-4 million people within the United States (*2*). This makes it the second most common autoimmune rheumatic disease after rheumatoid arthritis (*2, 3*). SjD is characterised by lymphocytic infiltration of the lacrimal and salivary glands (SGs), with symptoms including keratoconjunctivitis sicca (dry eyes) and xerostomia (dry mouth) (*4, 5*). Like many autoimmune diseases, clinical manifestations often extend to systemic involvements causing fatigue, chronic joint pain, numbness, and neurological, pulmonary, and renal sequelae (*6*). The elevated risk of B cell lymphoma is perhaps the most concerning complication associated with SjD. An estimated 5-10% of SjD patients develop mucosal-associated lymphoid tissue lymphomas, a group of low-grade non-Hodgkin lymphomas which are most commonly localised at the SGs (*7, 8*). Despite this clinical need, there are currently few therapeutic options, with no approved biological therapies (*9*).

Due to the diversity in symptoms, and overlaps in presentation with other diseases, diagnosing SjD is challenging (*10*). Certain autoantibodies are commonly present in patients, with anti-Ro/SSA (antibodies against TRIM21/Ro52 and TROVE2/Ro60) and anti-La/SSB being traditional biomarkers (*11*). Additionally, labial SG biopsies for histopathological scoring are important for SjD diagnoses (*12–14*). Whilst these autoantibodies and pathological presentations are known, little is understood about the pathogenic mechanisms driving autoantibody production, inflammation, and tissue damage.

One of the classical SjD autoantigens, TRIM21/Ro52/SSA1, is a common autoantigen in multiple autoimmune diseases. Its structure comprises an N-terminal RING domain, inhibitory B-box domain, predicted central coiled-coil domain and C-terminal PRYSPRY domain (*15*). Although PRYSPRY domains are found on multiple other proteins, and there are many TRIM proteins in humans, TRIM21 is unique in its ability to bind the Fc-portion of antibodies with high affinity (*16*). TRIM21 has well-defined roles as an intracellular E3 ubiquitin ligase. It senses and binds the antibodies of opsonised pathogens which have evaded extracellular immunity and entered the cytosol. Therefore, TRIM21 is sometimes referred to as an “intracellular Fc receptor” (FcR) (*17, 18*). TRIM21 then ubiquitinates both the target pathogen and antibody for degradation via the proteasome (*19*). TRIM21 tissue expression is ubiquitous, and particularly high at mucosal and immune sites including the respiratory system, gastrointestinal tract, reproductive tissues, bone marrow, and lymphoid organs (*20*). It remains unclear how and why this intracellular protein becomes an autoantigen.

Models for autoantibody production have often cited impaired clearance of apoptotic cell debris as a means for intracellular antigen exposure (*21, 22*). Others have referenced the uptake of antigen-containing apoptotic blebs by dendritic cells, which drives antigen presentation to facilitate autoimmune reactions (*23, 24*). However, apoptotic blebs are believed to prevent the inadvertent release of intracellular contents which could otherwise drive autoimmunity (*25*). Instead, forms of lytic cell death (pyroptosis, necroptosis or necrosis) promote immune responses through uncontrolled release of intracellular contents, often with concomitant release of inflammatory cytokines and alarmins. Excessive lytic cell death has been implicated in autoimmune diseases (*26*). In particular, pyroptosis has been suggested as a form of pro-inflammatory cell death which may contribute to SG damage and dysfunction in SjD (*27*). Evidence suggests macrophages expressing high levels of inflammasome products including caspase-1, interleukin (IL)-1β and IL-18 may infiltrate SjD patient SGs (*28*). However, as TRIM21 is expressed in most cells, unscheduled pro-inflammatory death of any cell could, in fact, be the source of this autoantigen. Additionally, SjD patients commonly present with overexpression of interferon (IFN)-induced genes, and elevated levels of serum type I IFNs. This is referred to as the “IFN signature” (*29, 30*). We hypothesise that this pro-inflammatory signature, combined with lytic cell death, may lead to TRIM21 release and autoimmune reactions for anti-TRIM21/Ro52 autoantibody production in SjD.

In this study, we focused on the classical SjD autoantigen, TRIM21/Ro52. We investigated potential mechanisms which predispose this protein to become an autoantigen and drive SjD pathogenesis. We found that TRIM21 is released upon lytic forms of cell death, rather than apoptosis. Upon its release, extracellular TRIM21 maintained its native antibody Fc-binding function, and formed higher-order immune complexes (IC) with plasma from TRIM21/Ro52-seropositive SjD patients. When taken up by macrophages, TRIM21-ICs induced antigen presentation, together with changes in pro-inflammatory and metabolic gene expression. These findings may go some way to explaining how TRIM21 contributes to inflammation and self-antigen presentation in SjD.

## RESULTS

### Cytosolic TRIM21 is ubiquitously expressed in both control and SjD patients

To determine whether TRIM21 is expressed in glands of SjD patients, SG biopsy slides from anti-TRIM21/Ro52 and/or anti-TROVE2/Ro60 seropositive SjD patients and sicca controls were obtained (Birmingham OASIS cohort). Slides were stained with H&E and analysed for the following: area, focus number and focus score (Table 1). Slides of adjacent sections were subjected to immunofluorescent staining.

**Table 1.** Sicca and SjD SG biopsy information. OASIS (Birmingham) FFPE biopsy slides were H&E stained and analysed for the following; area, foci and focal score. Information about “Ro/La” antibody expression, Sjögren’s and ARC2016 confirmation is provided.

| Slide | Ro | La | Sjögren's | ACR2016 | Area | Foci | Focus Score |
| --- | --- | --- | --- | --- | --- | --- | --- |
| 1 | 1 | 1 | 1 | 1 | 14623507.16 | 4 | 1.094128776 |
| 2 | 1 | 0 | 1 | 1 | *(Atrophy, could not evaluate)* |  |  |
| 8 | 1 | 1 | 1 | 1 | 8869240.456 | 4 | 1.803987622 |
| 12 | 1 | 0 | 1 | 1 | 4701817.812 | 2 | 1.701469585 |
| 14 | 1 | 1 | 1 | 1 | 15604053.98 | 7 | 1.794405482 |
| 4 | 0 | 0 | 0 | 0 | 16764352.01 | 2 | 0.477203055 |
| 5 | 0 | 0 | 0 | 0 | 15513490.62 | 0 | 0 |
| 18 | 0 | 0 | 0 | 0 | 10634976.03 | 0 | 0 |
| 20 | 0 | 0 | 0 | 0 | 11650822.04 | 2 | 0.686646828 |
| 21 | 0 | 0 | 0 | 0 | 6064994.634 | 0 | 0 |

Sicca control SG biopsies were anatomically normal, with clear glandular acini and ducts, as indicated by H&E and cytokeratin (CK) epithelial staining (Fig. 1A, i, ii). TRIM21 was ubiquitously expressed throughout the SG (Fig. 1A, iii). SjD biopsies (Fig. 1B, C) had areas of focal infiltration, demonstrated by higher focus scoring (≥1) (Table 1) compared to sicca controls, consistent with known literature and diagnostic criteria for the disease (*12, 31*). One representative SjD biopsy had severe tissue atrophy, with extensive adipose tissue replacement and glandular deformation, as commonly observed in SjD biopsies (*32*) (Table 1, Fig. 1B, i, ii). This SjD biopsy demonstrates the severe SG damage patients can present with, but was too atrophied to reliably assess TRIM21 expression. This was due to the absence of epithelia and tissue disintegration during repeated de-coverslipping for multiplex imaging (Fig. 1B, iii).

**Fig. 1.**
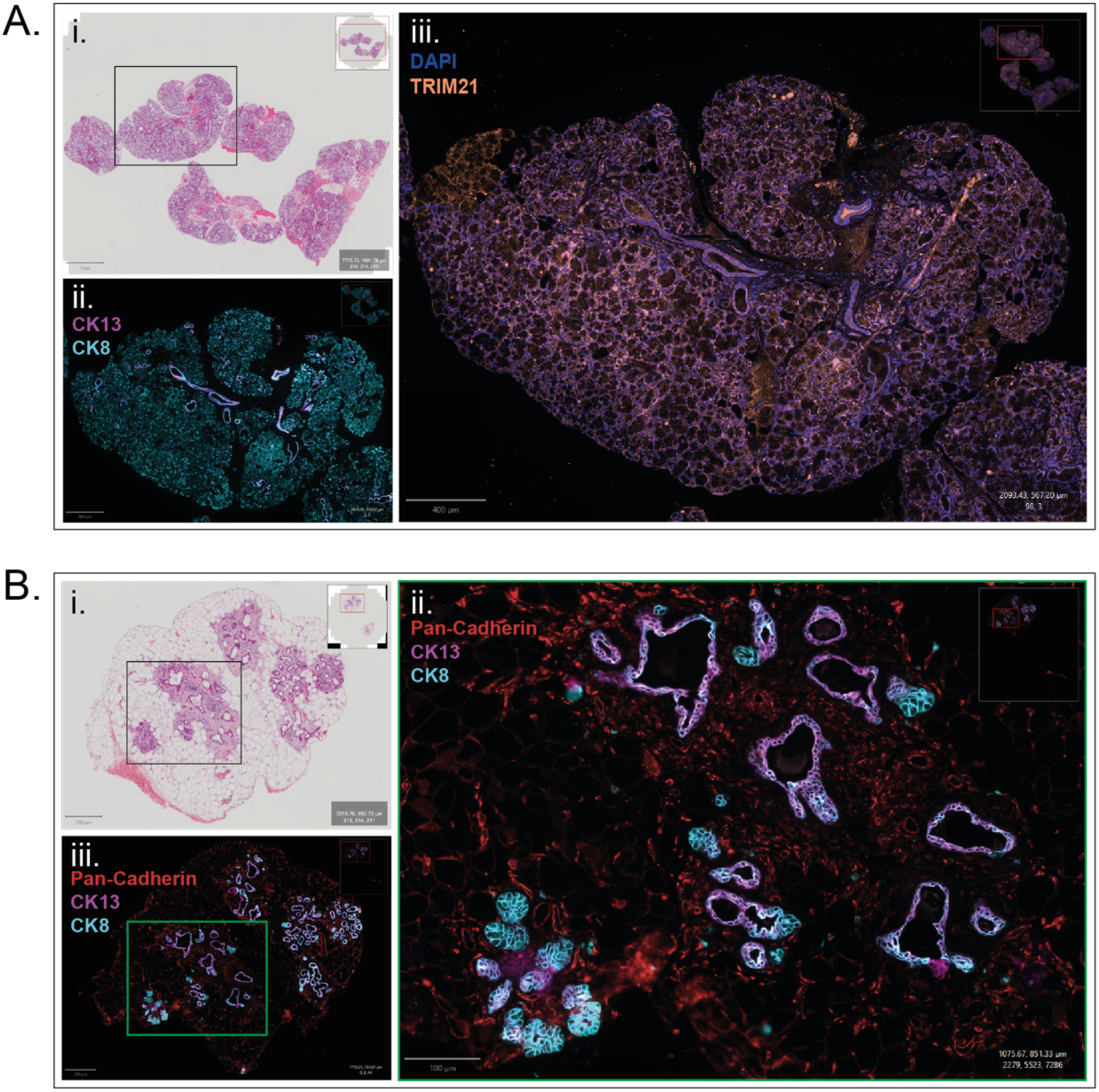
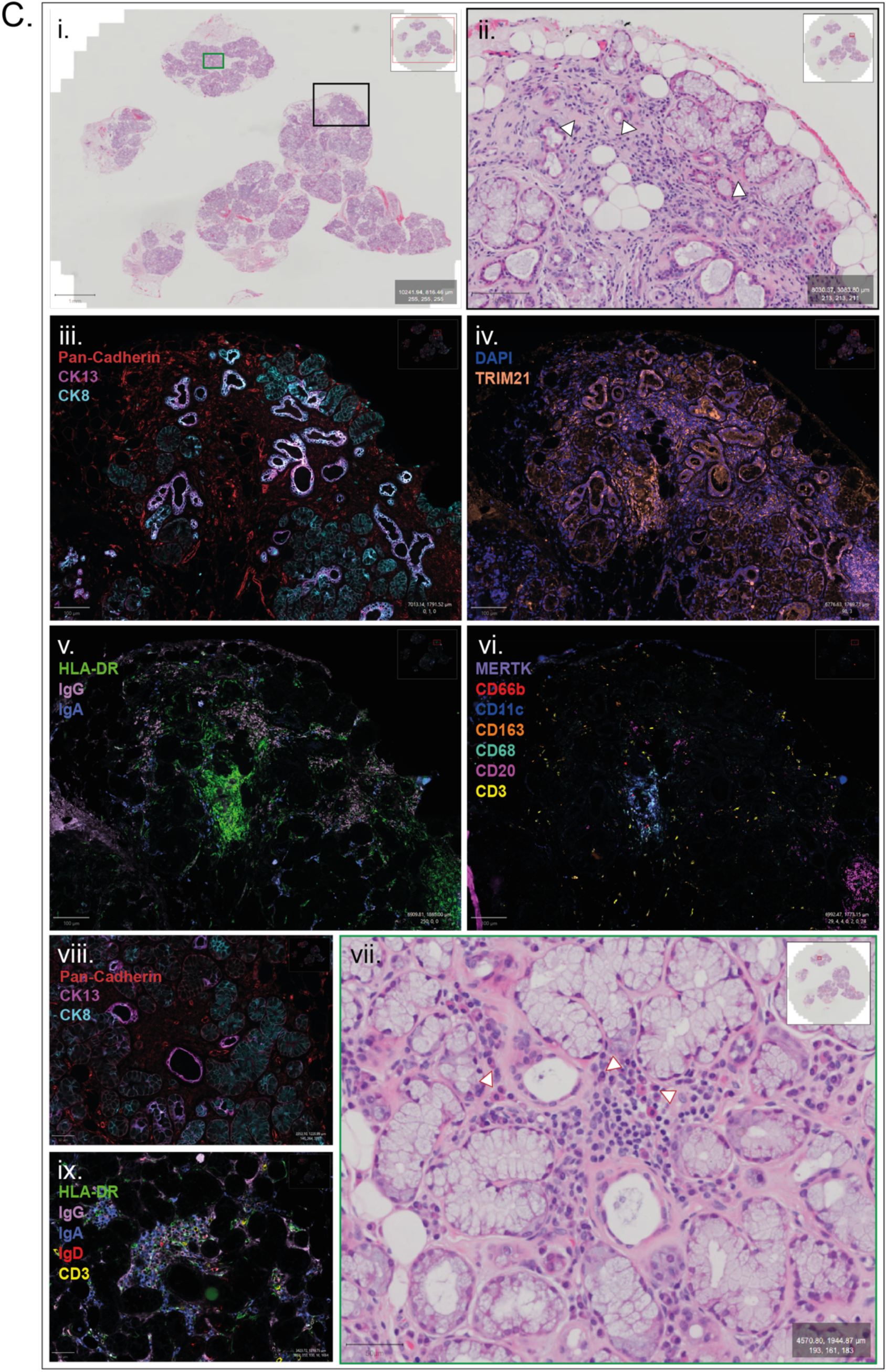
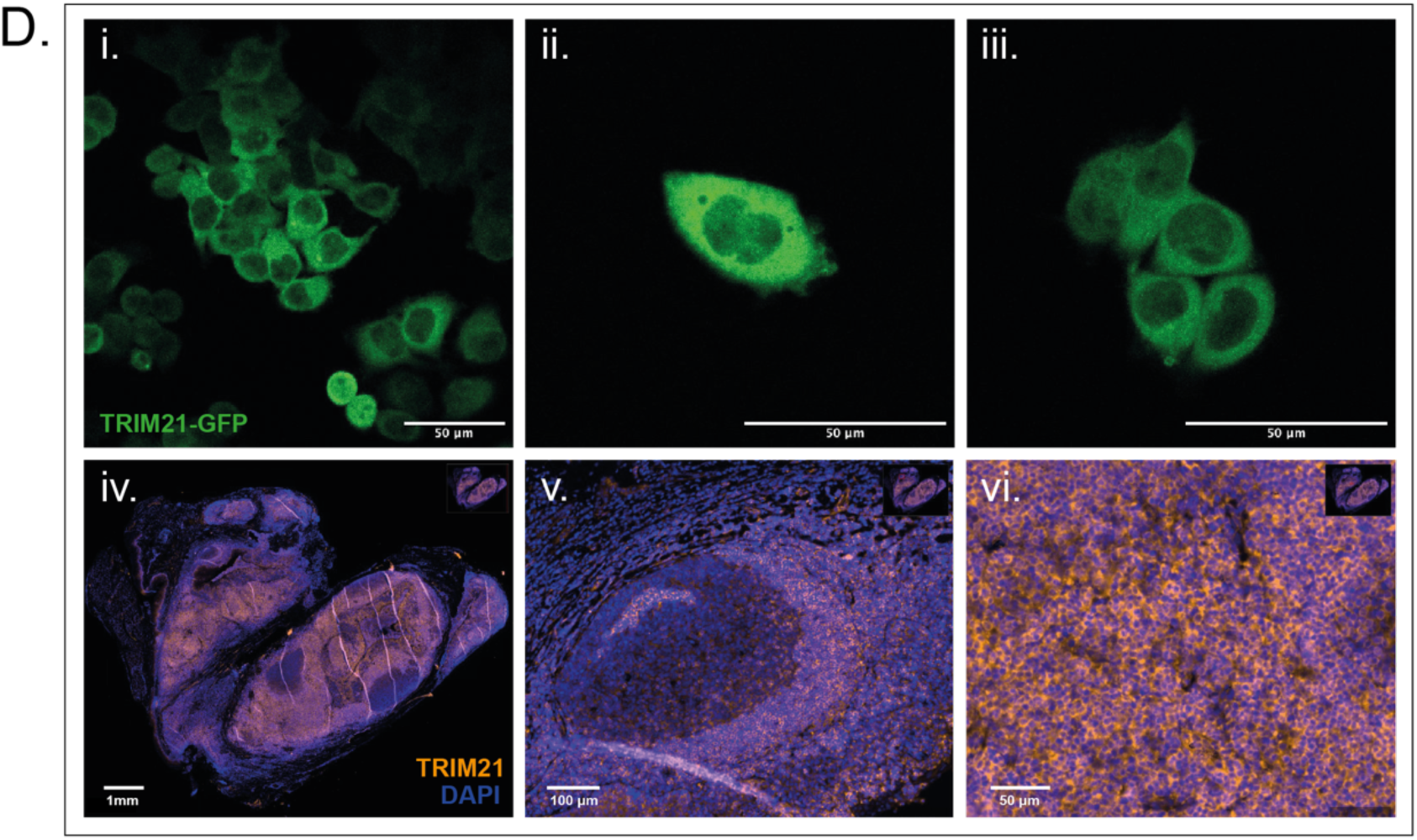
Cytosolic TRIM21 is found in SG biopsies of both control and ‘Ro’-positive SjD patients. **(A)**, **(B)**, **(C)** Serial sections of FFPE fixed SG biopsy slides were H&E stained or stained and imaged by multiplex fluorescence imaging. Representative H&E stained SG biopsy scans from sicca controls **(A) (i)** and SjD patients **(B) (i), (C) (i-vii),** were imaged on Zeiss Axioscan 7 at 20X magnification. Black and green boxes highlight areas of interest for visualisation at higher magnification. Black arrows indicate collagen deposition (fibrosis). Red arrows highlight plasma cells. **(A) (ii, iii), (B) (ii, iii), (C) (iii-vi, viii-ix)** Sequential antibody stains and bleach rounds were performed using CellDIVE^TM^ protocols (*72*). The In Cell Analyzer 2500HS (GE system) was used for image acquisition; images were merged and visualised using QuPath. Fluorescent exposures were tuned to permit visualisation of relevant markers. For example, HLA DR fluorescence was decreased to permit visualisation of IgG and IgA. Scale bars and DAPI/antibody markers are indicated in the figure. **(D) (i-iii)** Representative images of lentiviral transduced HeLa overexpressing TRIM21_w_PRYSPRY-GFP, imaged by live confocal microscopy. Scale bars, 50 µm. **(D) (iv-vi)** Representative images of tonsil slide stained with DAPI and anti-TRIM21/SS-A1 antibody, imaged using In Cell Analyzer 2500HS. **(iv)** Full tonsil section image scan. Scale bar, 1 mm. **(v, vi)** Tonsillar follicle region shown at different magnifications. Scale bars, 100 µm and 50 µm, respectively.

Another representative SjD biopsy showed regions of fibrosis, surrounding areas of acinar atrophy (Fig. 1C, i, ii, iii). TRIM21 was ubiquitously expressed (Fig. 1C, iv). Furthermore, the SjD biopsy showed infiltration by multiple immune cell types including: plasma cells (H&E stain, morphology, IgG, IgA), T cells (CD3), macrophages (CD163, CD68, MerTK), and granulocytes (CD66b) in regions of HLA-DR and immunoglobulin staining (Fig. 1C, v-ix). Together these data provided evidence for localised immune cell infiltration, inflammation and atrophy. Owing to a lack of satisfactory staining antibodies, we were unable to observe cell death in situ. However, cell death commonly occurs before immune cell infiltration and inflammation (*33*). Additionally, the destruction of SGs is a feature of SjD (*34*), and previous studies have provided evidence of epithelial cell death (*35, 36*). Thus, we hypothesised that cell death may contribute to the atrophy observed in the SjD biopsies.

We showed that TRIM21 is ubiquitously expressed in both sicca control (Fig. 1A) and SjD patient tissues (Fig. 1C). Similarly, TRIM21 was also widely expressed throughout human tonsillar tissue (Fig. 1D, iv, v, vi). To investigate TRIM21’s subcellular localisation, we stably overexpressed TRIM21-monomeric EGFP (TRIM21-mEGFP) in HeLa cells. We confirmed that TRIM21 is cytosolic in HeLa cells overexpressing TRIM21-mEGFP (Fig. 1D, i, ii, iii). Mock-transfected HeLa were negative for GFP, and a nuclear export sequence (NES)-GFP served as a control (Fig. S1A). This agrees with previously proposed subcellular localisation (*37*) and ubiquitous expression of TRIM21 in multiple cell types and tissues (*38–40*).

### TRIM21 is upregulated by inflammatory signals and released during lytic cell death

TRIM21 is upregulated by IFN signalling and proposed to regulate IFN homeostasis by participating in negative feedback loops, as part of its role in intracellular pathogen degradation (*39, 41–43*). Therefore, we expanded the variety of inflammatory stimuli known to induce IFN secretion, to explore changes in TRIM21 expression. We validated a TRIM21 antibody, suitable for immunoblotting in both humans and mice, using TRIM21 overexpression and siRNA knockdown techniques (Fig. S1B, C). TRIM21 protein was detectable in untreated (UT) mouse wild-type (WT) bone-marrow-derived macrophages (BMDMs) and human epithelial HeLa cells (Fig. 2A, B, C). TRIM21 was modestly upregulated in BMDMs in response to the Toll-like receptor (TLR) 4 agonist, lipopolysaccharide (LPS) after 8 and 24 hours of stimulation (Fig. 2A), poly I:C (TLR3 agonist) (Fig. 2A), and IFN (α, β, γ) stimulation (Fig. 2B). TRIM21 was also modestly upregulated in HeLa after 24 hours of stimulation with IFNγ (Fig. 2C). Pro-IL-1β induction by LPS was used as a positive control for LPS stimulation (Fig. 2A). These data agree with our findings that TRIM21 is constitutively expressed in many cell types (Fig. 1) and reports by others that TRIM21 can be upregulated in response to IFNs and infections (*17, 44*).

**Fig. 2.**
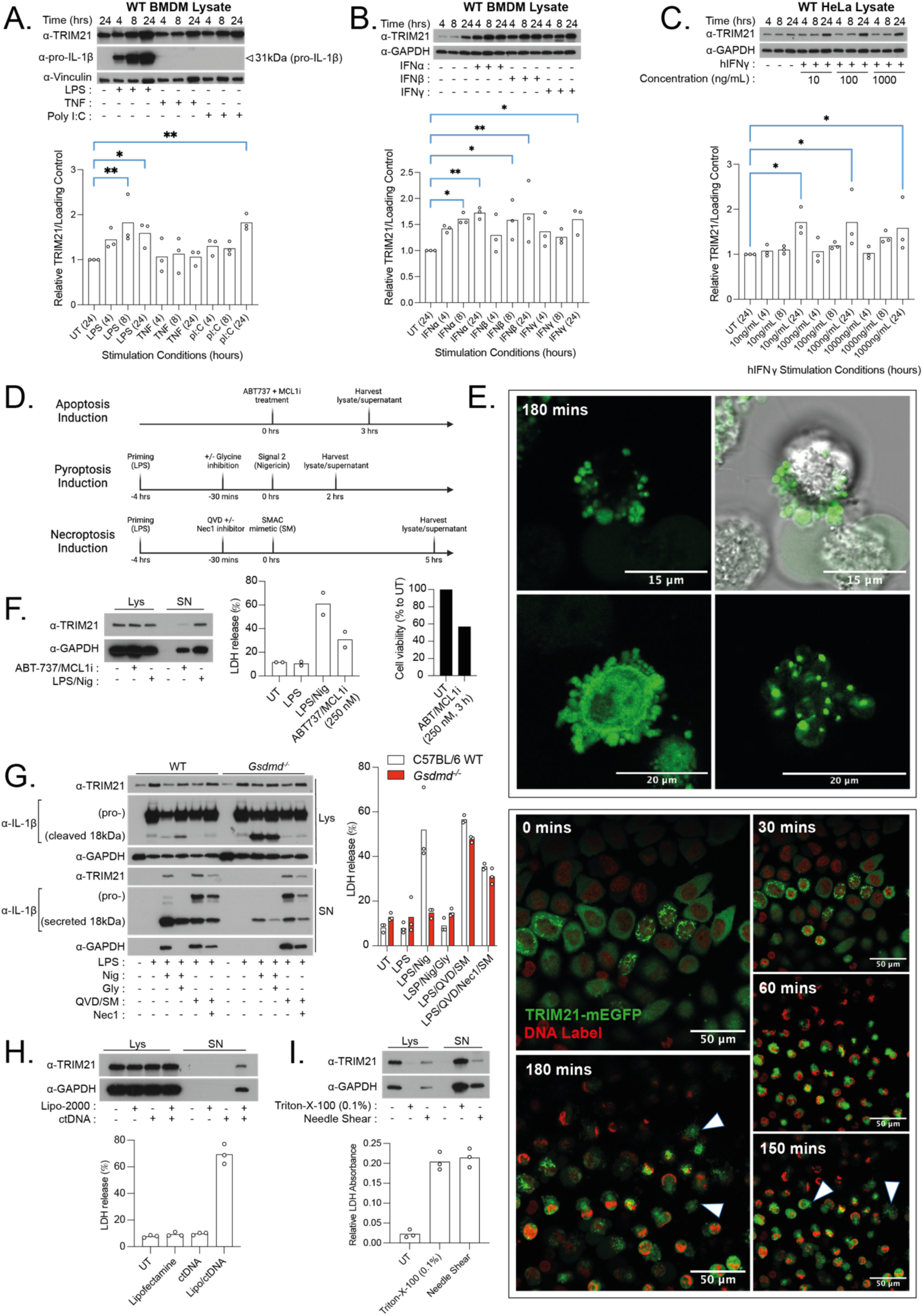
TRIM21 is upregulated by inflammatory signals and released during lytic cell death. BMDMs were stimulated with the following: **(A)** ultra-pure LPS (100 ng/mL), mTNFα (100 ng/mL), poly(I:C) (2.5 µg/mL), **(B)** mIFNα (100 ng/mL), mIFNβ (100 ng/mL), mIFNγ (100 ng/mL) for 4, 8 and 24 hours. **(C)** WT HeLa were stimulated with hIFNγ at 10 ng/mL, 100 ng/mL, or 1 μg/mL, for 4, 8 and 24 hours. **(A)**, **(B)**, **(C)** At indicated timepoints, SN were removed and cells lysed. One representative immunoblot image is shown (top). Semi-quantification of TRIM21 relative to loading control for each stimulation condition was calculated (bottom, n = 3 independent experiments). TRIM21 protein level for each stimulation condition timepoint was compared to UT 24 hour timepoint by one-way ANOVA and Fisher’s least significant difference post-hoc test. Normalised semi-quantification data are represented as single points for each of 3 independent experiments. * P ≤ 0.05, ** P ≤ 0.01, *** P ≤ 0.001. **(D)** Schematic showing cell death induction experiment designs for apoptosis, NLRP3-dependent pyroptosis and necroptosis, highlighting treatments and collection timings. **(E)** HeLa overexpressing TRIM21_w_PRYSPRY-GFP were treated with apoptosis-inducing drugs (1 μM ABT737, 10 μM MCL1i) and imaged by live confocal microscopy. Upper panels show representative images of TRIM21_w_PRYSPRY-GFP within apoptotic blebs. Scale bars, 15 µm and 20 µm. Lower panels show HeLa overexpressing TRIM21_w_PRYSPRY-GFP which were treated with 1 μM of sirDNA dye 30 mins before imaging. Apoptosis drugs were added and imaged by live confocal microscopy time-lapse recording of the same field of view. Images were selected at given intervals. White arrows highlight apoptotic blebs. Scale bars, 50 µm. Upper and lower panel images representative of 3 independent live time-lapse microscopy experiments. **(F)** WT murine BMDMs were treated with combined apoptosis-inducing drugs (250 nM ABT-737, 250 nM MCL1i) and incubated for 3 hours. For NLRP3 pyroptosis induction (LPS/Nig), BMDMs were primed with LPS (100 ng/mL) for 4 hours, then treated with Nigericin (10 μM) for an additional 2 hours. Lys and SN were collected and analysed by immunoblot. Cell death was measured by measured by LDH release assay and viability was determined by cell viability assay. Immunoblot, LDH and cell viability assays are representative results for 3 independent experiments. **(G)** WT and *Gsdmd-/-* BMDMs were primed with LPS and treated with pyroptosis/necroptosis-inducing drugs. Cells were primed with ultra-pure LPS (100 ng/mL) for 4 hours. Sterile-filtered glycine (20 mM) was added 30 minutes before Nigericin treatment for glycine inhibition of pyroptosis. To induce necroptosis, cells were treated with Q-VD-Oph (10 μM) in the last 30 mins of LPS priming, before stimulation using SMAC mimetic AZD 5582 (1 μM) for 5 hours. Nec1 (10 μM) was added at the Q-VD-Oph time-point to inhibit necroptosis. Lys and SN were collected and analysed by immunoblot. Cell death was measured by LDH release assay. Data are representative of 3 independent experiments. **(H)** WT BMDMs were treated with 200 ng ctDNA (2 µg/ml) and 0.25 % Lipofectamine-2000 by spinfection to induce AIM2-dependent pyroptosis. Lys and SN were collected and analysed by immunoblot. Cell death was measured by LDH release assay. Data are representative of 3 independent experiments. **(I)** WT BMDMs were treated with 0.1% Triton X-100 or needle sheared to induce necrosis. Lys and SN were collected and analysed by immunoblot. Relative LDH absorbance was measured. Data are representative of 3 independent experiments.

We next asked how TRIM21 is released from cells. Recently, pro-inflammatory forms of lytic cell death such as pyroptosis have been proposed to induce autoimmunity by promoting antigen and inflammatory alarmin release (*45*). Thus, we sought to compare apoptosis with lytic cell death as possible mechanisms for intracellular TRIM21 release. The following forms of programmed cell death were tested: apoptosis, pyroptosis, necroptosis, and mechanical lysis mimicking necrosis (Fig. 2D).

In HeLa cells undergoing apoptosis, apoptotic bodies (blebs) containing TRIM21 were imaged (Fig. 2E, upper four panels). Live confocal microscopy time-lapse recording of the same field of view showed cytoplasmic condensation and blebbing after 30 minutes of drug treatments (ABT-737/MCL1i) (Movie S1). Highly condensed TRIM21-mEGFP puncta were still visible 180 minutes after drug treatment, after many cells had undergone secondary necrosis (Fig. 2E, lower five panels). Membrane-bound apoptotic blebs containing nuclear and cytoplasmic material are usually targeted for clearance by neighbouring phagocytes, thus preventing the release of intracellular contents into the extracellular space (*46*).

The release of free TRIM21 into the extracellular space was next tested by immunoblotting cell lysates (Lys) and supernatants (SN) after apoptosis versus lytic cell death, pyroptosis. BMDMs were treated with ABT-737/MCL1i to induce intrinsic apoptosis, or LPS/Nigericin to induce pyroptosis and cell lysis. Release of intracellular proteins TRIM21 and GAPDH was strongly detected in SN of pyroptotic cells but less so in SN of apoptotic cells (Fig. 2F), when proteins are expected to remain trapped in apoptotic cells and blebs. We used standard in vitro techniques to confirm which specific cell death pathways were induced. Lytic death downstream of pyroptosis was confirmed by the detection of another cytosolic protein, lactate dehydrogenase (LDH) which is only released upon membrane lysis. Less LDH was detected after apoptosis, with some level of cell lysis by secondary necrosis which occurs during late apoptosis (Fig. 2F). Loss of viability downstream of apoptosis was confirmed by the CellTiter-Glo Luminescent Cell Viability Assay (Fig. 2F).

To further explore the release of TRIM21 during pro-inflammatory lytic cell death (pyroptosis/necroptosis), we utilised BMDMs from WT and Gasdermin D KO (*Gsdmd^-/-^*) mice (Fig. S1D). *Gsdmd^-/-^* BMDMs are unable to form Gsdmd pores at the membrane, and have reduced lytic death response and reduced secretion of processed IL-1β (*47, 48*). As expected, WT BMDMs which had been treated with LPS/Nig to induce NLRP3-dependent pyroptosis, upregulated pro-IL-1β in the cell (as confirmed by cell lysate immunoblot), underwent lytic cell death (as confirmed by LDH release), and enabled the secretion of IL-1β, and release of TRIM21 and GAPDH in the culture SN (as confirmed by cell SN immunoblot) (Fig. 2G). TRIM21 and GAPDH were not present in the SN of *Gsdmd-/-* BMDM which had been stimulated with LPS/Nig, indicating that blocking cell lysis inhibited TRIM21 release (Fig. 2G).

As the GSDMD pore facilitates the secretion of processed IL-1β, we tested whether TRIM21 could pass through the GSDMD pore. Before signal 2 (Nig) treatment, 20 mM osmoprotectant glycine was added to WT BMDMs. This permits GSDMD pore formation and IL-1β secretion but inhibits downstream Ninjurin-1-mediated membrane lysis (*49*). IL-1β but not TRIM21 was detected in the SN of glycine-treated WT BMDMs after pyroptosis induction (Fig. 2G). This suggests TRIM21 does not pass through the GSDMD pore but is released by cell lysis. This was expected as the GSDMD pore only permits molecules of a certain size and charge (*50*); TRIM21, at 52 kDa, is likely too large and possibly the wrong charge.

To further test if TRIM21 is released from cells lysed in response to pathogenic insult, we induced pyroptosis by the AIM2-inflammasome (cytosolic dsDNA sensor) in BMDMs (*51*). AIM2-dependent lytic cell death was induced by lipofectamine spinfection of calf-thymus DNA (ctDNA) into BMDMs, releasing TRIM21 and LDH into cell SN (Fig. 2H).

Necroptosis is another programmed form of pro-inflammatory lytic cell death, often induced by pathogenic insult (e.g. virus) (*52*). Treatment of either WT or *Gsdmd^-/-^* BMDMs with necroptosis-inducing drugs (LPS/QVD/SMAC mimetic (SM)) (Fig. 2D), was sufficient to induce cell lysis, releasing TRIM21 into SN for detection (Fig. 1G). As expected, this extracellular TRIM21 release was reduced by an inhibitor of necroptosis, Necrostatin (Nec)-1 (*53*) treatment (Fig. 2G).

Finally, necrotic cell death of BMDMs by either needle shear (to mimic mechanical lysis), or by Triton-X-100 (0.1%), almost completely depleted TRIM21 and GAPDH from the cell lysates and released them into the cell SN (Fig. 2I).

Collectively, these data indicate that cytosolic TRIM21 is widely expressed, and upregulated following pathogenic (LPS/Poly I:C) or IFN stimulation. TRIM21 gets released from cells during pro-inflammatory forms of lytic cell death (pyroptosis, necroptosis, necrosis), following membrane rupture, but not during the normal physiological process of apoptosis when it remains trapped in membrane blebs.

### Once released, extracellular TRIM21 binds to the Fc domain of serum antibodies

Inside the cytosol, TRIM21 binds the Fc domain of immunoglobulins via its PRYSPRY domain (*54*). We aimed to test whether TRIM21 maintains its antibody-binding abilities once released outside the cell, for example after lytic cell death. To mimic natural TRIM21 release, we overexpressed three GFP-tagged TRIM21 constructs in HeLa cells (Fig. 3A). WT TRIM21_w_PRYSPRY is a full-length construct, whilst TRIM21_NO_PRYSPRY is a truncated protein construct lacking the PRYSPRY domain. Thirdly, TRIM21_FcMUT is full-length and contains two PRYSPRY domain point mutations (W381A/W383A) previously shown to ablate Fc binding (*55*) (Fig. 3A).

**Fig. 3.**
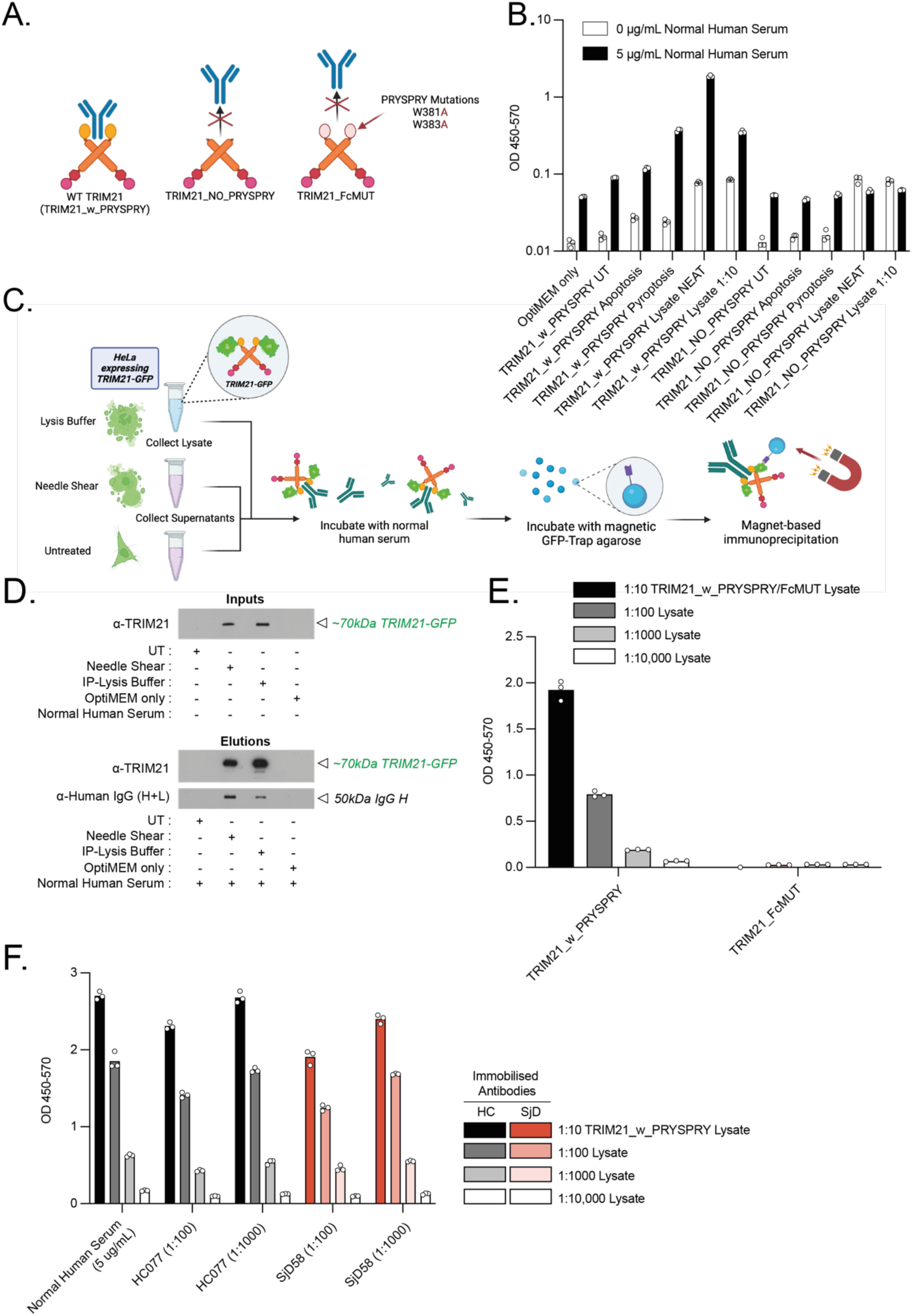
Once released, extracellular TRIM21 binds to the Fc domain of serum antibodies. **(A)** Schematic showing architectures and binding abilities of TRIM21 overexpression constructs; WT TRIM21 (w_PRYSPRY), TRIM21 without PRYPSRY (NO_PRYSPRY) and TRIM21 with PRYSPRY mutations (FcMUT). TRIM21_NO_PRYSPRY and TRIM21_FcMUT constructs are unable to bind Fc portion of antibodies. **(B)** GFP ELISA to detect TRIM21-GFP from HeLa after different cell deaths. HeLa overexpressing TRIM21_w_PRYSPRY-mEGFP or TRIM21_NO_PRYSRY-mEGFP were used for experimentation. To induce apoptosis, HeLa were treated with ABT-737 (1 μM) and S63845/MCL1i (10 μM) for 2 hours. For non-canonical pyroptosis induction, HeLa were primed for 16 hours with 10 ng/mL hIFNγ and spinfected with final concentration per well of 1 μg LPS and 0.25% Lipofectamine-2000. Cells were incubated for 24 hours to ensure complete pyroptosis. SN were collected from UT, apoptotic and pyroptotic HeLa. Lysates were collected from HeLa using non-denaturing IP lysis buffer. ELISA plates were coated with normal human serum (5 μg/mL) or PBS for background control. Samples were detected by anti-GFP-HRP antibody. Data shown as all triplicate points for single experiment. ELISA is representative of 3 independent experiments. **(C)** Schematic outlining GFP-trap experiment methodology. HeLa overexpressing TRIM21_w_PRYSPRY were lysed, needle sheared or UT. Lys or SN were collected and incubated with 0.01 mg/mL normal human serum and incubated for 2 hours under rotation at 4°C. Pre-equilibrated magnetic GFP-trap beads were added and incubated for 1 hour under rotation at 4°C. Beads were washed and material eluted by magnet-based IP. **(D)** GFP-trap inputs and elutions were analysed by immunoblot. Immunoblot results are representative of 3 independent experiments. **(E)** GFP ELISA to detect antibody binding of TRIM21 constructs. HeLa overexpressing TRIM21_w_PRYSPRY-GFP or TRIM21_FcMUT-GFP were lysed in non-denaturing IP lysis buffer. Lysates were diluted in blocking buffer and analysed by ELISA on plates coated with normal human serum (5 μg/mL). Data shown as all triplicate points for single experiment. ELISA is representative of 3 independent experiments. **(F)** GFP ELISA to detect binding affinity of diluted TRIM21_w_PRYSPRY lysates to the following: normal human serum (5 μg/mL), HC077 plasma and SjD58 patient plasma. Patient plasmas were diluted 1:100 and 1:1000, as indicated. Data shown as all triplicate points for single experiment. ELISA is representative of 3 independent experiments.

We developed a custom GFP-sandwich ELISA to detect TRIM21 binding to normal human serum antibodies pre-bound to the ELISA plates. All treatments were performed in serum-free OptiMEM media to prevent confounding results with antibodies present in FBS. In agreement with our previous results showing TRIM21 trapped inside apoptotic blebs (Fig. 2F), minimal binding was observed between SN from UT or apoptotic HeLa_TRIM21_w_PRYSPRY cells, to plate-immobilised antibodies (Fig. 3B). As HeLa lack canonical NLRP3 inflammasomes, we induced non-canonical pyroptosis by LPS-spinfection (*56*), to trigger GSDMD-mediated lytic death. TRIM21 released from pyroptotic HeLa_TRIM21_w_PRYSPRY cells bound strongly to plate-bound antibodies, at levels similar to TRIM21 lysate cells. Binding interactions were PRYSPRY-dependent, and lost in HeLa_TRIM21_NO_PRYSPRY cells (Fig. 3B).

To test whether extracellular TRIM21 can also bind to circulating (not plate-bound) antibodies, we performed co-immunoprecipitation (co-IP) whereby TRIM21 is free to bind antibodies in three-dimensional space (Fig. 3C). We mechanically lysed cells (needle shear or immunoprecipitation (IP) lysis buffer) to release TRIM21 and incubated SN with normal human serum. We then used GFP-trap IP with anti-GFP nanobody to pull down GFP-tagged TRIM21 and any serum antibodies bound to it (Fig. 3C). TRIM21 was only present in needle sheared cell SN and IP buffer-lysed HeLa_TRIM21_w_PRYSPRY inputs (Fig. 3D). After incubation with normal human serum and subsequent GFP-trap IP, only inputs containing TRIM21 were able to co-immunoprecipitate antibodies detectable by immunoblotting with anti-IgG heavy-chain (Fig. 3D), suggesting that TRIM21, when released from cells, retains its ability to bind antibodies in solution.

To confirm that TRIM21-antibody binding was PRYSPRY-dependent, we performed a GFP-ELISA. Lysates from HeLa overexpressing either TRIM21_w_PRYSPRY-GFP or TRIM21_FcMUT-GFP were diluted and analysed for binding plate-bound antibodies from normal human serum. Plates were washed, and bound TRIM21 detected with anti-GFP. Only WT TRIM21_w_PRYSPRY-GFP but not TRIM21_FcMUT-GFP retained antibody binding capacity (Fig. 3E), suggesting that TRIM21 released from cells retains its ability to bind antibodies, and that binding is Fc-dependent.

Finally, we tested whether TRIM21-Fc binding differs between healthy control (HC) versus SjD patient serum samples. We coated ELISA plates with normal human serum, HC plasma, or SjD patient plasma. Regardless of TRIM21_w_PRYSPRY lysate dilution, TRIM21-Fc binding was similar for all samples (Fig. 3F), suggesting that in two-dimensional plate-bound setting, when binding is mediated via Fc, TRIM21 can bind both healthy and SjD antibodies.

These results confirm that TRIM21 binds the Fc portion of antibodies via its PRYSPRY domain, consistent with previous literature (*54*). This binding activity is maintained even after lytic cell death, and TRIM21’s Fc binding affinity does not differ between HC and SjD plasma.

### Plasma from SjD patients creates higher-order TRIM21-IgG complexes

According to the 2016 ACR-EULAR criteria for diagnosis, patients with primary SjD must present with autoantibodies against the Ro/SSA antigens (*11*). However, standard laboratory autoantibody tests often do not distinguish between the specific “Ro” antigens (Ro52/TRIM21 or Ro60/TROVE2) (*57*), despite evidence showing clear diagnostic utility in separate detection (*58*). Therefore, we employed the specific luciferase immunoprecipitation system (LIPS) assay for accurate autoantibody detection against the separate classical SjD autoantigens (Ro52/TRIM21_NO PRYSPRY, Ro60/TROVE2, and La) (*59*) (Fig. 4A). We identified three HC plasmas (HC099, HC077, HC087) without SjD autoantigen reactivity (Fig. 4B). Analysis of SjD plasma confirmed four samples with anti-TRIM21/Ro52 autoreactivity, one of which also had anti-Ro60/TROVE2 autoreactivity (Fig. 4B). We selected three SjD patient plasmas (SjD58, SjD59, SjD61) with a range of autoantibody titres as biological replicates for subsequent experiments.

**Fig. 4.**
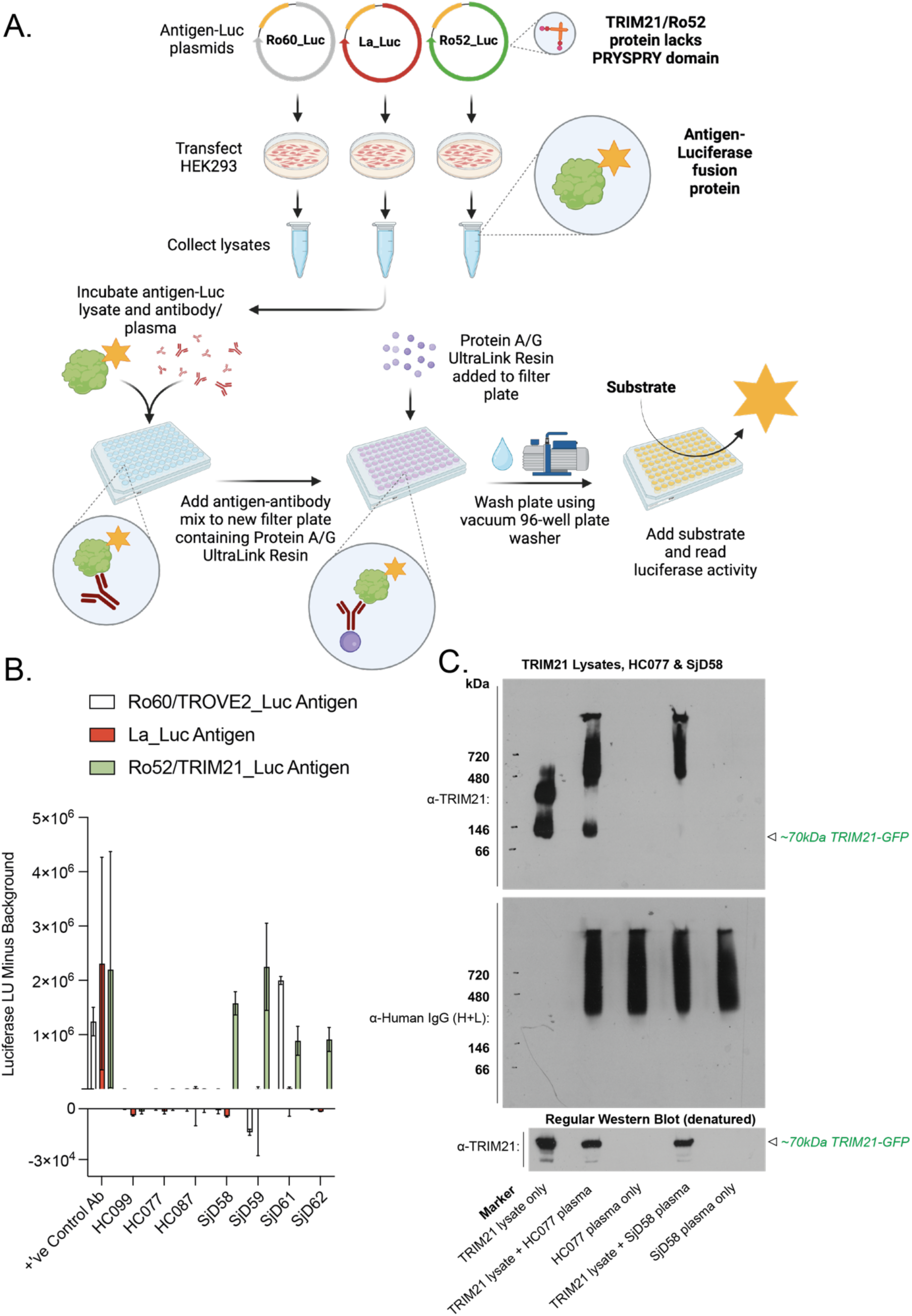
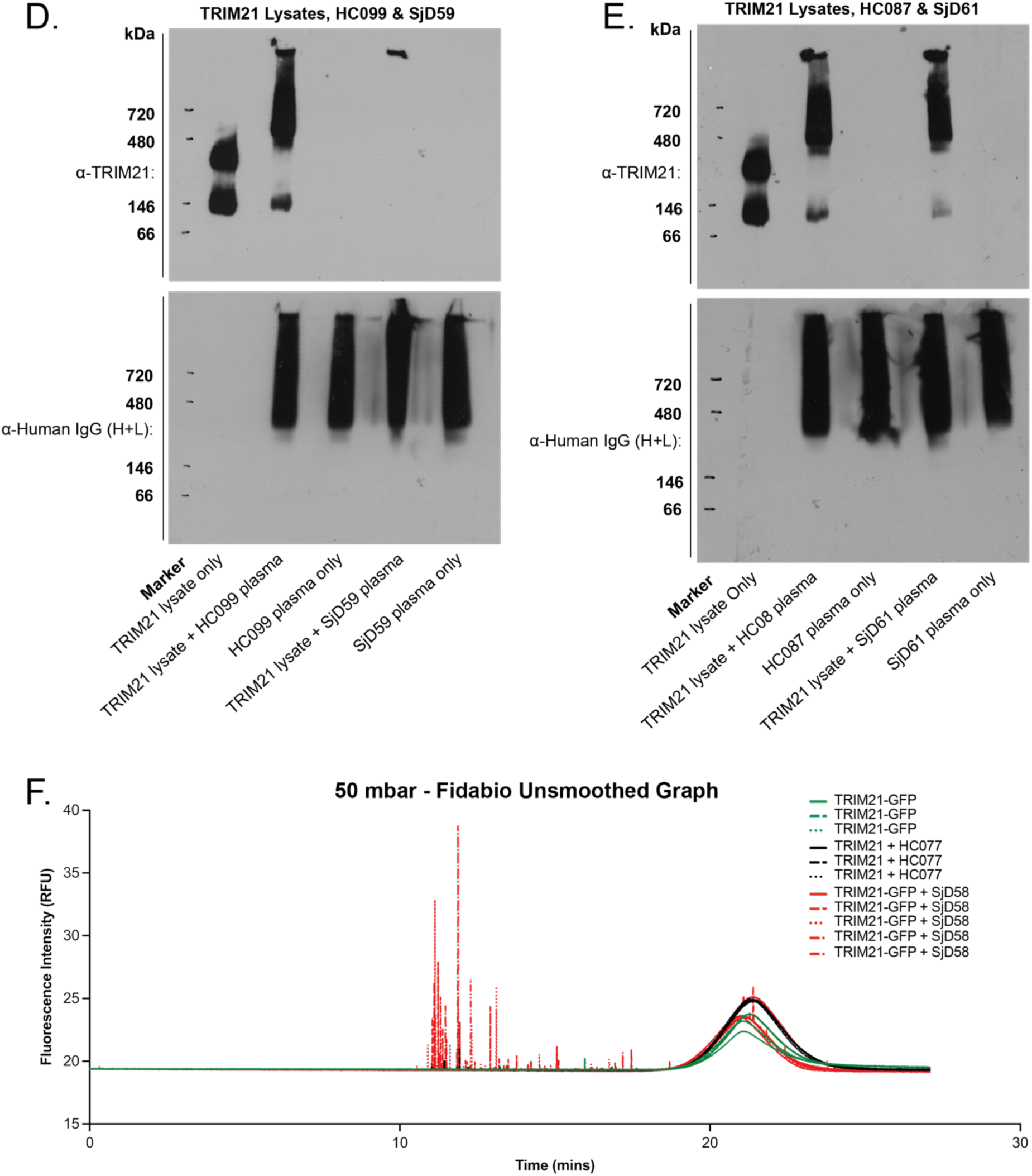
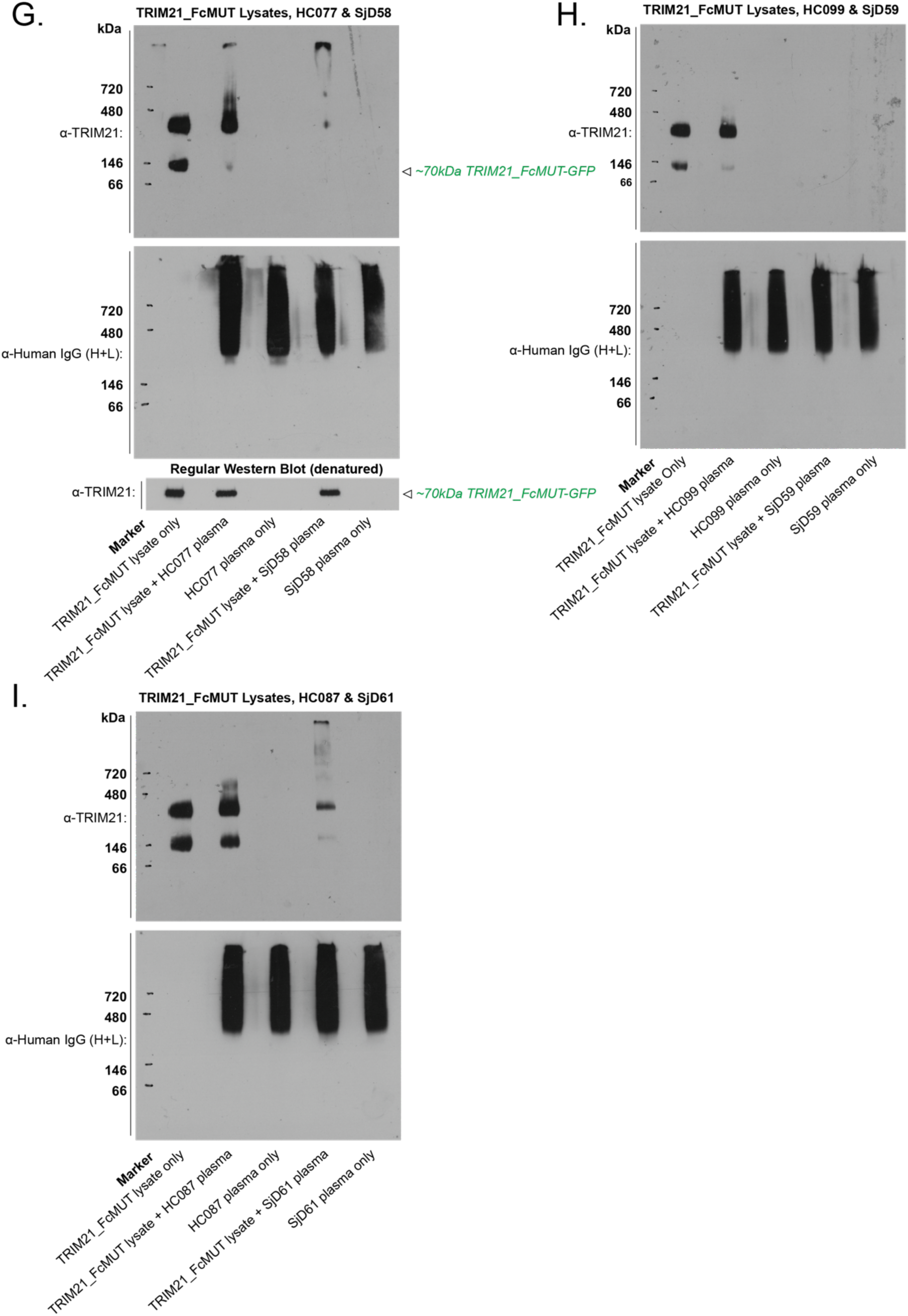
Plasma from SjD patients generates higher-order TRIM21-IgG complexes because the interaction is mediated both via Fc and Ag-specific Fab domains. **(A)** Schematic illustrating the LIPS assay to quantify antigen-specific antibodies from patient plasma (*106*). Autoreactivity against the three classical Sjögren’s autoantigens (TROVE2/Ro60, La, TRIM21/Ro52) was measured by luciferase activity. **(B)** Plasma samples (HC/SjD) were diluted 1:10 for analysis. Positive control antibodies were diluted to final 0.01 mg/mL concentration. Samples were incubated with one of the following classical SjD Luc-antigen lysates, as indicated: Ro60/TROVE2_Luc, La_Luc, Ro52/TRIM21_Luc. Luciferase activity was measured, subtracting background Luc_antigen-only luciferase activity to obtain final LU. Representative data are shown as a mean with SD of triplicates from one of 3 independent experiments. Non-denaturing BN-PAGE separation of **(C**, **D**, **E)** TRIM21_w_PRYSPRY-GFP lysates or **(G**, **H**, **I)** TRIM21_Fc_MUT-GFP lysates from HeLa, TRIM21 lysates complexed with plasmas (HC/SjD), or plasma alone. Molecular weight markers are indicated. **(C**, **G [bottom])** Regular denaturing immunoblotting of samples identified TRIM21-GFP proteins at expected molecular weights. All biological replicate data are shown to represent range of complex sizes formed as a result of different amounts of autoantibodies present in plasma samples. **(F)** TRIM21_w_PRYSPRY-GFP lysates alone or complexed with patient plasma (HC077 or SjD58) were analysed by FIDA. All unsmoothed fluorescence intensity data obtained at 50 mbar represented for single experiment with technical triplicates.

We previously showed that TRIM21 binds Fc domains of plate bound HC and SjD antibodies in a PRYSPRY domain-dependent manner (Fig. 3F). However, it remained unclear whether cell-free TRIM21 could form higher-order ICs with SjD plasma autoantibodies, which would have additional F(ab’)_2_-mediated TRIM21 binding capacity. To test this, TRIM21 was released by lysis from HeLa_TRIM21_w_PRYSPRY-GFP cells and incubated with HC or SjD plasma before analysis using non-denaturing Blue-Native Page (BN-PAGE) to detect higher-order complexes (*60*). In lysates, TRIM21-GFP was identified at both monomeric (∼70kDa) and multimeric form (Fig. 4C, D, E, lysate only lane), as expected from some level of intrinsic PRYSPRY protein self-oligomerisation. When incubated with HC plasma, higher-order TRIM21-antibody complexes were formed (Fig. 4C, D, E, HC lanes). These complexes were specific to conditions containing purified TRIM21-GFP and not seen in lanes containing HC plasma only.

The complexes between TRIM21_w_PRYPSRY and SjD plasmas were even larger than those formed with HC plasma, and correlated with the range of anti-TRIM21/Ro52 autoantibody titres (Fig. 4B, C, D, SjD plasma lanes). TRIM21 complexed with SjD58 presented higher-order TRIM21-autoantibody complexes with sizes greater than 480kDa (Fig. 4E). For SjD59, with the highest anti-TRIM21-autoantibody titres, the complexes formed were too large to be adequately resolved by BN-PAGE methods, and barely entered the gel (Fig. 4B, D). SjD61 formed the smallest IC sizes, with some remaining monomeric TRIM21, consistent with lower autoantibody titres (Fig. 4B, E). Total IgG levels were similar in all HD and SjD plasma samples (Fig. 4C, D, E, G, H, I, α-Human IgG detection). Together, these results indicated the size of TRIM21-IgG IC was affected by the levels of TRIM21 autoantibodies detected within the SjD plasma.

To more accurately quantify complex sizes, we utilised flow-induced dispersion analysis (FIDA) technology. This enabled us to track the GFP-tag on TRIM21, using very small sample volumes, and measure Taylor dispersion to characterise complex sizes. Samples containing TRIM21-GFP alone did not demonstrate conditions outside Taylor conditions, indicative of small complex sizes. Results for TRIM21-GFP complexed with HC077 indicated a hydrodynamic radius size range of 17-53 nm, whereas when complexed with SjD58 plasma these were 17-212 nm. Some of these very large complexes formed insoluble aggregates, presenting as “spikes” when graphically represented (Fig. 4F).

To determine whether PRYSPRY-mediated Fc binding was necessary and sufficient for production of large ICs, we performed BN-PAGE analyses using TRIM21_FcMUT-GFP lysates. Higher-order complex formation between TRIM21_FcMUT and HC plasma was impaired, with complex size matching TRIM21_FcMUT lysates alone (Fig. 4G, H, I, HC plasma lanes), suggesting that TRIM21 binding to healthy serum antibodies is dependent on PRYSPRY-mediated Fc binding. However, incubation with SjD plasma yielded surprising results, with higher-order complex formation retained and even enhanced with SjD58 and SjD61 when Fc-binding was eliminated (Fig. 4G, I, SjD plasma lanes). Incubation with SjD59 led to higher-order complex formation (Fig. S2A, B), which again could not be adequately resolved by BN-PAGE methods (Fig. 4H). This suggested that antibodies in SjD patient plasma can still form higher-order TRIM21-ICs even in the absence of PRYSPRY-mediated Fc binding, as binding is likely mediated by F(ab’)_2_. Collectively, these results indicated that in SjD patients both Fc and F(ab’)_2_ regions contribute to high-order TRIM21-antibody complex formation.

### Higher-order TRIM21-IgG complexes are taken up by macrophages

Macrophages capture IgG-opsonised materials through binding to Fcγ receptors (FcγRs) (*61–63*). Due to limited protein and patient plasma availability, we established a micro-scale assay for TRIM21-antibody complex uptake. TRIM21 proteins were first micro-purified by GFP-trap (Fig. 3C), then combined with Opti-MEM control or serum/plasma antibodies before feeding to IFN-stimulated macrophages for 5-60 min. IFN priming was used to upregulate the expression of FcγRs and mimic the inflammatory state seen in SjD. We first compared the uptake of TRIM21-GFP alone versus that of TRIM21-GFP-IC when ICs were prepared with normal human serum (NHS). After washing, cells were lysed and uptake analysed by immunoblot using anti-TRIM21 antibodies to distinguish exogenously added TRIM21-GFP (∼70 kDa) from endogenous TRIM21 (∼52 kDa). TRIM21-GFP-IC were more readily taken up by BMDMs at all 37°C timepoints, compared to TRIM21-GFP alone. Uptake was reduced in cells kept on ice (permitting surface binding but not internalisation) (Fig. 5A).

**Fig. 5.**
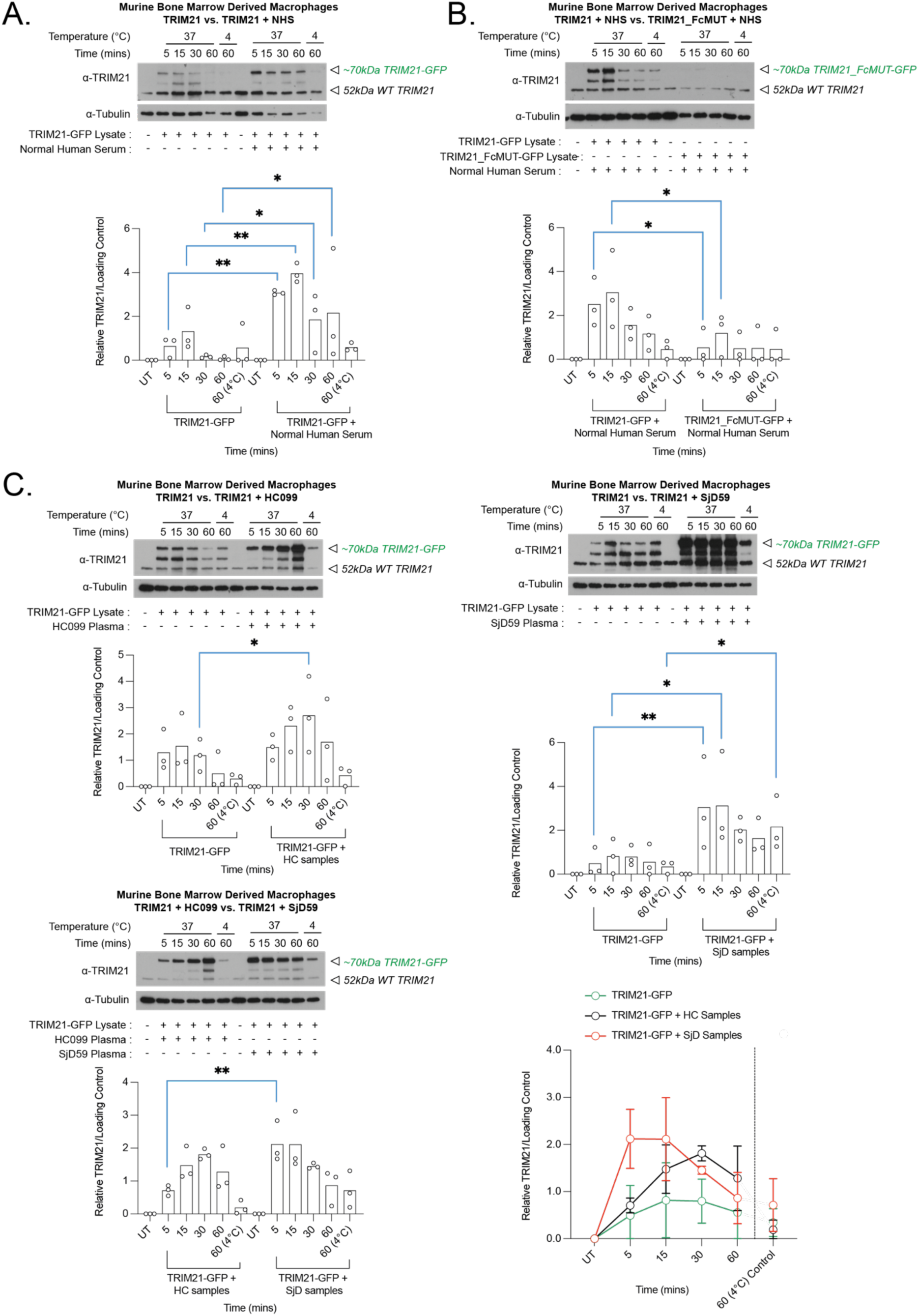
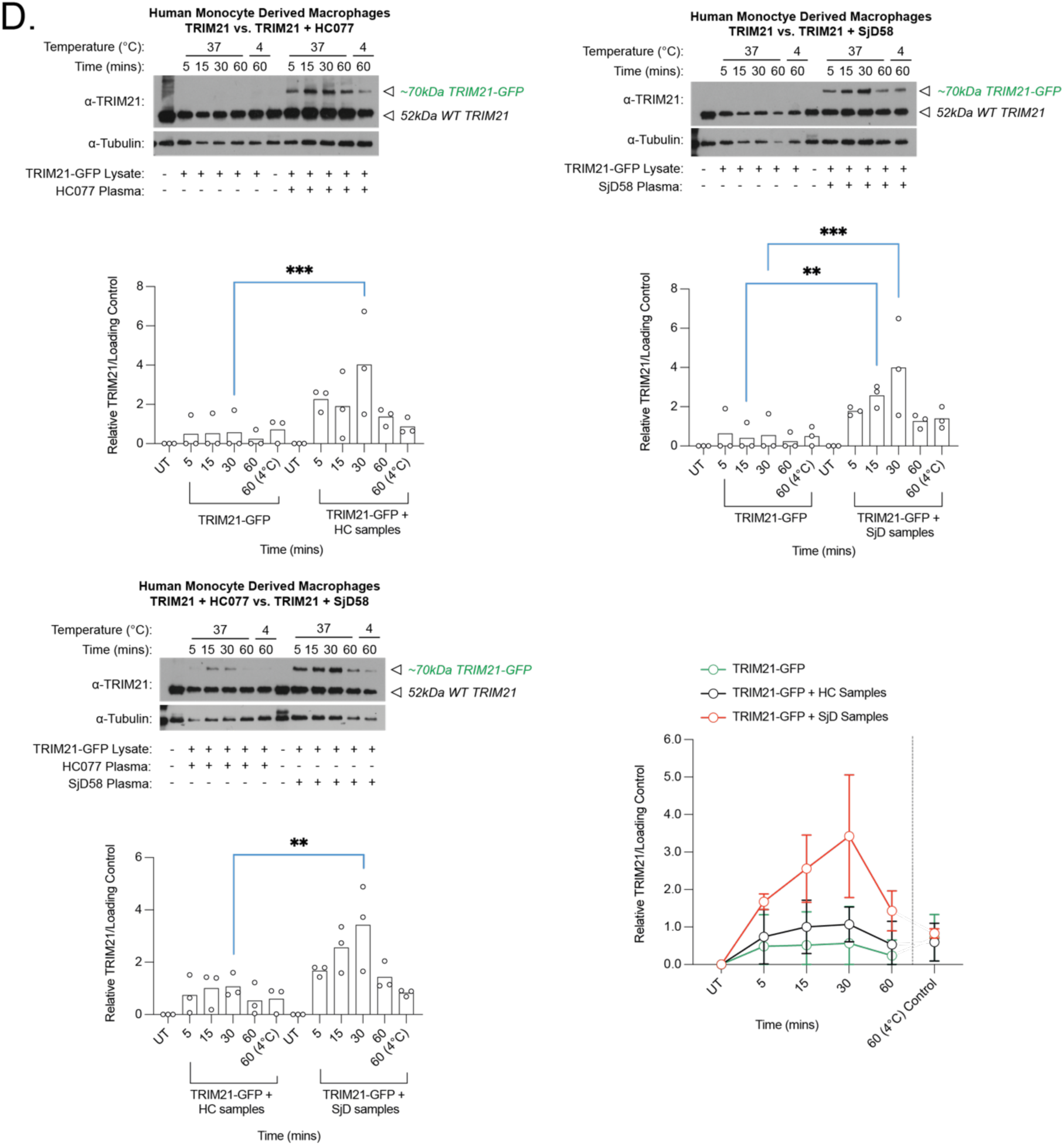
Higher-order TRIM21-IgG complexes are taken up by macrophages. BMDMs/HMDMs were stimulated with 100 ng/mL murine/human IFNγ for ∼24 hours prior to feeding with the following GFP-trap eluted proteins; **(A)** TRIM21_w_PRYSPRY-GFP alone, TRIM21_w_PRYSPRY-GFP complexed with normal human serum (0.01 mg/mL), **(B)** TRIM21_FcMUT-GFP alone, TRIM21_FcMUT-GFP complexed with normal human serum (0.01 mg/mL), **(C**, **D)** TRIM21_w_PRYSPRY was complexed with plasma (HC/SjD) (1:100 dilution). Uptake reactions were stopped after 5, 15, 30 and 60 mins. Cells kept on ice (4°C) provided control background binding levels. Lysates were collected and analysed by immunoblot. One representative immunoblot image is shown. Semi-quantification of TRIM21-GFP relative to loading control for each timepoint was calculated. **(C)** Immunoblots are representative of three independent biological replicates, using the following patient pairs: HC077 vs. SjD58, HC099 vs. SjD59, HC087 vs. SjD61. Kinetic graph shows combined relative immunoblot TRIM21-GFP uptake data from all three experiments **(A**, **B**, **C)** TRIM21-GFP protein level for each paired timepoint was compared by one-way ANOVA and Fisher’s least significant difference post-hoc test. Normalised semi-quantification data are represented as single points for all 3 independent experiments. * P ≤ 0.05, ** P ≤ 0.01, *** P ≤ 0.001. **(D)** HMDM uptake immunoblot is representative of three independent biological replicates, using the following subject pairs; HC077 vs. SjD58, HC099 vs. SjD59, HC087 vs. SjD61. Kinetic graph shows combined relative immunoblot TRIM21-GFP uptake data (all samples, n = 3 HC and 3 SjD).

TRIM21_FcMUT lysates were then used to confirm that uptake of TRIM21 ICs prepared with NHS was dependent on direct TRIM21-antibody binding via the PRYPSRY domain, rather than nonspecific antibody coating of TRIM21 protein. TRIM21_FcMUT-IgG complexes were not taken up as efficiently by BMDMs compared to WT TRIM21-IgG complexes (Fig. 5B).

As shown above, TRIM21 formed higher-order ICs with SjD patient plasma (Fig. 4C, D, E). Micro-purified TRIM21 was complexed with HC or SjD patient plasma. BMDM uptake of TRIM21-ICs had faster uptake kinetics for SjD plasma-complexed-TRIM21 compared to HC-complexed-TRIM21 (Fig. 5C). We performed these uptake assays for 3 SjD patients with variable anti-TRIM21/Ro52 autoantibodies (SjD58, SjD59, SjD61) (Fig. 4B). As expected, the uptake was slightly increased and accelerated for patient SjD58 with medium autoantibody levels, compared to SjD61 with lower autoantibody titres (Fig. S3A, B). These data suggested that larger TRIM21-ICs formed with anti-TRIM21/Ro52-positive SjD plasma are more rapidly taken up by BMDMs.

Finally, we confirmed that this uptake is not limited to murine BMDM, and is recapitulated by IFN-primed human monocyte derived macrophages (HMDMs). HMDM uptake of TRIM21-ICs was faster and stronger for IC formed with SjD patient plasma compared to those formed with plasma from HCs or uptake of TRIM21 alone (Fig. 5D). Additionally, uptake was accelerated and stronger for ICs prepared from SjD58 plasma with higher autoantibody levels (Fig. 5D).

### Once taken up, TRIM21 escapes into the cytosol and is preferentially degraded by the proteasome

We consistently detected reduced TRIM21 levels 60 minutes post-uptake in the macrophage TRIM21-uptake experiments (Fig. 5). Therefore, we sought to explore whether this reduction is due to TRIM21 degradation, and if it could be rescued with lysosomal (Bafilomycin A1 (BafA1)) or proteasomal (MG132) inhibitors.

In BMDMs treated with the lysosomal acidification inhibitor, BafA1, TRIM21 complexed with HC or SjD plasma Ig, but not TRIM21 alone, was protected from degradation after uptake (Fig. 6A). However, in BMDMs treated with MG132 proteasomal inhibitor, all three conditions (TRIM21 alone, TRIM21 complexed with HC plasma, or TRIM21 complexed with SjD plasma) showed protection from degradation at 60 minutes (Fig. 6A). In fact, in MG132-treated BMDMs fed TRIM21 complexed with either HC or SjD plasma antibodies, we observed increased accumulation of TRIM21 within the cell at 60 minutes (Fig. 6A), suggesting that both proteasomes and lysosomes can degrade TRIM21 taken as part of IC, but the proteosome may be the dominant pathway for TRIM21 alone. Data also suggest that, after the uptake, TRIM21 can escape the endo-lysosome and gain access to the cytosol, for proteasomal degradation.

**Fig. 6.**
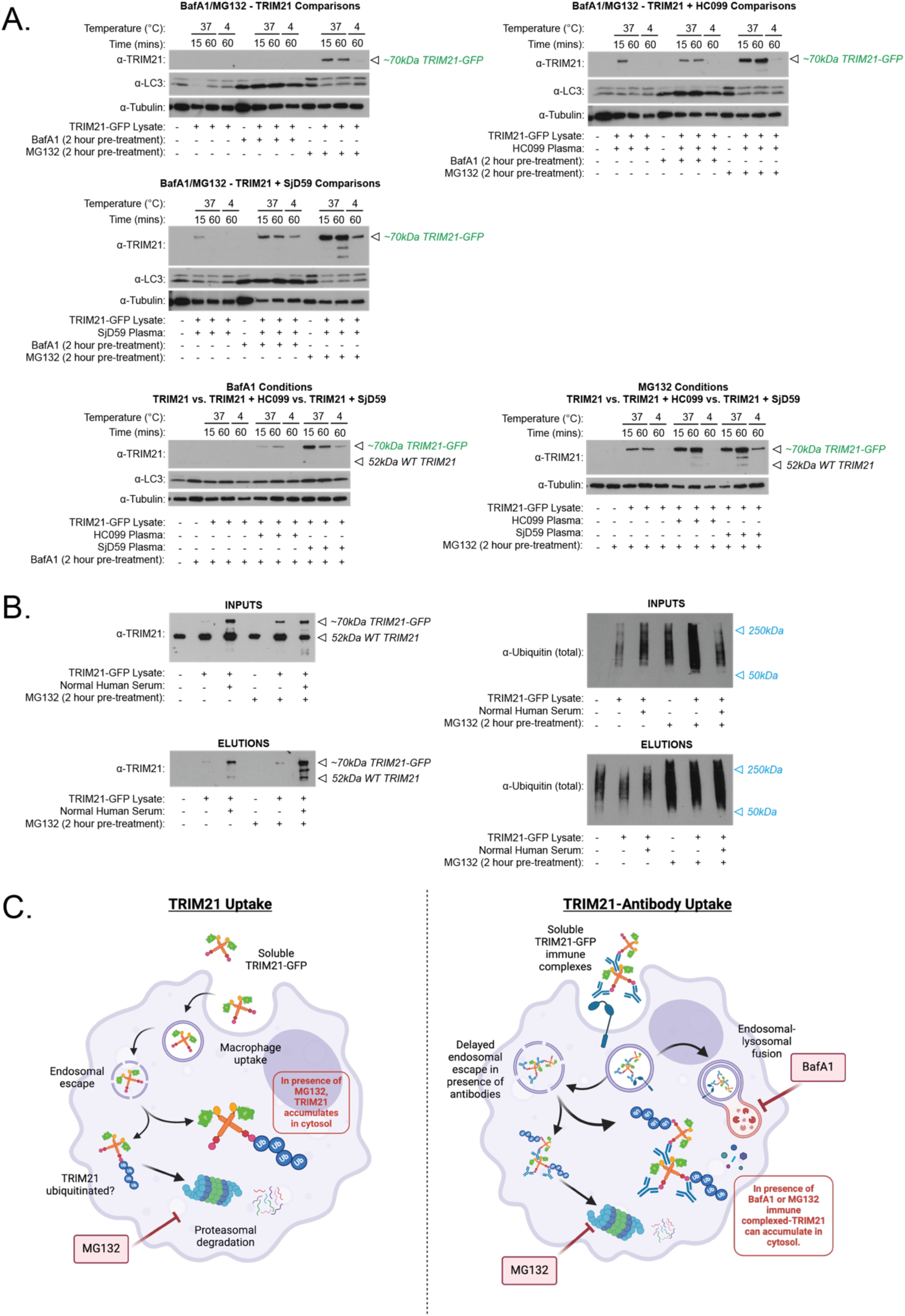
Once taken up, TRIM21 escapes into the cytosol and is preferentially degraded by the proteasome. **(A)** BMDMs were stimulated with 100 ng/mL mIFNγ for ∼24 hours. Before feeding, cells were pre-treated with either 10nM BafA1 or 10 μM MG132. Cells were fed with the following GFP-trap eluted proteins: TRIM21_w_PRYSPRY-GFP alone, TRIM21_w_PRYSPRY-GFP complexed with plasma (HC/SjD) (1:100 dilution). Uptake reactions were stopped after 15 and 60 mins. Cells kept on ice (4°C) provided control background binding levels. Lysates were collected and analysed by immunoblot, in multiple comparative gels to include the various conditions. One representative immunoblot image for each comparison is shown. Immunoblots are representative of three independent biological replicates, using the following subject pairs; HC077 vs. SjD58, HC099 vs. SjD59, HC087 vs. SjD61. **(B)** IFNγ-stimulated BMDMs were fed with TRIM21 +/- antibodies, in the presence or absence of 10 μM MG132. After 60 mins, cells were washed and lysates collected. Ub-TUBE IP was performed to pull-down ubiquitinated proteins from cell lysates. Inputs and elutions were analysed by immunoblot, using replicate membranes to detect TRIM21 and total ubiquitin. Results are representative of three independent biological replicates. **(C)** Schematic illustrating proposed model for intracellular degradation pathways of soluble monomeric TRIM21, or TRIM21 in complex with antibodies.

It is well established that cytosolic TRIM21 can undergo autoubiquitination, leading to its degradation together with any pathogen-antibody cargo (*64, 65*). To confirm whether TRIM21-GFP taken by macrophages can be ubiquitinated upon endo-lysosomal escape, we performed a pan-Ubiquitin-Tandem Ubiquitin Binding Entity (Ub-TUBE) IP. We pulled down ubiquitinated proteins from BMDM lysates after TRIM21-IC uptake, and detected TRIM21-GFP from the eluted Ub-TUBE material in both UT and MG132-treated BMDMs, with a stronger signal from MG132-treated cells (Fig. 6B). As expected, global ubiquitin levels were increased for MG132-treated BMDMs (Fig. 6B). Importantly, additional controls showed that TRIM21 feeding materials were not ubiquitinated prior to uptake (Fig. S4A). The ubiquitinated TRIM21-GFP which was pulled down by Ub-TUBE was heavier (∼70kDa) than endogenous WT TRIM21 (Fig. S4B).

Together these results suggest that, once taken up by macrophages, both free TRIM21 and TRIM21-IC can escape the endo-lysosomal compartment into the cytosol, undergo ubiquitination, and get degraded by proteasomes (Fig. 6C). For TRIM21-ICs, both lysosomal and proteasomal degradation pathways become engaged, with proteasomal degradation still preferred (Fig. 6C).

### TRIM21 enhances antigen cross-presentation

As we observed that TRIM21 and TRIM21-ICs could escape endo-lysosomal compartments into the cytosol, we reasoned that antigen cross-presentation via MHC Class I may occur (*66*). To examine whether TRIM21 influences antigen cross-presentation after uptake and escape, we utilised CD8^+^ T cells purified from the ovalbumin (OVA)-reactive OT-I transgenic mouse. We exploited TRIM21’s Fc-binding abilities to generate larger ICs from OVA, α-OVA antiserum, and GFP-trapped TRIM21 (OVA-α-OVA-TRIM21). These ICs were fed to IFNγ-primed BMDMs, and CD8α T cell activation in response to OVA was evaluated by flow cytometry (Fig. S5A, B). As controls, we used IFNγ-primed macrophages fed OVA alone, OVA-α-OVA antiserum, OVA-normal rabbit serum-TRIM21, or macrophages pulsed with the OT-I cognate peptide, SIINFEKL (Fig. S5A). Macrophages were primed to increase their capacity for cross-presentation. As expected, relative to OVA alone, OVA-α-OVA ICs increased T cell activation (CD69^+^ expression) at all concentrations tested (Fig. 7). At the lowest OVA concentration of 10 ng/mL, when larger complexes were made with TRIM21 (OVA-α-OVA-TRIM21), there was significant increase in T cell activation compared to all other controls, and trending towards the levels of activation induced by SIINFEKL peptide (Fig. 7A). This effect was specific for OVA-α-OVA-TRIM21 ICs, as activation was diminished when using the control “normal rabbit serum” (TRIM21 + normal rabbit serum + OVA), at all concentrations tested (Fig. 7). At higher OVA concentrations, OVA-α-OVA-TRIM21 still induced greater T cell activation than did OVA alone or OVA-normal rabbit serum-TRIM21, but no longer significantly higher than OVA-α-OVA ICs, as OT-1 T cell activation plateaued (Fig. 7B, C). TRIM21 + OVA (without antibody) was only sufficient to induce T cell activation at high OVA concentrations, and this was similar to the OVA control (Fig. 7C).

**Fig. 7.**
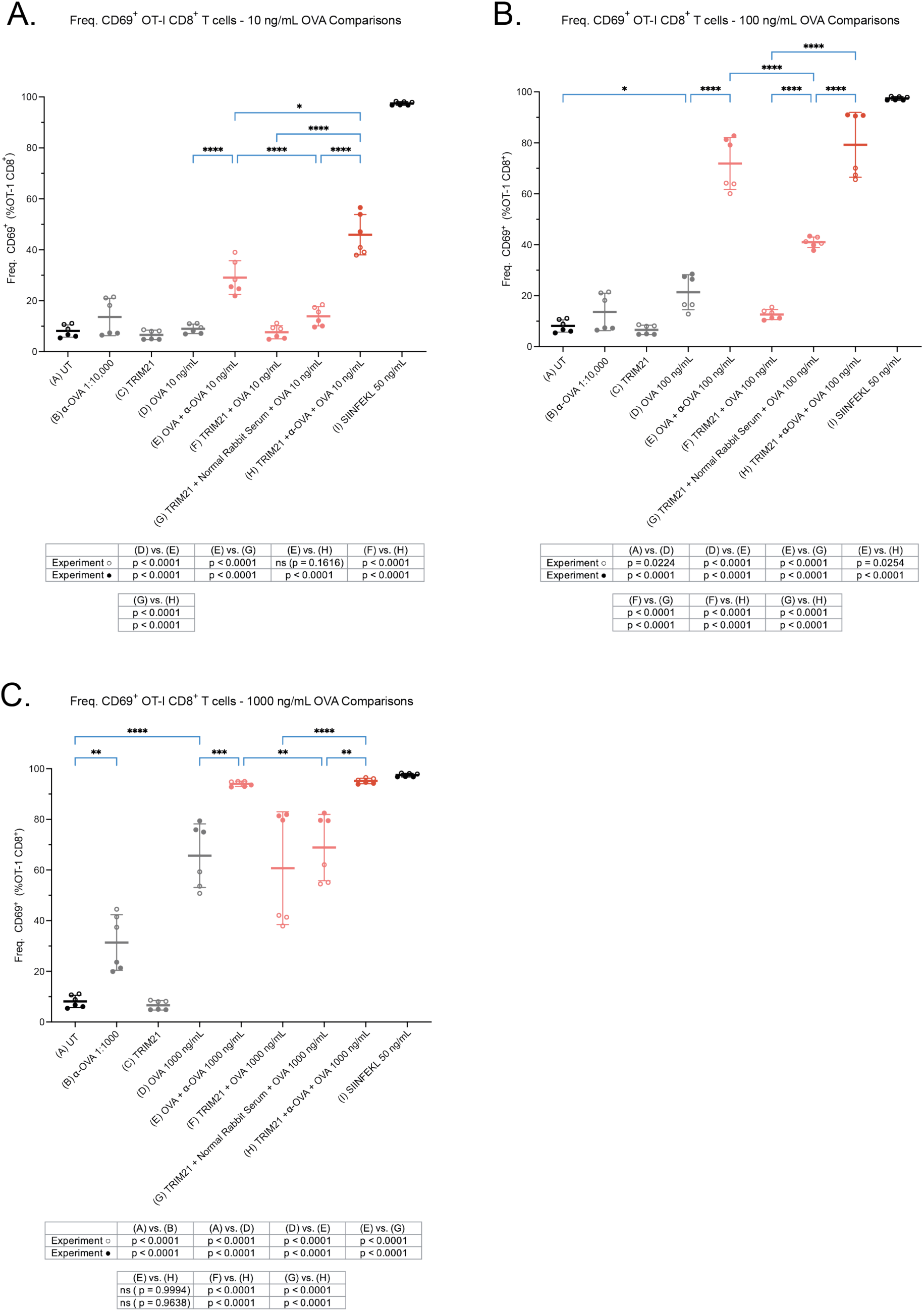
TRIM21 enhances antigen cross-presentation. BMDMs were stimulated with 100 ng/mL mIFNγ for ∼24 hours and fed with 10 μL of the following materials: OVA (+/- rabbit α-OVA), GFP-trap eluted TRIM21 (+/- OVA, OVA/α-OVA or OVA/normal rabbit serum), 50 ng/mL SIINFEKL peptide. Cells were incubated for 6 hours and co-cultured overnight with OT-I CD45.1 cells. OT-I/macrophage co-cultures were stained for extracellular CD45.1, CD8α and CD69. Frequency of CD69^+^ expression as a percentage of CD8^+^ OT-I cells was determined for a range of OVA concentrations: **(A)** 10 ng/mL, **(B)** 100 ng/mL, **(C)** 1000 ng/mL. Results from 2 biologically independent experiments were pooled, normalised to UT, and analysed by one-way ANOVA and Tukey’s post-hoc test. Duplicate experiments are distinguished by filled or empty symbols. Data represented as mean + SD for each condition. Tables denoting specific p values for each independent experiment are shown. * P ≤ 0.05, ** P ≤ 0.01, *** P ≤ 0.001.

Additionally, we observed downregulation of CD3 and T-cell receptor beta, variable 5.1 (Tcrb-V5.1), which corresponded to levels of T cell activation (Fig. S5C, D). Mean fluorescence intensity (MFI) levels for both CD3 and Tcrb-V5.1 were significantly lower for both OVA-α-OVA and OVA-α-OVA-TRIM21 ICs compared to TRIM21 + OVA, or TRIM21 + normal rabbit serum + OVA (Fig. 5SE, F).

Together these results suggested when complexed with OVA, TRIM21 was able to increase the uptake and antigen presentation of OVA, thereby enhancing OVA-specific T cell activation.

### TRIM21 uptake induces pro-inflammatory and metabolic changes in macrophages

When taken up by IFNγ-primed HMDMs, larger TRIM21 ICs induced higher TNFα cytokine secretion, with the level of TNFα tending to correlate with IC size (Fig. 8A). SjD59 plasma alone also induced some cytokine secretion, indicating that multiple mechanisms may contribute to pro-inflammatory HMDM activation (Fig. 8A). To assess changes in gene expression six hours after feeding, IFNγ-primed HMDM RNA was isolated and subjected to bulk RNA sequencing (RNAseq) analyses. Principal component analysis showed that PC1 divided the dataset according to the presence or absence of TRIM21, as highlighted by oval shadows (Fig. 8B).

**Fig. 8.**
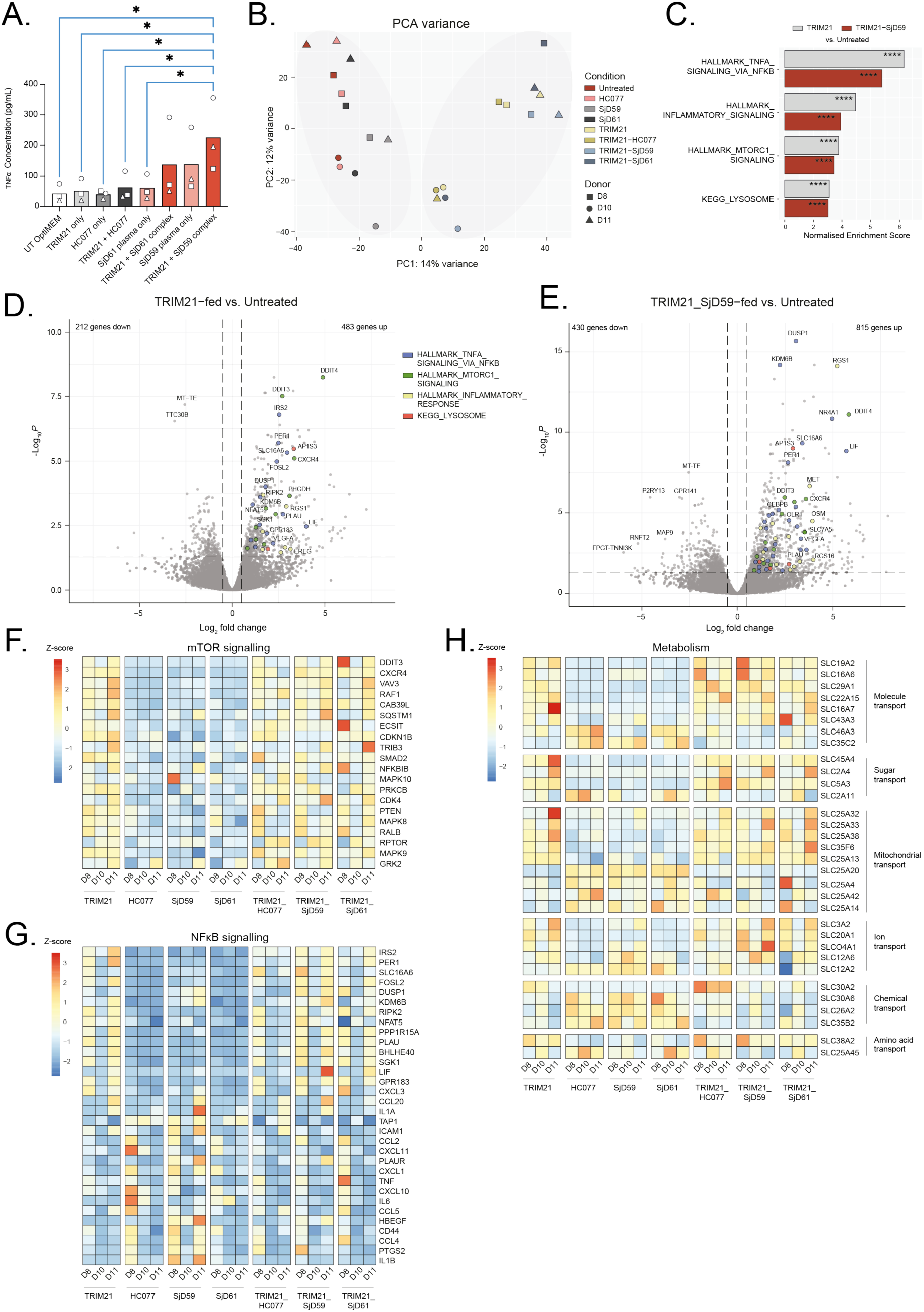
TRIM21 uptake by macrophages induces pro-inflammatory and metabolic changes. **(A)** HMDMs were stimulated with 100 ng/mL human IFNγ for ∼24 hours and fed with 10 μL of GFP-trap eluted TRIM21 (+/- complexed with indicated plasma). Cells were incubated for 6 hours, and SN were analysed by ELISA for human TNFα. Average results from 3 biologically independent experiments were pooled and analysed by one-way ANOVA and Fisher’s least significant difference post-hoc test. Each HMDM donor is represented by different symbols. * P ≤ 0.05. **(B-F)** HMDMs from 3 donors (D8, D10, D11) were fed with 20 μL of GFP-trap eluted TRIM21 (+/- complexed with patient plasma) and incubated for 6 hours. RNA was extracted, sequenced, aligned to reference genome and analysed. Significance was calculated using the Wald test (two-tailed), followed by Bonferroni correction. **(B)** PCA plot for each donor and feeding conditions (UT, plasma alone (SjD/HC), TRIM21 +/- plasma). **(C)** Normalised enrichment score plots for TRIM21 and TRIM21_SjD59 vs. UT feeding conditions for designated gene sets. Volcano plots show log_2_ fold changes in differentially expressed genes for **(D)** TRIM21 alone vs. UT and **(E)** TRIM21_SjD59 complex vs. UT feeding. Colours indicate genes assigned in Hallmark and KEGG gene set pathways. Heatmaps for donors and feeding conditions highlighting changes in **(F)** mTOR, **(G)** NFκB signalling, and **(H)** metabolism gene expression.

As these HMDMs were IFNγ-primed, inflammatory genes may already be highly expressed. Nevertheless, further gene expression changes were observed when comparing TRIM21 or TRIM21_SjD59 vs. UT conditions (Fig. 8C, D, E). No significant differential expression was shown by TRIM21_SjD59 or TRIM21_SjD61, vs. TRIM21_HC077 ICs (Fig. S6A, B). This may indicate that TRIM21-ICs alter the dynamics of uptake, but that at this 6-hour time-point, TRIM21 internalisation itself is driving the effects on gene expression.

Changes induced by TRIM21 uptake alone (vs. UT) included upregulation of genes in multiple pathways; notably inflammatory, lysosomal responses, TNFα via NFκB, and mTORC signalling pathways (Fig. 8C, D). Similar pathways were also altered for TRIM21_SjD59 feeding (vs. UT), with almost twice the numbers of genes upregulated (815) and downregulated (212) compared to TRIM21 alone (Fig. 8E). We used specific pathway gene sets (Hallmark, KEGG, and Biocarta), to discern global changes in interrelated genes from HMDMs fed with TRIM21 or TRIM21_SjD59 vs. UT. Gene sets with strong enrichment included TNFα via NFκB (Hallmark, Fig. S6D, E), ribosomal pathways (KEGG. Fig. S6F, G), and MAPK (Biocarta, Fig. S6H, I).

Noticeable TRIM21-induced changes in mTOR pathway genes, (Fig. 8F) and possibly NFκB signalling were observed (Fig. 8G). However, many NFκB signalling genes are shared with other pathways and thus, interpretation of these results requires caution (Fig. 8G). Changes in metabolic solute carrier (SLC) transporter genes were also detected (Fig. 8H). Transporter genes for various cargoes were upregulated, including those to transport molecules, sugars, ions and mitochondrial cargoes (Fig. 8H). Together, IFNγ-primed HMDMs exhibited changes in inflammatory and metabolic gene expression in response to TRIM21 uptake, whether in-complex with antibodies or alone.

Some plasma-only effects were observed, particularly for SjD59 plasma, showing increased expression of some genes in the mTOR and NFκB pathways (Fig. 8G), as well as multiple adhesion genes (Fig. S6C). This may suggest that high levels of autoantibodies in the patient’s plasma were sufficient to activate inflammatory and metabolic gene expression changes.

Combined, these results suggested TRIM21, alone or in complex with SjD patient plasma, induced transcriptional changes after uptake in IFNγ-primed macrophages. We primarily observed changes in pro-inflammatory and mTOR pathways, perhaps indicative of cell stress responses.

## DISCUSSION

TRIM21 studies have so far mainly investigated its intracellular antiviral and regulatory roles (*42*). For example, it has previously been suggested that dysregulation of TRIM21 autoubiquitination and turnover may drive downstream pro-inflammatory signalling and potentially promote autoimmunity (*15, 67*). Based on our observations, we propose an alternative model, whereby TRIM21 is released during lytic cell death. Once released, TRIM21 can bind antibodies outside the cell. By BN-Page, we determined that TRIM21-IC formation could be mediated by Fc domain binding in healthy people, and by both F(ab’)_2_ and Fc binding in SjD patients. Regardless, such binding may generate circulating IC which are detected and picked up by macrophages, where level of uptake correlates with the size of the IC. We speculate that should this occur in patients, it may contribute to the IC deposition and local autoinflammation that has previously been described in SjD patients’ SGs (*68, 69*).

Histopathology of SjD SG biopsies is widely applied in clinical practice, and is important for disease diagnosis, stratification and prognosis (*70*). The histological features identified by the H&E staining in this study correspond with known literature, including loss of tissue architecture, fibrosis, fatty deposition and focal infiltrates (*71*). However, few studies have performed multiplexed immunofluorescent imaging of SjD tissues. Reports so far are limited to investigating fibroblast niches (*72*), ectopic lymphoid structures (*73*) and stromal cell subsets (*74*). In this study, we expanded marker panels to include those for monocytes, granulocytes, immunoglobulins and the TRIM21 autoantigen. TRIM21 was ubiquitously expressed throughout SG tissues from both sicca controls and SjD patients, whilst multiple immunoglobulin subclasses were detected in close proximity to HLA-DR expression in SjD biopsies. These immunoglobulin findings correspond with previous studies, which show that focal infiltrates in SjD SG biopsies are rich in IgG and IgA-expressing B cells, compared to sicca controls (*75–77*). Together our findings indicated that antigen presentation mechanisms may occur in SjD SGs.

Additionally, SG IgD detection was striking and of interest with regards to the associations between IgD and control of autoreactive B cells (*78*). Previous SjD studies have correlated elevated levels of CD38^high^IgD^+^ B cells from peripheral blood, with anti-Ro seropositivity and disease activity (*79*). TRIM21 is highly promiscuous and known to bind all four human IgG subclasses, IgA and IgM. Whether TRIM21 binds IgD remains unclear, yet it is possible that IgD is a component of TRIM21-IC in vivo (*15*).

It had previously been shown that TRIM21 was upregulated by LPS stimulation (*43*), yet it remained unclear whether this upregulation was cell-type specific and could be induced with other pro-inflammatory stimuli. TRIM21 protein was upregulated by multiple stimuli in both BMDMs and HeLa. This is relevant in the context of SjD because pro-inflammatory IFN signatures have been detected in patient SG tissues and peripheral blood mononuclear cells (PBMCs). This includes the detection of the viral-inducible gene IFNα-inducible protein 27 (*80, 81*). Chronic viral infections, and in particular Epstein-Barr virus (EBV), have been strongly correlated with SjD (*82*). A combination of molecular mimicry of autoantigens, inflammatory cytokine stimulation, infection and induction of B cell proliferation and transformation, may together contribute to lymphoma risks in SjD patients (*83*). Many viral infections drive lytic cell death and, although the evidence is limited, one study found that EBV infection of human monocytes induced AIM2-dependent pyroptotic cell death (*84*). Pathogens such as EBV that gain entry through the mouth may drive localised upregulation of TRIM21 at SGs during anti-pathogen responses, leading to antigen release during pathogen-induced lytic cell death.

The difficulties of identifying true TRIM21-binding proteins are well-established. False-positive TRIM21 interacting partners may be identified, not due to direct interactions between TRIM21 and the protein of interest, but instead between TRIM21 and Fc domains of antibodies used for co-IP (*85*). To avoid this pitfall, we employed a GFP-trap micro-purification method to isolate TRIM21-GFP proteins and to detect TRIM21-immunogobulin co-IP. GFP-trap co-IP uses a nanobody-conjugated agarose to robustly and efficiently pull-down GFP-fusion proteins, thus eliminating the need for the addition of target antibodies (*86*). The advantages of utilising a TRIM21-GFP tagged protein were the following: the heavier protein (∼70 kDa) could be separately identified from endogenous TRIM21 (52 kDa) by immunoblot; the GFP could be detected by ELISA; and IC sizes could be quantified by FIDA which requires a fluorescent analyte sample (*87*).

Due to the known TRIM21-Ig binding, all our studies were performed in Opti-MEM to avoid confounding results due to the antibodies in foetal bovine serum. Ideally, to determine that TRIM21-IC uptake was FcR-mediated, FcR blockade control experiments would be performed. However, commercially available FcR-blocking reagents contain monoclonal antibodies against CD16, CD32 and CD64, although manufacturers do not share specifics (*88*). Therefore, we were unable to block TRIM21-Ig uptake as TRIM21 would instead bind the anti-FcR antibodies, giving false-positive uptake results.

It is widely acknowledged that phagocytosis is more efficient when target particles are opsonised with antibodies recognised by cell-surface FcRs (*62, 89*). FcR cross-linking by ICs drives uptake, downstream signalling cascades, and antigen presentation (*61*). Thus, it was unsurprising that in our study, TRIM21 uptake was faster and stronger when complexed with SjD patient plasma containing anti-TRIM21 antibodies.

We made use of the OT-I system, which is well-established for measuring CD8 T cell activation in response to OVA peptide presentation (*90, 91*). Additionally, emerging studies have indicated a role of granzyme K-producing cytotoxic CD8^+^ T cells, which are proposed to destroy epithelia in SjD (*36, 92*). Although we did not measure the presentation of TRIM21 itself, TRIM21 stimulated the cross-presentation of α-OVA-opsonised OVA protein. This suggests TRIM21 can bind opsonised antigens and directly enhance their uptake for cross-presentation. Although not directly tested, we cannot exclude the possibility that endogenous TRIM21 in the IFNγ-primed macrophages, could engage the internalised α-OVA-OVA complexes to drive cross-presentation. Additionally, whether such enhancement occurs only via increased antigen uptake and/or via enhanced endo-lysosomal escape pathways, remains to be elucidated.

We found that plasma containing larger TRIM21-ICs stimulated macrophage TNFα production. It has previously been shown that combining sera from systemic lupus erythematosus with necrotic SN containing nuclear antigens, induced TNFα secretion from U937 cells (*93*). Therefore, our results support this understanding that ICs formed from patient sera and autoantigens can activate macrophages and drive downstream pro-inflammatory signalling.

Multiple gene expression changes were observed in macrophages treated with TRIM21 alone or complexed with antibodies (SjD/HC), vs. UT. Genes in inflammatory, lysosomal, mTOR and TNFα (via NFκB) pathways were upregulated. Together these may suggest lysosomal stress responses are promoted, perhaps via lysosomal damage which activates mTOR, triggers inflammation and may permit lysosomal escape for cross-presentation (*94–96*). Fewer differences were observed when comparing TRIM21_SjD vs. TRIM21_HC, suggesting that the expression changes may not all be IC size-dependent. However, larger ICs (TRIM21_SjD61/TRIM21_SjD59) upregulated some mTOR pathway genes. This corresponded to the upregulation of SLC transporter genes, known to contribute to mTOR pathway activation (*97*). mTOR is a central regulator of metabolism (*98*), specifically regulating phagosomal fusion and termination of autophagy after macroendocytosis (*99*). It is possible that after the uptake of TRIM21-ICs, mTOR is activated to deal with the uptake of these large protein complexes and promote cellular recovery (*100*).

Together, our results suggest TRIM21 may be particularly immunogenic due to its inherent capacity for binding antibody Fc domains and forming ICs. In the context of SjD, SG damage may allow extracellular TRIM21 release, which binds to circulating antibodies and possibly forms ICs. Together with high IFN levels in SjD patients, this may drive TRIM21 uptake, inflammation, metabolic changes, antigen presentation, and TRIM21/Ro52 autoantibody production.

### Limitations of the study and future perspectives

This study showed that *in vitro*, TRIM21 binds antibodies outside the cell and forms higher-order ICs with SjD patient plasma. These complexes are taken up and activate macrophages. Due to a lack of suitable animal models which faithfully recapitulate SjD, these experiments were performed using the most appropriate available primary immune cells and cell lines. Despite this, our *in vitro* studies cannot reproduce the milieu of immune and stromal cells which may contribute to autoimmunity *in vivo*.

Due to limited patient plasma and TRIM21-GFP protein purification methods, our study was limited to small-scale methods for detecting uptake and activation. Hence, we were unable to identify the specific mode of TRIM21 uptake, and how TRIM21-GFP escapes endosomes into the cytosol for ubiquitination. Future studies could test specific uptake methods by flow cytometry and imaging of different organelles e.g. galectins, to define such pathways. Additionally, it would be worth testing whether TRIM21 itself is presented by macrophages or DCs after uptake, thus providing possible mechanisms for autoimmune T cell activation. If more SjD plasma was available, sensitive proteomic analyses could be performed to determine whether TRIM21 antigen is detected in SjD patient plasma.

In SjD SG biopsies, we identified immune cell infiltration, glandular deformation and fat deposition. Due to limitations with longitudinal biopsy cohorts, only biopsies of late-stage, dysfunctional SGs from patients were imaged. Taking multiple biopsies is not preferred by patients, or considered ethical outside of a trial where patients benefit with treatments. We propose that lytic cell death may contribute to SG disease pathology, however we were unable to observe specific markers of pyroptosis/necroptosis/necrosis/apoptosis using immunofluorescent staining due to tissue atrophy and lack of suitable reagents. Additionally, it remains unclear what happens in SjD SGs during early disease stages, before visible pathology. Unfortunately, accurately identifying markers of specific forms of cell death on fixed tissues remains challenging (*101, 102*). As new cell death probes are developed for immunofluorescence imaging, future studies could investigate whether forms of lytic cell death occur in SjD SGs.

Our study investigates one potential mechanism for autoimmune induction; however, there are additional contributing factors which were not explored, including HLA associations and possible pathogen contributions such as those from EBV. These are worth investigation to explore why lytic cell death in healthy individuals does not drive autoimmune responses, despite the possibility of TRIM21 binding circulating antibodies.

## MATERIALS AND METHODS

### Study Design

This study was designed to assess mechanisms for how intracellular TRIM21 may become an autoantigen, using various in vitro experiments.

Anonymised human samples used in this study were obtained from the following sources. Tonsillar tissue was processed and provided by the Oxford Centre for Histopathology Research. Commercially supplied SjD patient plasma samples were obtained from NeoBiotech (custom order) and normal serum controls from Invitrogen. HC plasma was originally obtained as part of a study approved by the Institutional Review Board at Rockefeller University Hospital (RUH) under human subjects protocol LDU-0437. Healthy donors were recruited through the RUH outpatient clinic. All donors provided written informed consent according to the Declaration of Helsinki, before enrolment. Samples were coded and anonymised. Blood was collected into acid citrate dextrose-treated vacutainer tubes; plasma was isolated by centrifugation within one hour of collection, aliquoted, and stored at -80°C until used. Storage and use of these samples at the University of Oxford was approved by the Berkshire Research Ethics Committee, REC; 13/SC/0466 (*103*). SjD and sicca syndrome patient biopsies were obtained from the Optimising Assessment of Sjögren’s Syndrome (OASIS) cohort (*104*). All plasma donors, and subjects in the OASIS cohort, signed written informed consent according to the principles of Helsinki. The OASIS study was approved by the Wales Research Ethics Committee 7 (WREC 7), formerly Dyfed Powys REC; 13/WA/0392. FFPE slides from the OASIS cohort were used for H&E and Cell DIVE multiplex imaging. The images represented in this study were chosen to reflect the SG damage, cellular infiltration and possible immune mechanisms present in SjD patients.

Due to limited availability of SjD patient plasma, we developed small-scale purification and in vitro methods to form ICs, and perform uptake and antigen presentation assays. Where possible, experiments were performed and analysed as independent triplicates. For experiments using SjD patient plasma, experiments were executed as independent biological replicates. We chose 3 SjD patients from the LIPS assay to cover the range of autoantibody titres. 3 HC plasmas (autoantibody negative) were also used. These same patient plasmas were used throughout the study. For BMDM and HMDMs experiments, different donors (mouse/human) were used for each replicate.

### Mice

C57BL/6 mice served as WT (Charles River). GSDMD^-/-^ mice, carry a CRISPR/Cas9-derived knock-out allele of the mouse *Gsdmd* gene (The Jackson Laboratory). C57BL/6 and GSDMD^-/-^ mice were used to derive BMDMs. OT-I mice (Charles River) were bred with CD45.1 mice (Charles River) to generate congenitally marked OT-I CD45.1 cells. Mice were bred in-house and kept in specific pathogen-free conditions, according to ethical standards approved by the UK Home Office and University of Oxford. Mice were kept in individually ventilated cages with environmental enrichment at 20–24 °C, 45–65% humidity with a 12 hour light/dark cycle (7am– 7pm) with half an hour dawn and dusk period. For experiments, a mixture of male or female mice aged 9-14 weeks were used, and independent replicates were performed using cells from different mice.

### Cell Lines

HeLa cells were kindly gifted by Dr. Iona Manley (Dunn School of Pathology, Oxford University) (originally purchased from ATCC). HEK293c18 (clone CRL-10852) were obtained from ATCC. HEK293T cells were kindly gifted by Prof. Michael Dustin (Kennedy Institute of Rheumatology, Oxford University). Cell lines were cultured in complete DMEM (DMEM supplemented with 10% heat-inactivated FBS, 5% Glutamax) at 37 °C and 5% CO_2_. For maintenance growth, cells were cultured in T75 vented tissue-culture-treated flasks. Cells were washed with DPBS, trypsinized with Gibco™ Trypsin-EDTA (0.05%) for cell detachment and centrifuged at 300 *xg* for 5 mins. The cell pellet was resuspended in pre-warmed media, cell counts and viability were measured using an automated cell counter. Cells were seeded at desired confluency as described in the relevant sections.

### Media and Supplements

HeLa/HEK293c18/HEK293T media (complete DMEM): DMEM supplemented with 10% heat-inactivated FBS, 5% Glutamax RPMI media for HMDMs: RPMI, supplemented with 10% FBS, 1× Pen/Strep/Glutamine RPMI media for isolated T cells: RPMI 1640 (without glutamine), supplemented with 10% FBS, 1x Pen/Strep/Glutamine, 50μM 2-Mercaptoethanol BMDM differentiation media: DMEM, supplemented with 10% FBS, 1× Pen/Strep/Glutamine, 50 ng/mL macrophage colony-stimulating factor (M-CSF) HMDM differentiation media: RPMI, supplemented with 10% FBS, 1× Pen/Strep/Glutamine, 100 ng/mL M-CSF IMDM media: IMDM, supplemented with 10% FBS, 1× Pen/Strep/Glutamine Serum-free media: DMEM, 1% Nutridoma, 1% Glutamax

### Generation of Primary Mouse BMDMs

Total mouse bone marrow (C57BL/6 and *Gsdmd*^-/-^) was isolated, cultured in 4 x non-TC treated 100 mm sterilin square dishes per mouse, and differentiated for 6 days in complete DMEM using human recombinant M-CSF (50 ng/mL), at 37 °C with 5% CO_2_. On day 3, 7 mL supplemented media was added and on day 6, adherent cells were harvested using ice-cold PBS and cell lifters and cryopreserved at a concentration of 10 million cells/vial using cold, sterile-filtered FBS with 10% DMSO. Cells were placed in liquid nitrogen for long-term storage.

### Isolating Mouse OT-I T cells

Spleens and lymph nodes were harvested from OT-I-positive mice and mechanically homogenized by mashing though a 70μM cell strainer using a syringe plunger. The cell suspension was then washed in 1x PBS and counted for CD8+ T cell isolation. CD8^+^ T-cell isolation was conducted using the MojoSort^TM^ CD8^+^ T-cell isolation kit and magnets, in accordance with the manufacturer’s protocol. Isolated OT-I cells were resuspended in complete RPMI (see media and supplements) and used for antigen presentation assay.

### Buffers and Solutions

Tris-buffered saline with tween (TBS-T): 200 mM Tris-HCl, 1500 mM NaCl, 0.05% Tween® 20 (pH 7.6)

TBS-T, supplemented for primary antibodies: TBS-T containing 5% skimmed-milk powder, 0.02% sodium azide

PBS-T: 1X PBS, 0.05% Tween® 20

Blocking buffer: 1% BSA in 1X PBS

Denaturing lysis buffer: 66 mM Tris-HCl (pH 7.4), 2% SDS 1X TBS pH 7.4: 150 mM Tris-HCl, pH7.4, 50 mM NaCl

Glycine elution buffer: 100 mM Glycine, pH 2.5

Pierce IP-lysis buffer, supplemented: For 10 mL lysis buffer; 400 μL 25X protease inhibitor cocktail solution (prepared as 1 pill in 2 mL dH_2_0), 1 pill of phosphatase inhibitor

LIPS lysis buffer: For 50 mL; 50 mM Tris, pH 7.5, 100 mM NaCl, 5 mM MgCl_2_, 1% Triton X-100, 50% glycerol and protease inhibitors (2 tablets of complete miniprotease inhibitor cocktail)

LIPS assay buffer A: 50 mM Tris, pH 7.5, 100 mM NaCl, 5 mM MgCl_2_, 1% Triton X-100

NativePAGE™ sample buffer: For 1 mL lysis buffer; 250 μL NativePAGE™ Sample Buffer (4X), 40 μL 25X protease inhibitor cocktail solution (prepared as 1 tablet in 2 mL dH_2_O), 100 μL DDM (from 10% stock), 610 μL dH_2_O

HBSS supplemented: HBSS, 0.3 mM EDTA MACS buffer: HBSS, 3% FBS, 10mM EDTA

Custom-GFP ELISA blocking buffer: 1% BSA in 1X PBS

Flow cytometry buffer: 2% FCS, 2.5mM EDTA, and 0.01% sodium azide in 1x PBS Cell DIVE mounting medium: 90% glycerol, 1% DABCO, 4% propyl gallate

Cell DIVE bleaching solution: 0.1 M NaHCO_3_ pH 11.2, 3% H_2_O_2_

### Resurrecting, Culturing and Plating BMDMs for Experiments

Cryopreserved day 6 BMDMs were thawed, diluted in complete media and centrifuged at 500 *xg* for 3 mins at room temperature to remove DMSO. The pellet was resuspended in complete, supplemented DMEM containing 50 ng/mL M-CSF. Cells were plated in 10 cm non-tissue-culture-treated petri dishes and incubated for 48 hours recovery at 37 °C with 5% CO_2_. After 48 hours, BMDMs (day 8) were harvested using ice-cold PBS and cell scrapers, and counted using an automated cell counter. Cells were seeded at a 5 × 10^4^ cells/mL density for all experiments unless otherwise specified, in 100 μL per well in a 96-well plate, in supplemented DMEM containing 50 ng/mL M-CSF. After plating, the cells were left undisturbed at room temperature for ∼20 minutes to promote even attachment density. The plate was incubated at 37 °C with 5% CO_2_ for 16 hours prior to experimentation. For uptake experiments, cells were primed with 100 ng/mL murine IFNγ (mIFNγ) for ∼24 hours.

### Isolating CD14^+^ Human Monocytes

PBMCs were isolated from anonymised, healthy donor blood cones (obtained from NHS Oxford blood bank). Blood was removed from the cone and volume was adjusted using HBSS supplemented with 0.3 mM EDTA to a final 25 mL volume. PBMCs were isolated by Ficoll gradient centrifugation (700 *xg,* 25 mins, room temperature, without brake), then transferred to a fresh tube, resuspended in a final volume of 50 mL in supplemented HBSS, and washed twice (500 *xg*, 10 mins at room temperature). The cell pellet was resuspended in 1 mL MACS buffer (HBSS, 3% FBS, 10 mM EDTA).

CD14^+^ monocytes were positively selected using MagniSort™ Positive Selection Protocol. Briefly, 200 μL MagniSort™ Enrichment Antibody was added to cells in MACS buffer and incubated for 10 mins at room temperature. An additional 3 mL MACS buffer was added and cells were pelleted at 500 *xg* for 3 mins at room temperature. The pellet was resuspended in 1 mL MACS buffer. Mixture was transferred to sterile, polystyrene FACS tubes. 300 μL MagniSort™ Positive Selection Beads were added and the mixture was vortexed vigorously before a 10-minute incubation at room temperature. Volume was adjusted to 2.5 mL in MACS buffer. FACS tubes were inserted into EasySep™ Magnet, incubated for 5 mins and the SN poured away (discarding CD14^-^ cells). Beads were resuspended in 2.5 mL MACS buffer, magnetised and SN poured away 3 more times. The bead-bound CD14^+^ cells were added to 30 mL complete IMDM (10% FBS, 1× P/S/G). Cells were counted and frozen in 90% FBS, 10% DMSO at a concentration of 3 × 10^7^ cells/vial. Cryopreserved cells were stored in liquid nitrogen.

### Generation of HMDMs

Frozen day 0 CD14^+^ monocytes were thawed, diluted in complete RPMI and centrifuged at 500 *xg* for 3 mins at room temperature to remove DMSO. The pellet was resuspended in complete RPMI supplemented with 100 ng/mL M-CSF. Cells were plated in 10 cm tissue-culture-treated petri-dishes and incubated for 7 days recovery at 37 °C with 5% CO_2_. On days 2 and 5, cells were fed with 5 mL complete RPMI containing 100 ng/mL M-CSF. On day 7, media containing floating DC-like monocytes was aspirated and 10 mL cold PBS was added. Dishes were incubated in the fridge for 10 mins to aid detachment and cells were lifted using cell lifter. Cells were counted and seeded at a 5 × 10^4^ cells/mL density for all experiments, in 100 μL per well in a 96-well plate, in supplemented RPMI containing 100 ng/mL human M-CSF and 100 ng/mL human IFNγ (hIFNγ) for ∼24-hour priming (for uptake and activation assays).

### BMDM Stimulation Experiments (TRIM21 Upregulation)

Day 8 BMDMs were plated as described above and incubated for 16 hours. On the day of experimentation, media was aspirated and replaced with 200 μL complete DMEM +/- priming substances from the following: ultra-pure LPS (100 ng/mL), murine TNFα (100 ng/mL), poly I:C (2.5 µg/mL), murine IFNα (100 ng/mL), murine IFNβ (100 ng/mL), mIFNγ (100 ng/mL). At 4, 8 and 24 hours, media was aspirated, replaced with 30 µL lysis buffer (66 mM Tris-HCl (pH 7.4), 2% SDS) and left for 2 mins at room temperature. Triplicate lysate wells were pooled and stored at -20 °C.

### WT HeLa Stimulation Experiments (TRIM21 Upregulation)

16 hours prior to experimentation, HeLa cells were resuspended and plated to a density of 15,000 cells/well in a 96-well plate, in complete DMEM. On the day of experimentation, media was aspirated and replaced with 200 μL complete DMEM, containing priming substances. Cells were primed with hIFNγ at concentrations of 10 ng/mL, 100 ng/mL, 1 μg/mL. At the indicated time points, media was aspirated, replaced with 30 µL lysis buffer and left for 2 mins at room temperature. Triplicate lysate wells were pooled and stored at -20 °C.

### SN Precipitation

SN stored in a 96-well plate at -20 °C were retrieved and thawed at room temperature. The plate was centrifuged at 500 *xg* for 5 mins at room temperature and SN transferred to microcentrifuge tubes. Samples were mixed with an equal volume of methanol and 0.33 volumes of chloroform, vortexed for 5s, and centrifuged at 4°C for 10 mins at max speed. The upper phase (of 3 visible phases) was aspirated and 1.33-fold SN volume of methanol was added. Samples were vortexed and centrifuged as above. The SN was removed, leaving a final pellet which was dried at room temperature. Samples were dissolved in 40 μL 1x Laemmli loading buffer containing 10 mM DTT, prior to immunoblotting.

### Immunoblotting

Cell lysates were thawed at room temperature and mixed with 0.33 volumes of 4x Laemmli loading buffer containing 10mM DTT. Samples (Lys/SN) were boiled at 95 °C for 5 mins and centrifuged at max speed for 2 mins prior to loading the gel. Proteins were separated on 4–20% precast polyacrylamide gels, and transferred onto nitrocellulose membranes using a Trans-blot Turbo (Bio-Rad). Antibodies for immunoblot were against TRIM21 (1:2000), LC3 (1:1000), Ubiquitin (1:1000) GAPDH (1:2000), Tubulin (1:2000). All antibodies were prepared in TBS-T containing 5% skimmed-milk powder, 0.02% sodium azide. Proteins were visualised by ECL detection.

### BMDM Cell Death Induction

For all experiments, unless otherwise specified, complete media was replaced with 200 µL Opti-MEM on the day of stimulation. All lyophilized drugs were dissolved in DMSO and unless specified, drugs were diluted to given final concentrations in Opti-MEM. Stimulations were performed at given time-points and after each treatment, the 96-well plate was returned to the incubator at 37 °C with 5% CO_2_.

To induce intrinsic apoptosis, BMDMs were stimulated with apoptosis-inducing drugs ABT-737 (250 nM) and S63845 (250 nM) and incubated for 3 hours. For both necroptosis and pyroptosis inductions, cells were first primed with ultra-pure LPS (100 ng/mL) for 4 hours. For pyroptosis, cells were then treated with Nigericin (10 μM) for an additional 2 hours. For glycine-inhibited pyroptosis treatment, sterile-filtered glycine (20 mM final concentration) was added 30 minutes before Nigericin treatment. For necroptosis, in the last 30 minutes of LPS priming, cells were treated with Q-VD-Oph (10 μM), before final stimulation using SM AZD 5582 (1 μM) for 5 hours. To inhibit necroptosis, the inhibitor Nec1 (10 μM) was added at the Q-VD-Oph time-point.

To induce AIM2 pyroptosis, cells were instead seeded at a density of 100,000 cells/well of a 96-well plate, in complete DMEM containing 50 ng/mL M-CSF, and incubated overnight. Media was not replaced in this case. Cells were treated per well with either; 0.25% Lipofectamine-2000 alone as vehicle control, 200 ng ctDNA (2 µg/ml) alone, or combined ctDNA plus Lipofectamine-2000 (0.25%). 10 µL media was removed from the top of each well undergoing transfection and 10 µL of the appropriate treatment/transfection mixture was added to the cells. The plate was centrifuged for 10 mins, 1000 *xg* at room temperature and transferred to 37°C for 4 hours.

To induce necrosis, BMDMs were treated with 0.1% Triton X-100 in 200 µL Opti-MEM. For needle shearing, media was replaced with 200 µL cold PBS to aid cell resuspension. Using a 30G needle, cells were forcibly sheared multiple times in the PBS. The plate was centrifuged at 500 *xg* for 5 minutes to separate the cells and SN.

At end points, SN were collected in fresh 96-well plates and stored at -20 °C. For lysate collection, 30 µL lysis buffer was added to the wells and left for 2 mins at room temperature. Lysates were collected and stored at -20 °C.

### LDH and Cell Viability Assays

LDH release was measured using the CytoTox 96® Non-Radioactive Cytotoxicity Assay kit, according to the manufacturer’s instructions. In brief, 20 µL of cell SN was transferred to ELISA plates. 20 µL assay substrate was added and the plate was covered in the dark for approximately 20 minutes. After a deep cherry-red colour was visible in the 100% lysis control, the reaction was stopped with 20 µL stop solution. Absorbance was measured at 490 nm and 690 nm. Final values were calculated as A490-A690.

Cell viability was measured using the CellTiter-Glo Luminescent Cell Viability Assay, according to the manufacturer’s instructions. In brief, 20 μL of CellTiter-Glo® Reagent was added to 20 μL of cell culture substrate. The plate was mixed on an orbital shaker for 2 minutes at room temperature, then incubated for 10 minutes for signal stabilisation. Luminescence was measured by plate reader. % LDH release and % cell viability values were normalised to 100% lysis treatments (Triton X-100 treated) and UT controls respectively.

### TRIM21 siRNA Knockdown

BMDMs were plated in 96-well plates, and incubated overnight (no simulation). Media was aspirated and replaced with OptiMEM. siRNA knockdown was performed according to the manufacturer’s recommendations. Per well, cells were transfected with 4 pmol of the following siRNA; siGENOME non-targeting siRNA control pools, or ON-TARGETplus mouse Trim21 siRNA, SMART pool, combined with 0.75 μL Lipofectamine RNAiMax in OptiMEM. After 6 hours, OptiMEM was replaced with cDMEM with M-CSF. Lysates were collected 72 hours after transfection for immunoblotting.

### TRIM21 Constructs

TRIM21 antigen constructs were designed for subcloning into fluorescent vectors. Sequences encoded the open reading frames of TRIM21/Ro52 protein (Gene ID: 6737, NCBI Reference Sequence: NM_003141.4). Sequences were modified to contain the following: suitable restriction sites, Kozak sequence upstream of the open reading frame, and, for GFP fusion proteins, a GGSSGG linker between the last codon of TRIM21 and the first codon of the mEGFP tag. WT TRIM21 (w_PRYSPRY) is a full-length construct (aa 476), whilst TRIM21_NO_PRYSPRY is a truncated protein construct with no PRYSPRY (aa 1-276). TRIM21_FcMUT (aa 176) contains 2-point mutations (W381A/W383A) within the PRYSPRY domain.

### GFP Fusion Proteins

cDNA sequences encoding antigens of interest were subcloned N-terminally to the fluorescent-protein tag of the pHR-mEGFP expression vector, kindly supplied by Dr. Alexander Mørch (Kennedy Institute of Rheumatology, University of Oxford). The following antigen sequences were purchased from ThermoFisher Scientific; TRIM21_w_PRYSPRY TRIM21_NO_PRYSPRY, TRIM21_FcMut, HIV_NES, Sequences were subcloned into the fluorescent plasmid vector using standard digestion and ligation protocols. Full sequences available upon request.

### Lentiviral Transduction

To prepare lentivirus, HEK293T cells were plated to a density of 1 × 10^6^ cells/well in 2 mL supplemented DMEM in 6-well TC-treated culture plates and incubated overnight at 37 °C and 5% CO_2_. After 24 hours, they should be approximately 80% confluent. On the day of transfection, the following mixtures were prepared (volumes for 1 well); to 100 μL serum-free DMEM, 4 μL GeneJuice transfection reagent was added and incubated for 5 mins at room temperature. Then each of the following plasmids was added; TRIM21-mEGFP fusion protein vectors (see above) for transfer (0.50 μg), pMD.G (VSV-G envelope expressing plasmid) packaging vector (0.50 μg) and P8.91 (plasmid expressing gag and pol) packaging vector (0.50 μg). The total mixture was incubated at room temperature for 15 minutes. Transfection mixtures were added to the relevant wells and the plate was incubated for 72 hours. SN were harvested, clarified by centrifugation, and passed through a 0.45 μm filter to be used for transduction on the day of harvest.

One day prior to transduction, HeLa cells were seeded at a density of 5 × 10^4^ cells/well in 2 mL supplemented DMEM in 6-well TC-treated culture plates. After overnight culture (37 °C, 5% CO_2_), cultures were transduced by adding 1.5 mL freshly-harvested filtered lentiviral suspension to the cells in 2 mL supplemented DMEM (3.5 ml final volume), and incubated for up to 72 hours for optimal transduction efficiency. Cells were checked for GFP expression and passaged as necessary for recovery and expansion.

### Apoptosis Live Confocal Microscopy, Cells Treated with Drugs at Microscope

HeLa_TRIM21_w_PRYSPRY-mEGFP were seeded in µ-Slide 8 Well High ibiTreat culture plates at a density of 15,000 cells/well in 300 μL supplemented DMEM and incubated overnight at 37°C and 5% CO_2_. On the day of experimentation, media was aspirated and replaced with 300 μL serum-free media (DMEM1% Glutamax, 1% Nutridoma). 30 minutes before imaging, 1 μM final concentration of SiR-DNA stain was added to the centre of each well. Live microscopy was performed on a Zeiss 980 confocal microscope, with additional 37°C incubation, 5% CO_2_ and definite focus (autofocus) settings. Apoptosis-inducing drugs were added during live time-lapse microscopy at the following final concentrations; 1 μM ABT, 10 μM MCL1i.

### Generating Luciferase (Luc) Antigens for LIPS assay

HEK293c18 cells were plated to a density of 10 million cells in 20 mL media in p150 tissue-culture treated petri-dishes and incubated overnight. The following day, the media was replaced with serum-free media for transfection. Plasmids encoding the autoantigen-Renilla luciferase fusion proteins, i.e. pREN-2_Ro52_Luc (encoding a truncated TRIM21 lacking the PRYSPRY domain, fused to luciferase), pREN-2_Ro60_Luc, and pREN-2_La_Luc, were generously provided by Dr. Peter Burbelo, National Institute of Dental and Craniofacial Research, NIH, USA (*105*). For transfection, FuGENE® HD was combined with 6 µg pREN-2_Antigen_Luc plasmid in a 3:1 FuGENE® HD:DNA ratio, according to the manufacturer’s recommendations, diluted to a final 300 µL volume in Opti-MEM. The mixture was gently inverted and left under the hood at room temperature for 10 minutes. The FuGENE® HD-DNA mixture was added to plates in a dropwise manner, plates were gently swirled and placed in the incubator for 48 hours.

Lysates were prepared after 48 hours, using a protocol modified from Burbelo et al. 2009 (*106*). 50 mL cold LIPS lysis buffer was prepared, as described in buffers and solutions section above. Transfected HEK293c18 cells were washed with PBS, SN aspirated, 4.2 mL cold LIPS lysis buffer added, and cells harvested with cell scrapers. Lysates were transferred into sterile 1.5 mL microfuge tubes stored on ice. Cells were lysed using the Bioruptor® Pico sonication device with the following settings; 3 cycles of 30 secs on, 30 secs off. Lysates were centrifuged at 15,000 *xg* for two 4-minute spins at 4°C. After the first spin, tubes were inverted to remove loosely attached debris from the sides of the tubes. SN were transferred to new tubes on ice. Final Luc-antigen lysates were stored at -80° C for long-term storage.

Luciferase activity for each Luc-antigen was measured using a luminometer (TriStar² S LB 942, Berthold Technologies). The light units (LU) per μL of lysate were measured by diluting 1 μL lysate in 8 μL of PBS in wells of Berthold White 96-well microplates (performed in triplicate). Luminometer settings were edited to directly add 100 μL of 1X *Renilla* luciferase substrate to the diluted mixture and immediately measure luminescence with a 5 second read.

### LIPS Assay

40 μL of LIPS Assay buffer A was added to each well of a V-bottomed 96-well plate. All plasma samples were diluted 1:10 in buffer A. Positive control polyclonal Ro60/Ro52/La antibodies (commercially available) were diluted to a final 0.01 mg/mL concentration in buffer A. 10 μL of prepared plasma/antibody samples was added in triplicate to wells of the V-bottomed 96-well plate already containing 40 μL of buffer A. A master mix for each Luc-antigen lysate was prepared such that there were 1 X 10^7^ LU/50 μL buffer A. 50 μL of this Luc-antigen mixture was added to each relevant well of the V-bottomed 96-well plate. The plate was covered and incubated for 1 hour at room temperature on a rotary shaker.

After incubation, 5 μL of a 30% suspension of Ultralink protein A/G beads (Pierce Biotechnology, Rockford, IL) in PBS was added to the base of each well of a new opaque 96-well vacuum filter plate (Millipore). 100 μL Luc-antigen-antibody reaction mixture was transferred from the V-bottom 96-well plate to the new filter plate, mixed gently and incubated on a rotary shaker for 1 additional hour at room temperature.

The plate was washed using a vacuum manifold plate wash setup, according to manufacturer’s recommendations. Each well was washed 8 times with 100 μL Buffer A, then twice with 100 μL PBS. After the final wash, the plate was removed and carefully blotted dry to remove any remaining moisture. Luciferase activity was measure using a Berthold Luminometer with the following settings; 50 μL of 1X *Renilla* luciferase substrate was directly added into the wells, with 2 second shake, followed by 5 second luminescence read.

### Preparing HeLa cells for IP/ELISA – inducing cell deaths

Unless otherwise stated, prior to treatments, HeLa_TRIM21_w_PRYSPRY-mEGFP or HeLa_TRIM21_NO_PRYSPRY were seeded in 96-well culture plates at a density of 20,000 cells/well in 100 μL supplemented DMEM and incubated overnight at 37°C

#### Pyroptosis

Note cells were pre-primed with 10 ng/mL hIFNγ during seeding. For pyroptosis induction, cells were treated with an LPS/lipofectamine mixture prepared as follows (material for 1 well); to 10 μL pre-warmed Opti-MEM, 2 μL LPS from 1 mg/mL stock was added (or 2 μL Opti-MEM for mock transfection control). The LPS-Opti-MEM mixture was vigorously vortexed for 30 seconds and centrifuged at briefly maximum speed at room temperature. 0.5 μL Lipofectamine 2000 was added, the mixture was flicked 10-15 times to ensure even mixing and incubated at room temperature for 20 minutes under the hood. During the incubation, media was aspirated from the cells and replaced with 200 μL pre-warmed Opti-MEM to wash the cells and remove FBS. This Opti-MEM was removed and replaced with 100 μL Opti-MEM. After 20 minutes, an additional 90 μL Opti-MEM was added to the transfection mixture. 100 μL LPS-transfection mixture was added to the cells (final volume should be 200 μL) and cells were centrifuged for 5 minutes, 500 *xg* at 37 °C. The plate was returned to the incubator and left for 24 hours to ensure complete lytic death.

#### Apoptosis

To induce apoptosis, HeLa cultures were washed once with Opti-MEM and media was replaced with 200 μL fresh Opti-MEM. HeLa were then treated for 2h with ABT-737 (1 μM) and S63845 (10 μM). After 2 hours, the top 150 μL SN was collected into low-protein binding microfuge tubes, without disturbing the cells, and stored at 4°C for downstream analyses within 24 hours.

#### Detergent lysis

For detergent lysis, cell monolayers were washed with PBS and cells resuspended in pre-warmed trypsin/EDTA (50 μL per well). After resuspension, trypsin was inactivated with 150 μL warm media and cells pelleted in low-binding microfuge tubes (5 mins, 300 *xg*, 4°C). Cell pellets were washed in 1 mL PBS and centrifuged again for 5 mins, 300 *xg* at 4°C. Pellets were resuspended in cold Pierce IP lysis buffer at the same volume as initial collection. Tubes were rotated at 4°C for 30 mins, then centrifuged at maximum speed (5 mins, 4°C). SN were transferred to fresh low-binding microfuge tubes on ice, ensuring the pellet was not disturbed.

#### Needle shear lysis

For needle shear lysis, medium was replaced with 200 µL cold PBS to resuspend the cells. Cells were passed forcibly, approximately 20 times, through a fine 30G needle with additional scraping at the base of the well to aid resuspension. The plate was centrifuged at 500 *xg* for 5 mins to separate the cells and debris from SN. The top 150 μL of SN was collected in low-binding microfuge tubes and stored at 4°C for downstream analyses within 24 hours.

### Magnetic GFP-Trap Agarose Co-IP of TRIM21 and Antibodies

30 μL of each SN (UT, needle, Opti-MEM only) or freshly prepared cell lysate was transferred to new tubes and combined with 10 µL 4x Laemmli loading buffer containing 10mM DTT. This was the whole cell lysate sample which was stored at -20°C for immunoblotting.

In fresh low-binding tubes, 1 mL of each SN sample was added. Where required, 0.01 mg/mL normal human serum (final concentration of total serum proteins) was added to each relevant sample and incubated at 4°C for 2 hours under rotation. Magnetic GFP-trap agarose beads were equilibrated by washing 4 times in 500 μL cold Pierce IP lysis buffer, using a pre-cooled DynaMagnet. After the final wash, beads were resuspended in IP lysis buffer at the original bead volume. 30 μL equilibrated beads was added to each sample for incubation at 4°C for 1 hour under rotation.

Samples were placed on the magnet and beads were washed 4 times with 500 μL cold Pierce IP lysis buffer, each time resuspending them carefully in the buffer. Washed beads were carefully transferred to fresh low-binding tubes and washed once with cold 1 X TBS pH 7.4 (150 mM Tris, 50 mM NaCl). Beads were then resuspended in 15 μL elution buffer (100 mM Glycine, pH 2.5) and incubated for 10 mins on a room temperature rocker, at fast speed. Note, every 2-3 minutes tubes were gently flicked to mix the beads. Final collection tubes were prepared by adding 15 μL neutralising 1M Tris pH 8.0 into each base, placing these tubes on ice. Sample tubes were magnetised and the 15 μL SN was transferred into the neutralising tubes, discarding the beads. 10 µL 4x Laemmli loading buffer containing 10 mM DTT, was added. This was the final eluate sample which was stored at -20°C ready for immunoblotting.

### BN-Page

Media was aspirated and cells were washed once with 200 μL PBS to remove any serum. Cells were lysed in 50 μL prepared NativePAGE™ Sample Buffer. Triplicate cell lysate wells were pooled (150 μL), transferred to low-protein-binding tubes, and centrifuged at maximum speed for 30 mins at 4°C using a table-top microcentrifuge. SN were transferred to new low-protein-binding tubes on ice, without disturbing the pellet. The following relevant materials were added to the 150 μL sample tubes: 0.01 mg/mL (final concentration) of normal human serum or 1:100 final dilutions of patient/HC plasma. Additional tubes were prepared containing 1:100 final dilutions of patient/HC plasma in 150 μL lysis buffer alone. Sample tubes were incubated under rotation at 4°C for 2 hours.

After incubation, 19 μL sample was transferred to fresh Eppendorf tubes and the remaining material was stored at -20°C. 1 μL of 5% G-250 sample additive was added to the samples just prior to loading the gel. 1 μL of 5% G-250 sample additive was also added to 19 μL Invitrogen™ NativeMark™ Unstained Protein Standard.

Samples were run on 4-16% native page gels in XCell SureLock™ Mini-Cell tanks. To prepare the gel for loading, dark blue cathode buffer (prepared 200 mL dark blue cathode buffer; 10 mL 20X NativePAGE™ running buffer, 10 mL NativePAGE™ blue cathode additive, 180 mL dH_2_O) was pipetted into the loading wells. This was in order to visualise the wells and prevent samples flowing out of the wells. Marker or samples were loaded and the inner chamber was filled with dark blue cathode buffer. The outer chamber was filled with 600 mL anode buffer (prepared 1 L; 50 mL of 20X NativePAGE™ running buffer + 950 mL dH_2_O). The gel was run at 150V until the samples could be visualised at approximately 1/3^rd^ down the gel. The run was paused, all the dark blue cathode buffer was pipetted out and replaced with light blue cathode buffer (prepared 200 mL light blue cathode buffer; 10 mL 20X NativePAGE™ running buffer, 1 mL NativePAGE™ blue cathode additive, 189 mL dH_2_O) and the run was continued.

Proteins were transferred onto methanol-activated PVDF membranes using a Trans-blot Turbo (Bio-Rad). After transfer the membrane was incubated in 8% acetic acid for 15 minutes to fix the proteins. The dark blue staining was removed by rinsing 2-3 times with methanol and then TBS-T. Ponceau S was added to visualise the unstained marker bands. The membrane was blocked with 5% milk TBS-T for 1 hr at room temperature.

Antibodies for Native Page immunoblot were against TRIM21 (1:2000) and anti-human IgG HRP conjugate (heavy and light chain) (1:3000). All antibodies were prepared in 5% milk TBS-T, supplemented with 0.02% (w/v) Na azide. Proteins were visualised by ECL detection. Note, between TRIM21 and anti-human IgG detection, the membranes were stripped using Restore™ Western Blot Stripping Buffer.

### IC formation and analysis by FIDA

1 x T75 flask of ∼80% confluent HeLa_TRIM21_w_PRYSPRY-GFP (described above) provided enough cells for IC formation and FIDA analysis. Medium was removed, cells washed with PBS, then resuspended in pre-warmed trypsin-EDTA (3 mL per flask). After resuspension, trypsin was inactivated with ∼7 mL warm complete media, cells collected and centrifuged for 5 mins, 300 *xg* at room temperature. Cell pellets were resuspended in 10 mL PBS, centrifuged again for 5 mins, 300 *xg* at 4°C to remove any remaining media. Pellets were resuspended in cold Native Page buffer (1 mL per T75 flask) and transferred to low-protein-binding tubes. Tubes were incubated at 4°C for 30 mins to ensure complete lysis, then centrifuged at 16,400 *xg*, 5 mins at 4°C. 30 μL SN (enough for FIDA analysis) were distributed into fresh low-protein-binding tubes. Where relevant, antibody plasma was added in a 1:10 ratio (3 μL per sample) and incubated under rotation for 2 hours at 4°C. Samples were not stored, and were analysed fresh by FIDA.

FIDA experiments were performed using a Fida 1 instrument employing light-emitting-diode induced fluorescence detection with an excitation wavelength of 480 nm and emission wavelengths >515 nm from Fida Biosystems ApS. Standard silica capillaries with inner diameter 75 µm, outer diameter 375 µm, total length 100 cm, length to detection window 84 cm, from Fida Biosystems were used for all experiments. The capillaries were coated with high-sensitivity (HS) coating reagent prior to use. The following stepwise procedure was applied for the FIDA experiments. First, ultrapure water was used for flushing the HS-coated capillary at 3500 millibar (mbar) for 60 s. The analyte sample (assay buffer or assay buffer with antibody plasma) was then filled into the capillary at 3500 mbar for 40 s followed by injection of the indicator (TRIM21-GFP alone or premixed with antibody plasma) at 50 mbar for 10 s. Lastly, the indicator was mobilized to the fluorescence detector with analyte sample (assay buffer or assay buffer with antibody plasma) at either 300, 100, 50 or 25 mbar. The equilibrium dispersion profile (Taylorgram) was recorded in the latter step. The Taylorgrams were analysed using FIDA software (V 2.37, Fida Biosystems). The samples and buffer vials were kept in a temperature-controlled environment at 4°C inside the instrument. The capillary was kept at 25°C. The viscosity of the assay buffer at 25°C was measured using a Lovis 2000 ME microviscometer. Viscosity compensation was included in the data analyses to correct for any changes in viscosity caused by the addition of the antibody plasma. All samples were analysed in at least triplicate.

According to the experimental evaluation tool in the Fida analysis software, species are outside Taylor conditions above certain mbar values. These are as follows: at 300 mbar species fall outside Taylor conditions when their hydrodynamic radius is larger than ∼17 nm; at 100 mbar > ∼53 nm; at 50 mbar > ∼106 nm; and > ∼212 nm at 25 mbar. Baseline shift detection at ∼1/2 the retention time of the main peak (∼1/2tR), allowed us to estimate complex sizes (hydrodynamic radius) and see soluble aggregates outside of Taylor conditions.

### Custom Sandwich GFP ELISA

Corning® 96-well High Bind Microplates were coated with 100 μL of the following antibodies diluted in PBS in triplicate; normal human serum (5 μg/mL), patient/HC plasma (diluted 1:100, 1:1000, as indicated). PBS-only coating blank control was included for each condition for relative background binding. Plates were coated overnight at 4°C. Between each stage of substrate addition, ELISA plates were aspirated and washed in PBS-T.

Plates were blocked with 300 µL/ well blocking buffer and incubated for one hour at room temperature. The following cell lysate samples in IP-lysis buffer were diluted in blocking buffer; HeLa_TRIM21_w_PRYSPRY (1:10, 1:100, 1:1000, 1:10,000) and HeLa_TRIM21_NO_PRYSPRY (1:10, 1:100, 1:1000, 1:10,000). The following SN/control samples were added neat; Opti-MEM only, UT cell SN, apoptosis, pyroptosis SN. 100 µL of samples were added to designated wells. Plates were incubated for two hours at room temperature with gentle shaking (∼500 RPM).

100 µL anti-GFP-HRP detection antibody (diluted to 200 ng/mL in blocking buffer) was added/ well. Plates were incubated for 1 hour at room temperature with shaking. 100 µL of TMB substrate solution was added to each well. Plates were incubated for 30 minutes in the dark at room temperature. 100 µL of stop solution (2M sulphuric acid) was added to each well and absorbance was measured at 450 and 570 nm. Final OD was calculated as A450-A570 values. For each relevant condition, the specific PBS-only blank was subtracted to calculate final OD values.

### Micro-purification of TRIM21-GFP and IC Formation

1 x T75 flask of ∼80% confluent HeLa_TRIM21_w_PRYSPRY-GFP or HeLa_TRIM21_FcMUT-GFP HeLa (described above) provided enough cells for 2 samples/conditions after protein elution. Medium was removed, cells washed and pelleted as previously described in “IC formation for FIDA” section. Pellets were resuspended in cold Pierce IP lysis buffer (2 mL per T75 flask). Lysates were split into 1 mL volumes in fresh low-protein-binding microfuge tubes on ice. Tubes were rotated at 4°C for 30 mins, then centrifuged at 16,400 *xg*, 5 mins at 4°C. SN were transferred to fresh low-binding microfuge tubes on ice, ensuring the pellet was not disturbed.

Magnetic GFP-trap agarose beads were equilibrated and proteins eluted as described under “Magnetic GFP-Trap Agarose Co-IP of TRIM21 and Antibodies”. The 30 μL elution volume was brought to 200 μL with 170 μL Opti-MEM. To relevant samples, the following antibodies were added: normal human serum (0.01 mg/mL final concentration), patient/HC plasma (1:100 dilution). To generate TRIM21-OVA ICs, the following initial concentrations/dilutions were used; OVA (10,000 ng/mL), rabbit α-OVA antiserum (1:100), normal rabbit serum (1:100). Samples were serially diluted to obtain 10x solutions, which after feeding gave final OVA concentrations of 10 ng/mL, 100 ng/mL and 1000 ng/mL. Samples were incubated at 4°C for 2 hours under rotation for IC formation.

### BMDM/HMDM Uptake Assays

Complete DMEM (BMDM) or RPMI (HMDM) was aspirated, and IFNγ primed macrophages washed once with 200 μL pre-warmed Opti-MEM to remove remaining serum. Media was replaced with 100 μL pre-warmed Opti-MEM. For degradation experiments, 2 hours prior to feeding, media was replaced with 100 μL pre-warmed Opti-MEM, with treatment conditions containing 10 nM BafA1 or 10 μM MG132. The 4°C plate condition was pre-chilled on ice for ∼10 mins prior to the experiment. TRIM21-GFP proteins were eluted from GFP-trap, and complexed with normal human serum or patient plasma, as described above. To the top of each relevant well, 10 μL of the following GFP-trap eluted proteins was added; TRIM21_w_PRYSPRY-GFP (+/- normal human serum), TRIM21_FcMUT-GFP (+/- normal human serum), TRIM21_w_PRYSPRY-GFP (+/- HC plasma), TRIM21_w_PRYSPRY-GFP (+/- patient plasma). Plates were returned to the incubator or maintained on ice. At relevant collection time points (5, 15, 30, 60 mins), media was aspirated and cells washed 3 times with excess cold PBS to remove unbound TRIM21/antibodies. Cells were lysed in 30 µL lysis buffer (66 mM tris-Cl (pH 7.4), 2% SDS) and triplicate wells were pooled for analysis. Lysates were collected and stored at -20 °C.

### BMDM Ub-TUBE Assay

BMDMs were plated at 2 × 10^6^ cells per well in TC-treated flat bottom 6-well plates, and primed with 100 ng/mL mIFNγ. 2 hours prior to feeding, BMDMs were washed and replaced with 1 mL OptiMEM +/- 10 μM MG132. Cells were fed with 100 μL of GFP-trap eluted TRIM21 (+/- complexed with antibodies), as previously described for the uptake assays. Cells were fed as indicated and incubated for 60 minutes, then media was aspirated and cells were gently washed 3 times with excess room temperature PBS to remove unbound TRIM21/antibodies without disturbing the cells. 500 μL cold Pierce IP-lysis buffer was added, and cells harvested with cell scrapers. Lysates were transferred into sterile low-protein binding microfuge tubes stored on ice. Tubes were rotated at 4°C for 30 mins, then centrifuged at 16,400 *xg*, 5 mins at 4°C. SN were transferred to fresh low-binding microfuge tubes on ice, ensuring the pellet was not disturbed. 30 μL of each SN sample was transferred to a fresh tube and combined with 10 µL 4x Laemmli loading buffer containing 10 mM DTT. This whole cell lysate sample was stored at -20°C for immunoblotting. Magnetic Ub-TUBE beads were equilibrated and proteins were eluted using the same methods as previously described under “Magnetic GFP-Trap Agarose Co-IP of TRIM21 and Antibodies”. To the 30 µL elution volume, 10 µL 4x Laemmli loading buffer containing 10 mM DTT, was added. This was the final eluate sample which was stored at -20°C ready for immunoblotting.

### HMDM Activation Assay

IFNγ primed HMDMs were fed with 10 μL of GFP-trap eluted TRIM21 (+/- complexed with antibodies), as described above for the uptake assays. Cells were fed, incubated for 6 hours, and SN were collected in fresh 96-well plates and stored at -20 °C. SN were analysed using the TNFα Human ELISA Kit, according to the manufacturer’s instructions.

### RNAseq

IFNγ primed HMDMs were plated as previously described for uptake and activation assays, but media was replaced with 80 μL OptiMEM prior to feeding. Cells were fed with 20 μL (1:5 ratio) of GFP-trap eluted TRIM21 (+/- complexed with antibodies) and incubated for 6 hours. SN were aspirated and cells were washed with room temperature PBS. Cells were lysed in 100 μL RLT buffer (with β-mercaptoethanol) per well. Individual donor conditions were pooled and RNA extracted using the RNeasy micro kit, according to manufacturer’s instructions.

Library preparation and RNAseq was performed by Novogene Inc. using the HiSeq 4000 (Illumina, paired-end mode, read length 150bp). Raw reads were aligned on the GRCh37 reference genome (release 103) using STAR mapper (v 2.7.11) (*107*). Read counts were estimated with RSEM (v 1.1.17) (*108*). Count normalization and differential gene expression analysis were performed using DESeq2 (v 1.44.0) (*109*). Gene set enrichment analysis was performed with the GSEA software (v 3.0, Broad Institute) using predefined gene sets from the Molecular Signatures Database (MSigDB 6.2). Classical enrichment statistics with 1,000 permutations were used to determine significant enrichment within gene sets. Heat maps were generated using the pheatmap package (v 1.0.12) (*110*), and volcano plots were generated using the EnhancedVolcano package (v 1.16.0) (*111*).

### Antigen Presentation Assay

IFNγ primed BMDMs were plated at 5 × 10^4^ cells per well in TC-treated U-bottom 96-well plates, primed with mIFNγ as described above. BMDMs were fed with 10 μL of materials as described for uptake assays. Feeding materials were as follows; OVA, α-OVA, OVA-α-OVA complex, GFP-trap eluted TRIM21, TRIM21 and OVA, TRIM21-α-OVA-OVA complex, TRIM21-normal rabbit serum and OVA (see “Micro-purification of TRIM21-GFP and IC Formation” for details). Plates were incubated for 5.5 hours at 37°C, after which SIINFEKL peptide (50 ng/mL) was added to relevant positive control wells. Plates were incubated for an additional 30 mins. Media was aspirated, cells were washed and media replaced with 100 μL pre-warmed RPMI (T cell formulation described in Media and Supplements). Isolated OT-I cells, resuspended in complete RPMI, were added at 1 × 10^5^ cells per well to achieve a 2:1 coculture ratio with the seeded BMDMs. The cells were incubated overnight at 37 °C in 5% CO_2_ before staining and flow cytometry.

### Flow cytometry

Cell suspensions collected from OT-I/macrophage co-cultures were stained in 96-well V-bottom plates. Live/dead staining and blocking was performed using Zombie NIR Fixable Viability Kit (1:1000) and TruStain FcXTM (1:200) diluted in 1x PBS. For extracellular staining, cells were incubated with fluorochrome-conjugated antibodies diluted in flow cytometry buffer (all 1:400) for 30 mins on ice. The following Biolegend primary antibodies were used: Alexa Fluor® 700 anti-mouse CD3, Brilliant Violet 650™ anti-mouse CD8α, PerCP/Cyanine5.5 anti-mouse CD45.1, APC anti-mouse TCR Vβ5.1, 5.2, PE anti-mouse CD69. Cells were fixed in 4% paraformaldehyde for 15 minutes on ice. Fixed cells were washed and resuspended in flow cytometry buffer for flow cytometric analysis. Data were collected on a BD LSRII using BD FACSDiva software and analysed on FlowJo^TM^ software (v10.9.0). Graphs and statistical tests were completed on GraphPad Prism (v10.0.3).

### H&E and Cell DIVE Slide Preparation and Imaging

5 μm thick formalin-fixed, paraffin-embedded (FFPE) tonsillar tissue mounted on TOMO slides was purchased from the Oxford Centre for Histopathology Research. 5 μm SG biopsy slides were obtained from the OASIS cohort (Birmingham). One SG slide for each patient was H&E stained, and imaged using a Zeiss Axioscan 7, at 20X magnification.

Antigen retrieval was performed according to the GE Cell DIVE system protocol (*72*). Slides were deparaffinised by overnight baking at 60 °C. Slides were cleared in xylene, rehydrated in decreasing concentrations of ethanol (100%, 95%, 70% and 50%) and permeabilised in 0.3% Triton-X-100 in PBS (1X) for 10 mins. Slides were finally washed in PBS for 5 mins. A NxGen decloaking chamber at 110 °C was used for antigen retrieval. Slides were placed in room temperature citrate solution, pH 6.0, and once the chamber reached 70 °C, the citrate protocol was started (including 4 minutes 110 °C). Slides were then transferred to tris-antigen retrieval working solution pH 9.0 (10 mM Tris base, 1 mM EDTA solution, 0.05% Tween 20) for 20 minutes in the chamber, before being left on the bench for a further 10 mins at room temperature. After cooling, slides were washed 4 times in PBS for 5 mins each, with gentle agitation on an orbital shaker.

Slides were blocked overnight 4 °C in blocking solution (PBS, 3% BSA, 10% donkey serum). Slides were washed for an additional 5 mins in PBS with orbital shaking. Next slides were blocked for 1 hour at room temperature with human FcR block, diluted 1:200 in antibody diluent (PBS, 3% BSA). Slides were washed 3 times in PBS for 5 mins each, then stained with DAPI solution for 15 mins with orbital shaking, before an additional wash in PBS for 5 mins. Finally, slides were coverslipped in mounting media (90% glycerol, 4% propyl gallate).

Background imaging was performed according to the GE Cell DIVE system manufacturer’s instructions on the IN Cell 2500HS. Briefly, an overall scan plan was acquired at 10X magnification. Entire tissue sections were selected for imaging and the background, and subsequent imaging rounds were imaged at 20X magnification. Background imaging enabled removal of autofluorescence from subsequent imaging rounds.

For staining, slides were de-coverslipped and washed 3 times in PBS, each for 5 mins with orbital shaking. Primary antibodies were diluted in antibody diluent at the following dilutions; TRIM21 (10 μg/mL), IgD, CD68, CD3, CD163, IgG, pan-cadherin, IgA, CD66b, CD11c, HLA-DR, CK8, CD20 (5 μg/mL), CK13 (2.5 μg/mL). Slides were stained with primary and directly-conjugated antibodies overnight at 4 °C before 3 washes in PBS, each for 5 mins with orbital shaking. Donkey anti-rabbit AF555 secondary antibody was diluted 1:500 and slides were incubated at room temperature for 1 hour in secondary antibody. Slides were finally washed 3 times in PBS for 5 mins, with orbital shaking and re-coverslipped in mounting media for staining imaging rounds.

To inactivate the dyes, slides were de-coverslipped and bleached 2 times. For this, slides were placed in bleaching solution (0.1 M NaHCO_3_ pH 11.2, 3% H_2_O_2_) which is freshly prepared every time, before a 1 minute PBS wash. Slides were placed in DAPI solution for a further 2 minutes at room temperature, with orbital shaking. Slides were then re-coverslipped in mounting media.

Iterative rounds of stained and bleached imaging was performed according to the manufacturer’s instructions. Briefly, slides were configured to subtract autofluorescence (from the background scan). Images were acquired using the 20X objective, aligned to previous scan rounds and merged. Images were imported to the QuPath software (*112*) for visualisation and selecting areas of interest.

### Statistical Analyses

Immunoblots were quantified using Image J, according to the protocol by Stael et al. 2022 (*113*). Results were analysed using GraphPad Prism 9 by ordinary one-way ANOVA, multiple mean comparisons between either: UT and treated conditions, or pre-selected pairs. Post-hoc testing used either Fisher’s least significant difference or Tukey’s post-test. Asterisks denote the following; * P ≤ 0.05, ** P ≤ 0.01, *** P ≤ 0.001.

## Abbreviations

BafA1: (Bafilomycin A1)
BMDMs: (Bone-marrow derived macrophages)
BN-PAGE: (Blue-Native Page)
CK: (Cytokeratin)
co-IP: (co-immunoprecipitation)
ctDNA: (Calf thymus DNA)
EBV: (Epstein-Barr virus)
FcR: (Fc receptor)
FcγRs: (Fcγ receptors)
FFPE: (Formalin-fixed, paraffin-embedded)
FIDA: (Flow induced dispersion analysis)
HC: (Healthy control)
hIFNγ: (human interferon-γ)
HMDMs: (Human monocyte derived macrophages)
HS: (High-sensitivity)
IC: (Immune complex/complexes)
IFNs: (Interferons)
IL: (Interleukin)
IP: (Immunoprecipitation)
LDH: (Lactate dehydrogenase)
LIPS: (Luciferase immunoprecipitation system)
LPS: (Lipopolysaccharide)
LU: (Light units)
Luc: (Luciferase)
M-CSF: (Macrophage colony-stimulating factor)
MFI: (Mean fluorescence intensity)
mbar: (Millibar)
mIFNγ: (murine interferon-γ)
mTOR: (mammalian target of rapamycin)
Nec: (Necrostatin)
NES: (Nuclear export sequence)
OVA: (Ovalbumin)
OASIS: (Optimising Assessment of Sjögren’s Syndrome)
PBMCs: (Peripheral blood mononuclear cells)
RUH: (Rockefeller University Hospital)
RNAseq: (Ribonucleic acid sequencing)
SLC: (solute carrier)
SG/SGs: (Salivary glands)
SjD: (Sjögren’s disease)
SM: (SMAC mimetic)
SN: (Supernatants)
Tcrb-V5.1: (T-cell receptor beta, variable 5.1)
TBS-T: (Tris-buffered saline with Tween)
TLR: (Toll-like receptor)
TRIM21-mEGFP: (Tripartite motif-containing protein 21-monomeric EGFP)
Ub-TUBE: (Ubiquitin-Tandem Ubiquitin Binding Entity)
WT: (Wild-type)

## Supplementary Materials

List of materials file

Figs. S1 to S6

Movie S1

## Acknowledgments

We thank members of the LB Dustin and JS Bezbradica laboratories for helpful discussions. Thank you to all at the Kennedy Institute of Rheumatology Digital Pathology Omics Core and Cell Dynamics Core for their technical expertise and support in multicolour imaging and flow cytometry. Henrietta Lacks, and the HeLa cell line that was established from her tumour cells without her knowledge or consent in 1951, have made significant contributions to scientific progress and advances in human health. We are grateful to Henrietta Lacks, now deceased, and to her surviving family members for their contributions to biomedical research.

## Funding

Kennedy Trust for Rheumatology Research Studentship KENN 192001 (ELJ)

Henni Mester Fellowship, University College, University of Oxford (ELJ)

Oxford-Bristol Myers Squibb Celgene Research Fellowship Programme (LBD)

Kennedy Trust for Rheumatology Research KENN212202 (JSB)

Medical Research Council MR/W001217/1 (JSB)

Kennedy Trust for Rheumatology Research KENN202112 (AG)

NIHR Birmingham Biomedical Research Centre (BAF)

The views expressed in this publication are those of the authors and not necessarily those of the NHS, the NIHR or the Department of Health

## Author contributions

Conceptualization: ELJ, BDM, JSB, LBD

Methodology: ELJ, BDM, MDJ, CH, SH, REG, GH, SN

Investigation: ELJ, BDM, MDJ, CH, GH

Visualisation: ELJ, MDJ

Funding acquisition: AG, JSB, LBD

Project administration: ELJ, JSB, LBD

Supervision: BAF, AG, JSB, LBD

Writing – original draft: ELJ

Writing – review & editing: ELJ, JSB, LBD

### Competing interests

SjD work in LBD’s laboratory is funded in part by a fellowship from the Oxford-Bristol Myers Squibb Celgene Research Fellowship Programme. The funders had no part in the preparation of this manuscript. The authors declare that no conflict of interest exists.

### Data and materials availability

RNASeq data will be deposited in a GEO database and accession number provided before publication. All other data are available in the main text or the supplementary materials.

## Supplementary Figures

[MOVIE LINK]

**Movie S1. TRIM21 is retained in apoptotic blebs.** HeLa overexpressing TRIM21_w_PRYSPRY-GFP were treated with 1 μM of sirDNA dye 30 minutes before imaging. At the microscope, apoptosis-inducing drugs (1 μM ABT737, 10 μM MCL1i) were added 5 minutes into 4 hour live confocal microscopy recording. Images were taken at 1 minute intervals, collated in FIJI/ImageJ and exported as a compressed 24 second recording. Scale bar, 50 µm. Recording is representative of 3 independent live time-lapse microscopy experiments.

**Fig. S1.**
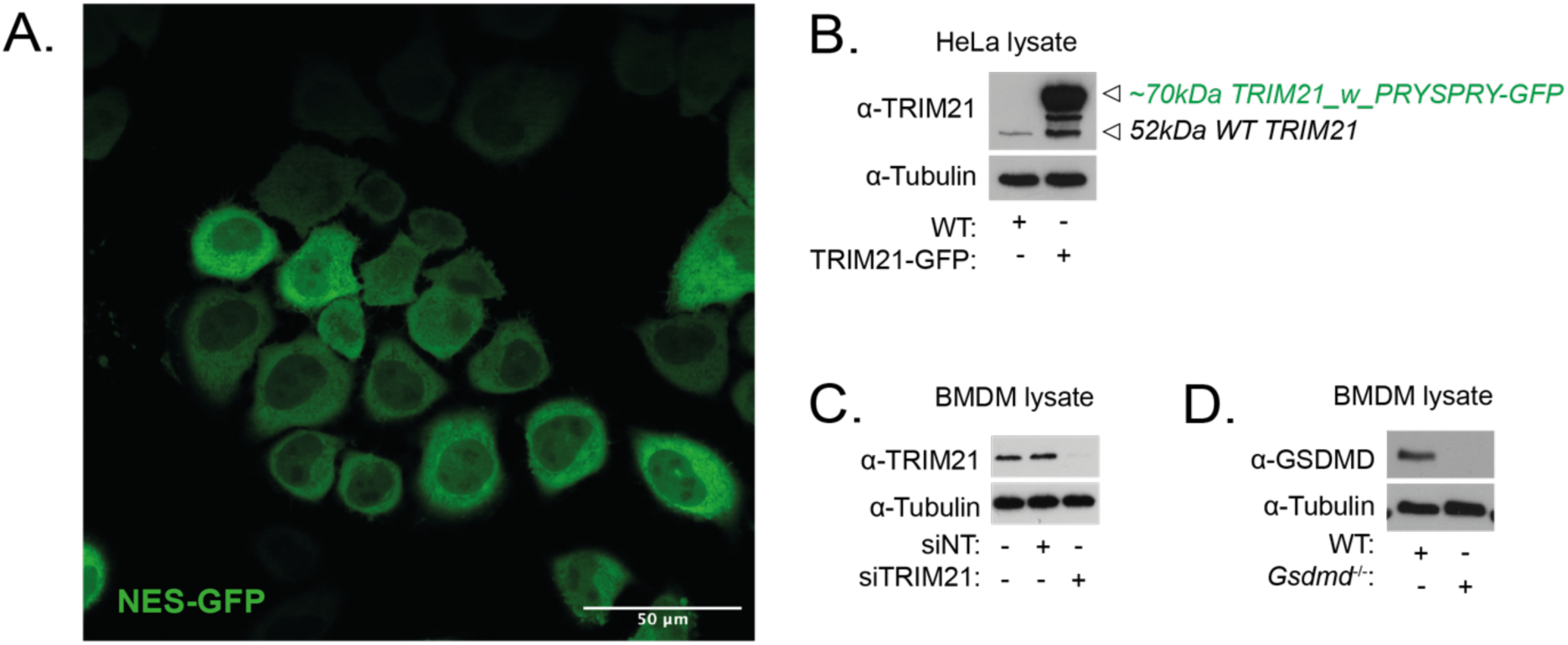
TRIM21 overexpression and antibody controls. **(A)** HeLa overexpressing HIV_NES-GFP imaged by live confocal microscopy. Scale bar, 50 µm. **(B)** WT and TRIM21_w_PRYSPRY-GFP HeLa lysates were collected. **(C)** For siRNA knockdown, BMDMs were transfected with 4 pmol of either siGENOME non-targeting (NT) siRNA control pools, or ON-TARGETplus mouse Trim21 siRNA. Lysates were collected after 72 hours. **(B, C)** Lysates were analysed by immunoblot using the Abcam anti-TRIM21/SSA rabbit monoclonal antibody (ab207728), 1:2000 dilution. **(D)** WT and *Gsdmd^-/-^* mouse lysates were collected and analysed by immunoblot for GSDMD protein expression.

**Fig. S2.**
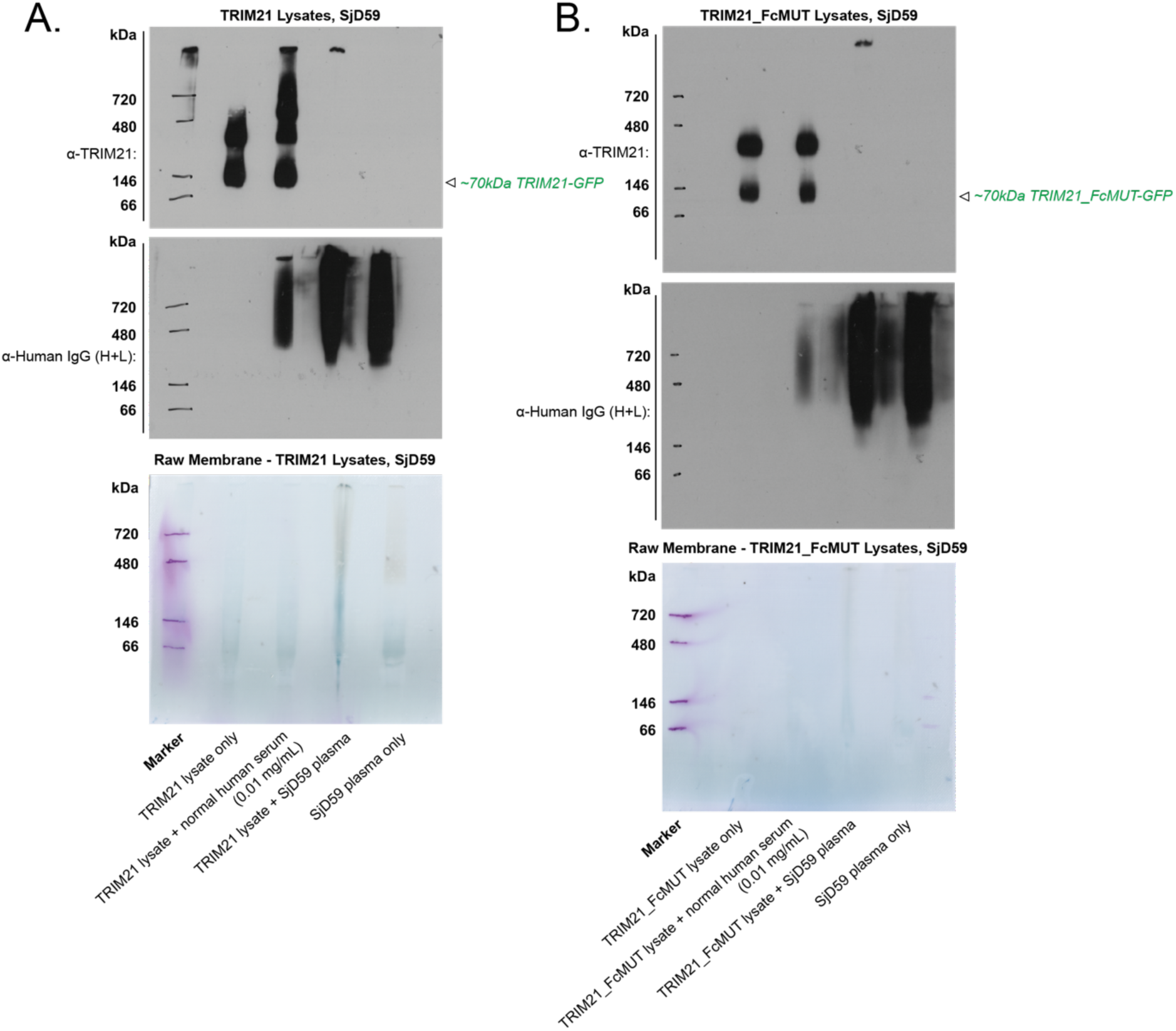
SjD59 BN-Page gels. Non-denaturing BN-PAGE separation of **(A)** TRIM21_w_PRYSPRY-GFP lysates **(B)** TRIM21_FcMUT-GFP lysates from HeLa, complexed with normal human serum (0.01 mg/mL), SjD59 patient plasma or plasma alone. Raw membrane scans and immunoblots are shown. Molecular weight markers are indicated.

**Fig. S3.**
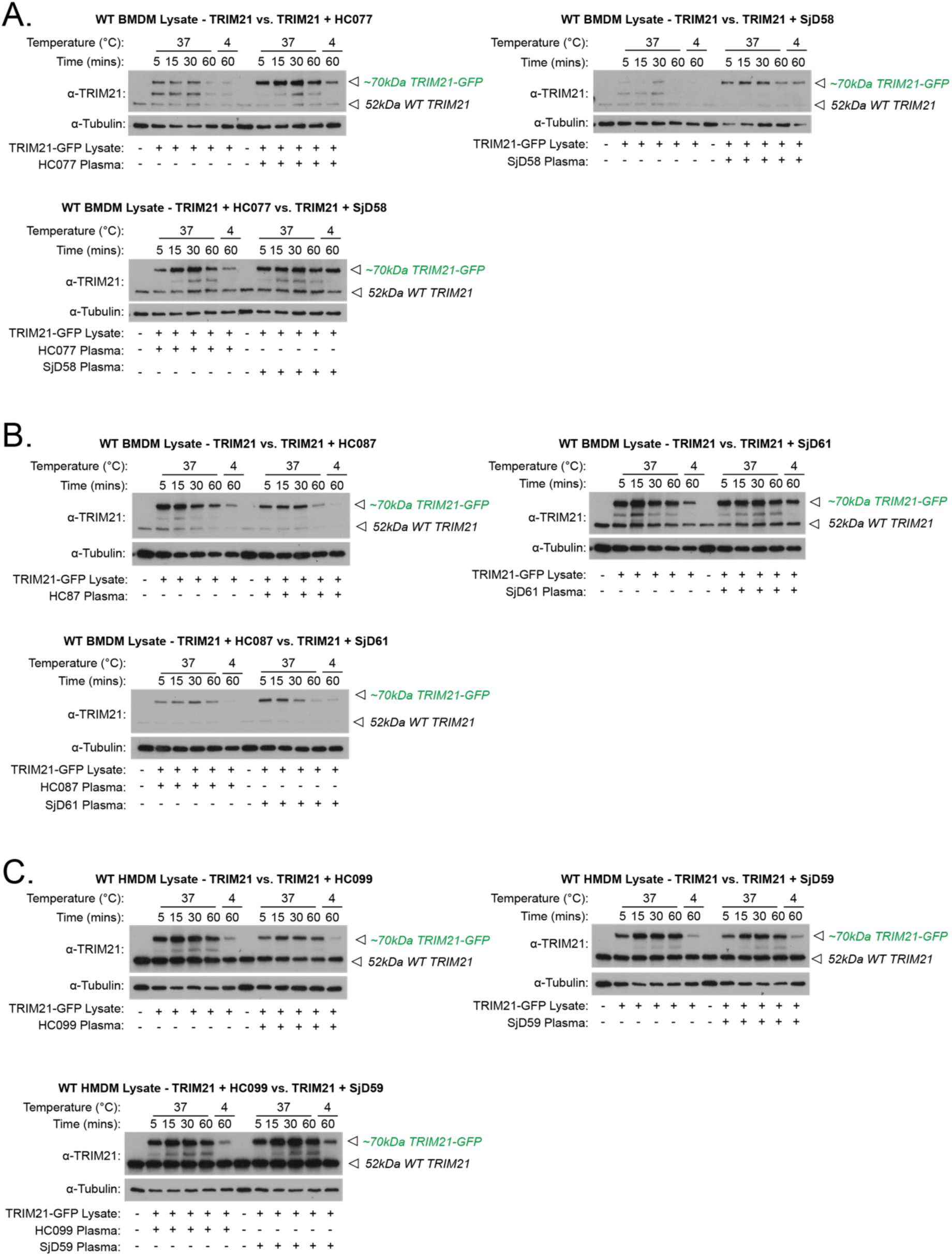
Macrophage uptake of TRIM21 complexed with patient plasma. IFNγ-stimulated BMDMs were fed with TRIM21_w_PRYSPRY alone, or complexed with plasma (HC/SjD) (1:100 dilution). Uptake reactions were stopped after 5, 15, 30 and 60 mins. Cells kept on ice (4°C) provided control background binding levels. Lysates were collected and analysed by immunoblot, comparing TRIM21 alone +/- complexed with patient plasma. The following three independent biological replicate pairings were analysed: **(A)** HC077 vs. SjD58, **(B)** HC087 vs. SjD61, **(C)** HC099 vs. SjD59.

**Fig. S4.**
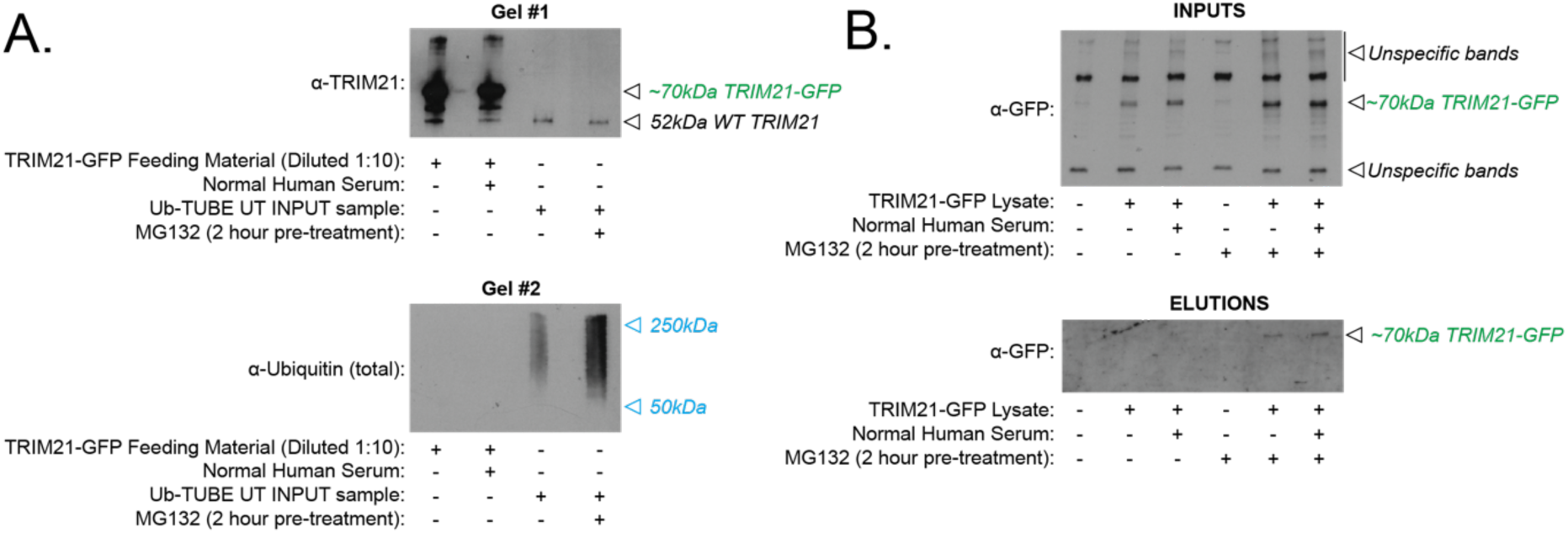
Ub-TUBE controls. **(A)** GFP-Trap eluted TRIM21 (+/- complexed with normal human serum) feeding material, and Ub-TUBE IP inputs (+/- 10 μM MG132) were analysed by immunoblot. TRIM21 and total ubiquitin levels were detected using replicate membranes. **(B)** Ub-TUBE inputs and elutions were analysed by immunoblot, detecting the GFP tag of TRIM21_w_PRYSPRY-GFP.

**Fig. S5.**
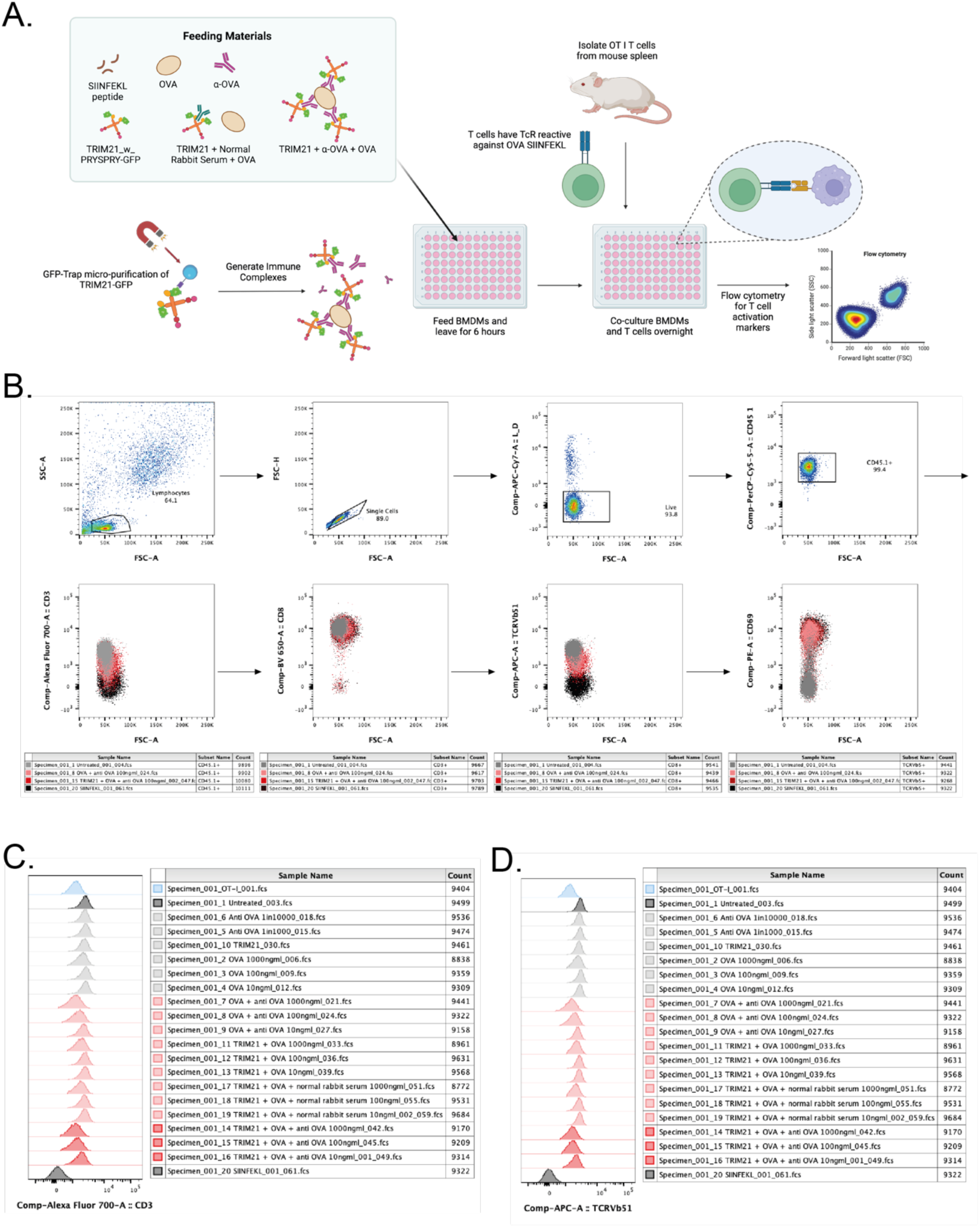
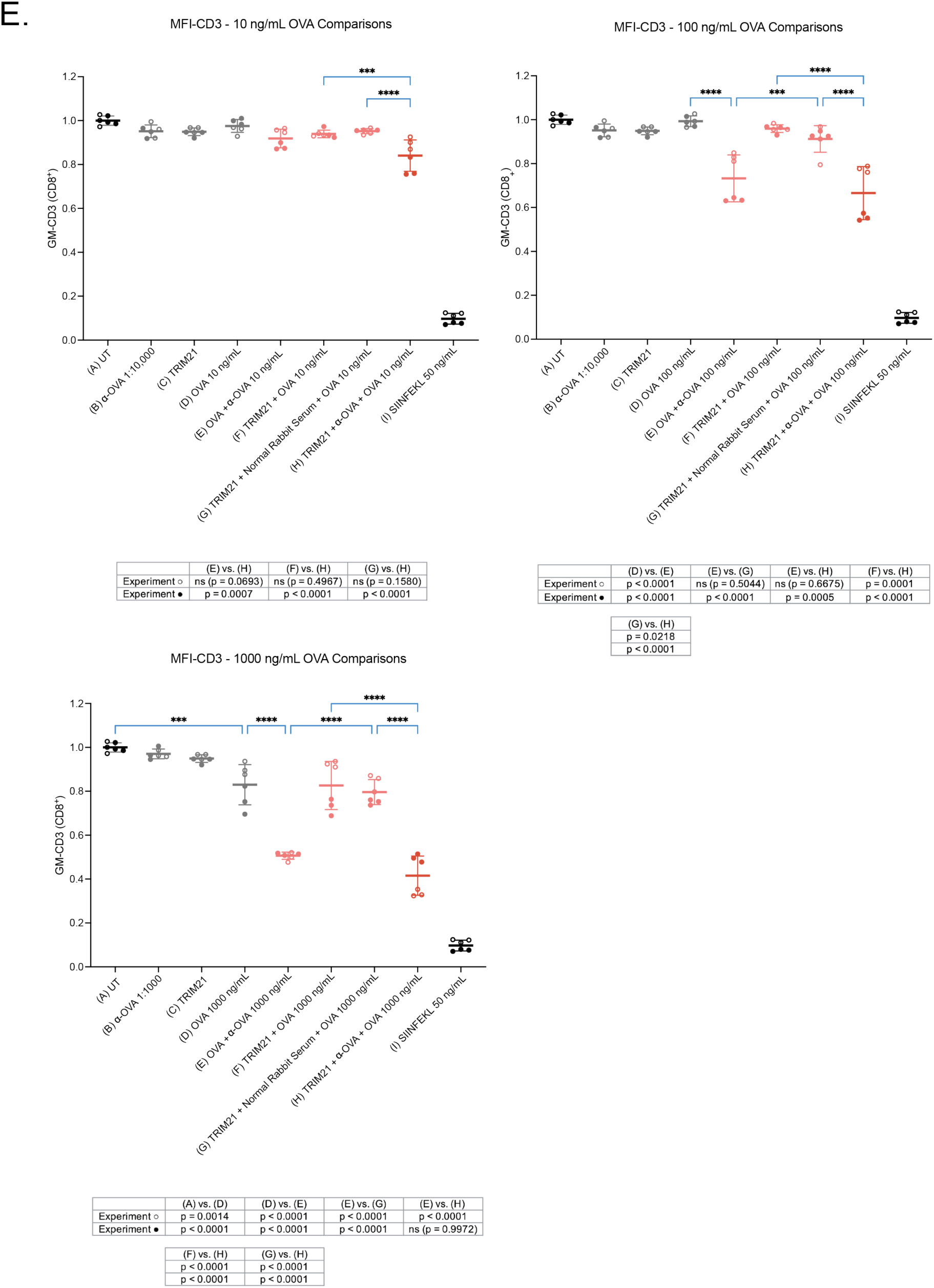
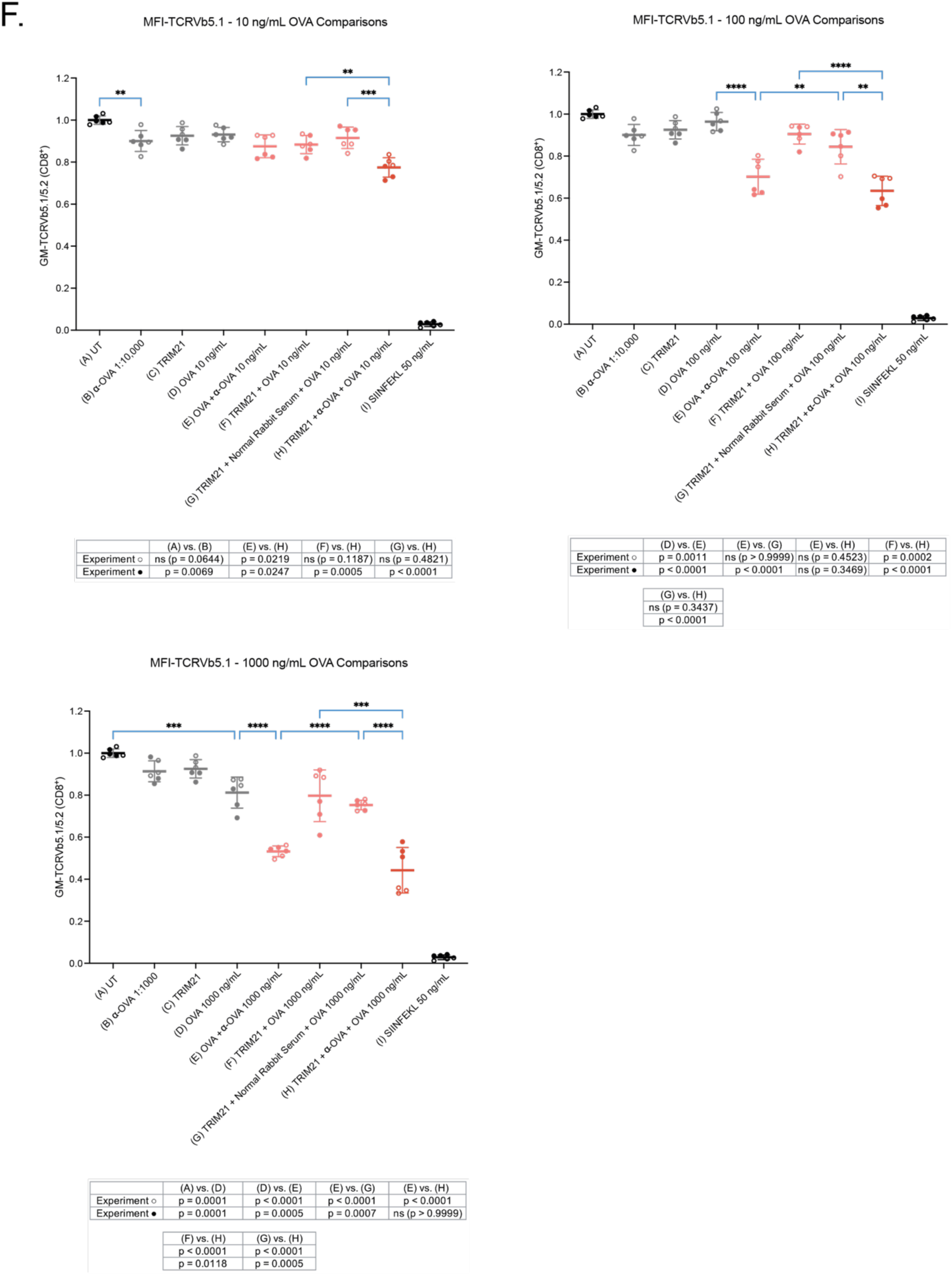
TRIM21-OVA antigen presentation. BMDMs were stimulated with 100 ng/mL mIFNγ for ∼24 hours and fed with 10 μL of the following materials: OVA (+/- rabbit α-OVA), GFP-trap eluted TRIM21 (+/- OVA, OVA/α-OVA or OVA/normal rabbit serum), 50 ng/mL SIINFEKL peptide. **(A)** Schematic illustration of antigen presentation experimental set-up. **(B)** Example gating strategy to select for CD45.1^+^ CD8α^+^ CD3^+^ OT-I cells and then for Tcrb-V5.1 and CD69 expression. Gating shown for T cells co-cultured with the following BMDM experimental conditions; UT, OVA/α-OVA (100 ng/mL), TRIM21 + OVA/α-OVA (100 ng/mL), and SIINFEKL (50 ng/mL). Representative histograms for **(C)** CD3 and **(D)** TcrbV5.1 expression. Results shown for one of two biologically independent experiments. **(E)** CD3 MFI and **(F)** Tcrb-V5.1 MFI, was determined for a range of OVA concentrations; 10 ng/mL, 100 ng/mL and 1000 ng/mL. Results from two biologically independent experiments were pooled, normalised to UT, and analysed by one-way ANOVA and Tukey’s post-hoc test. Duplicate experiments are distinguished by filled or empty symbols. Data represented as mean + SD for each condition. Tables denoting specific p values for each experiment are shown.* P ≤ 0.05, ** P ≤ 0.01, *** P ≤ 0.001.

**Fig. S6.**
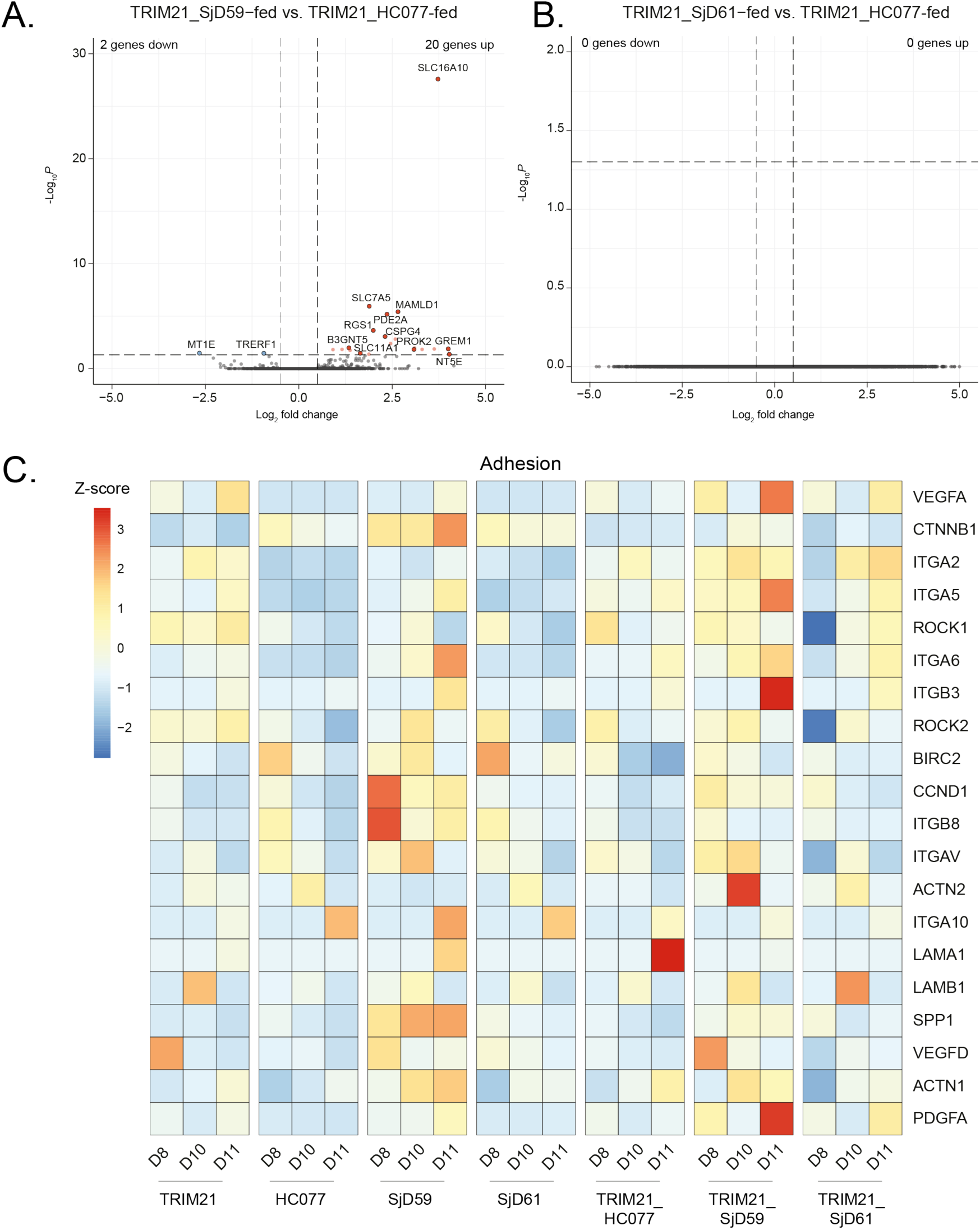
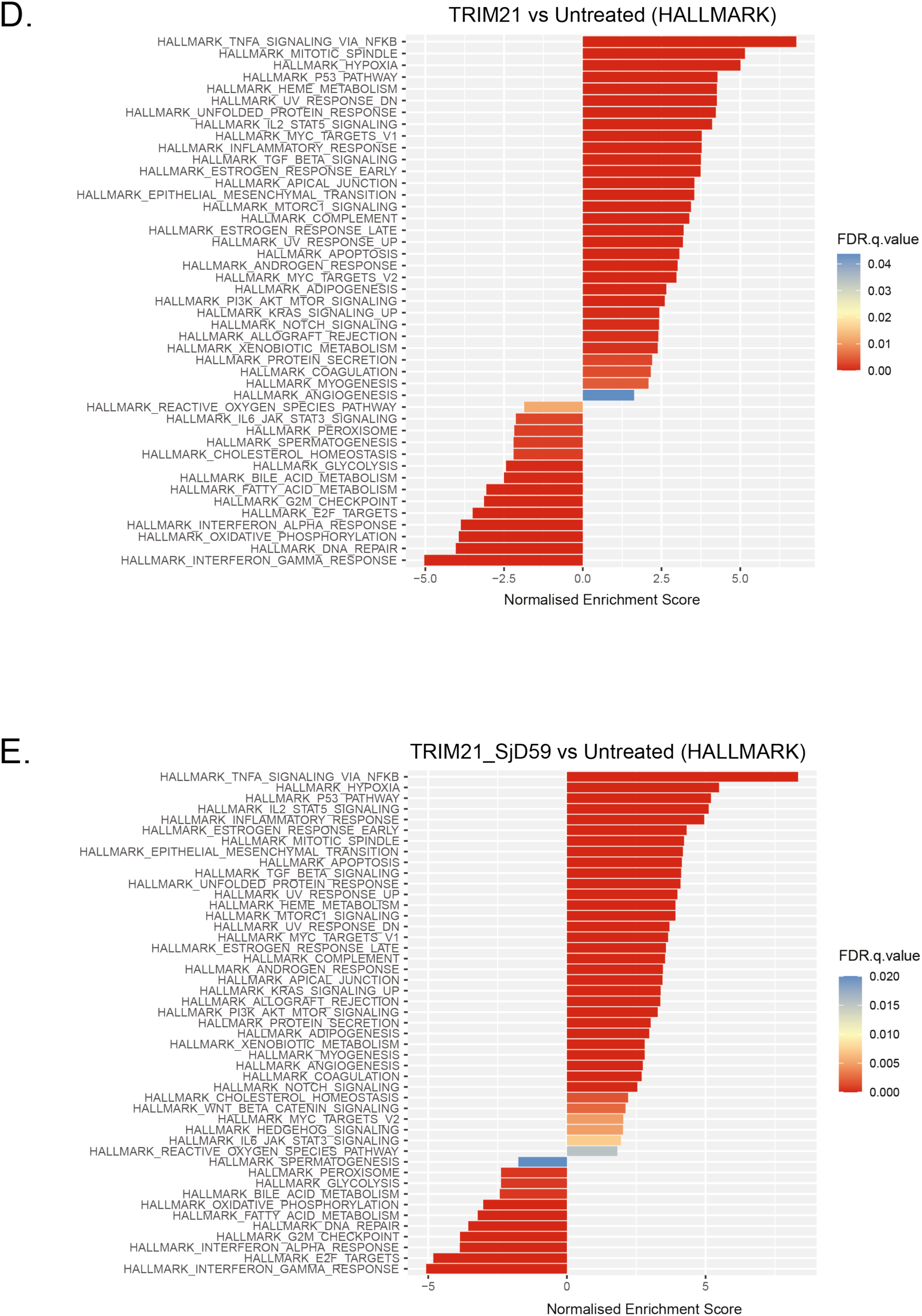
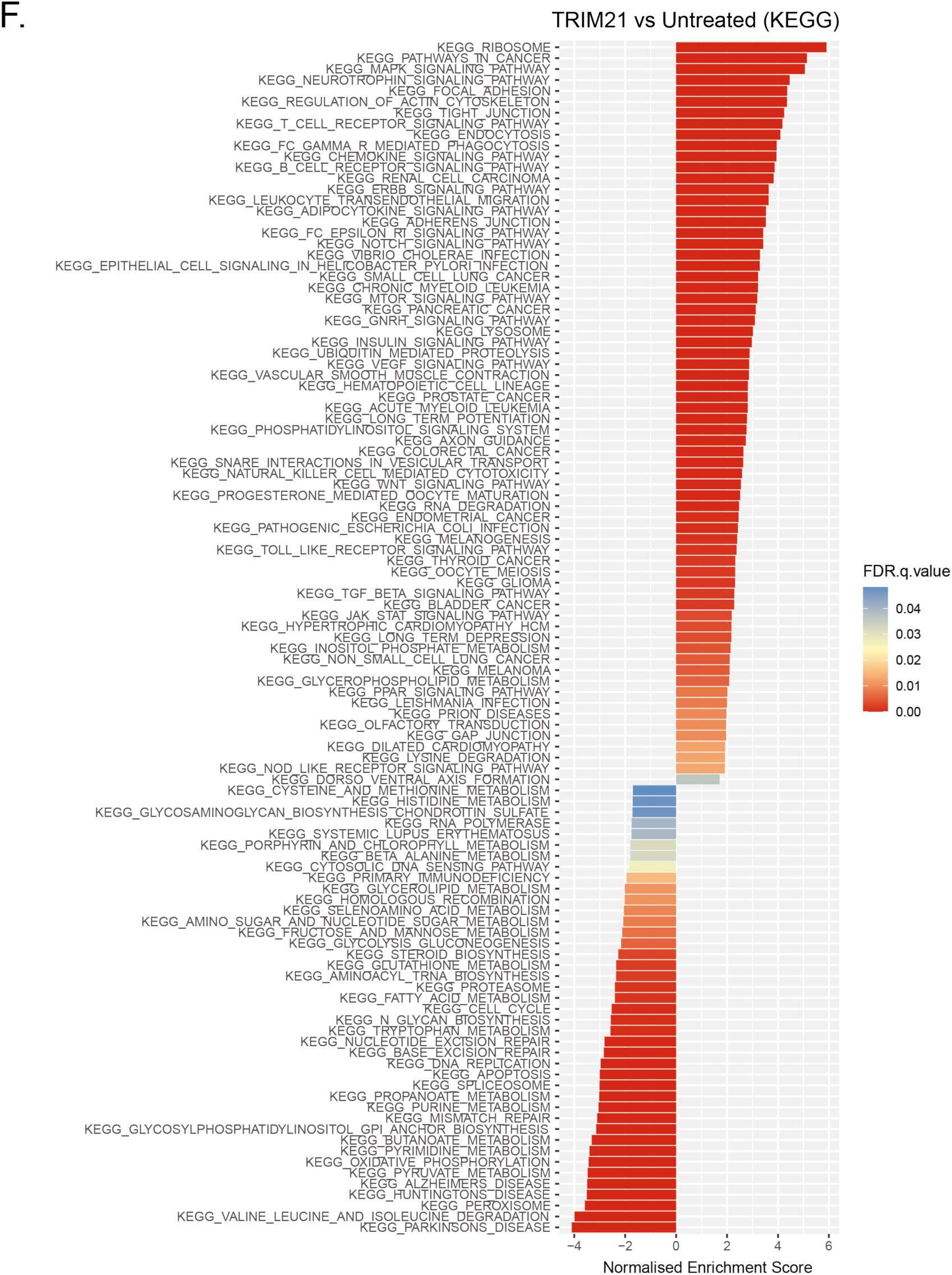
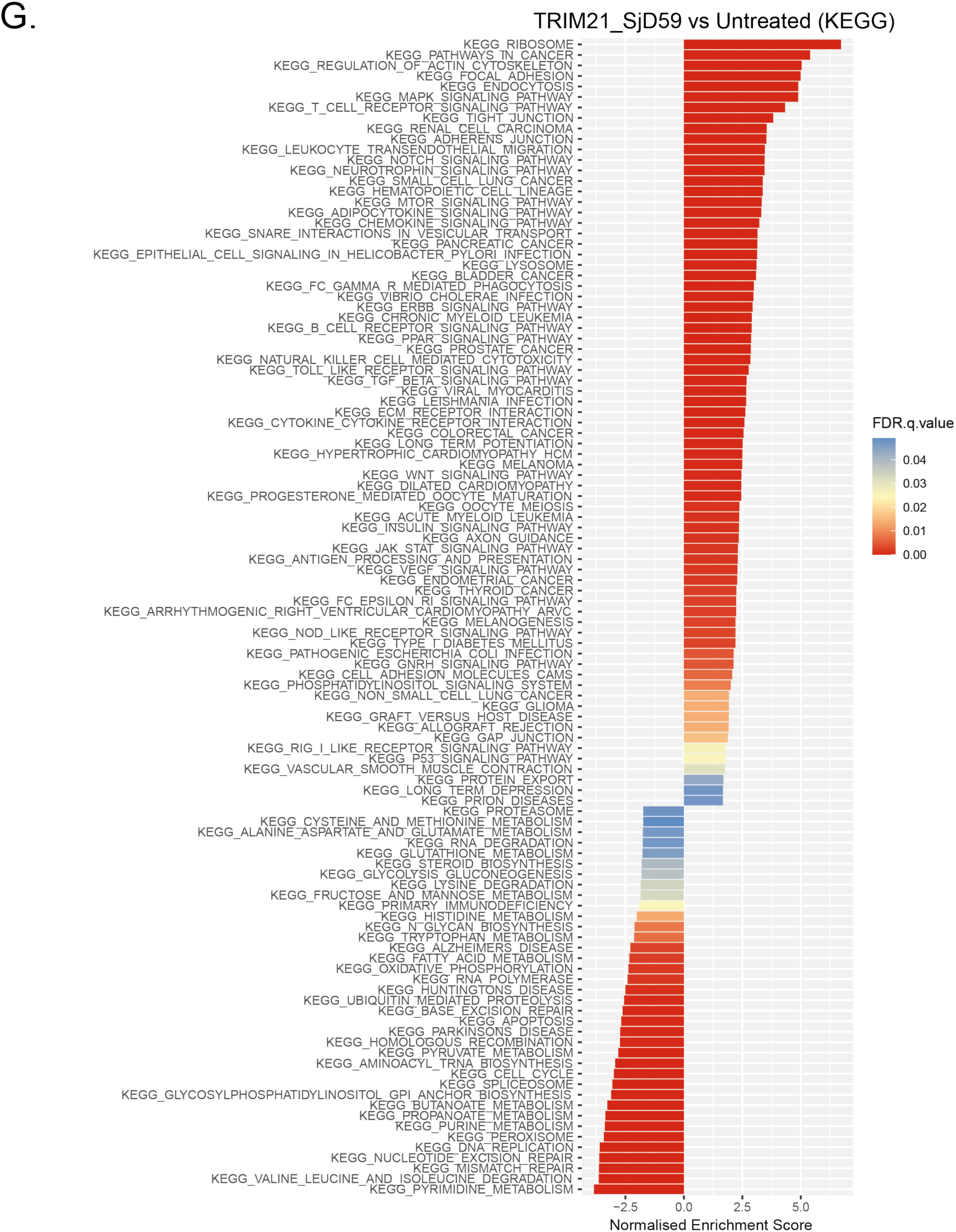
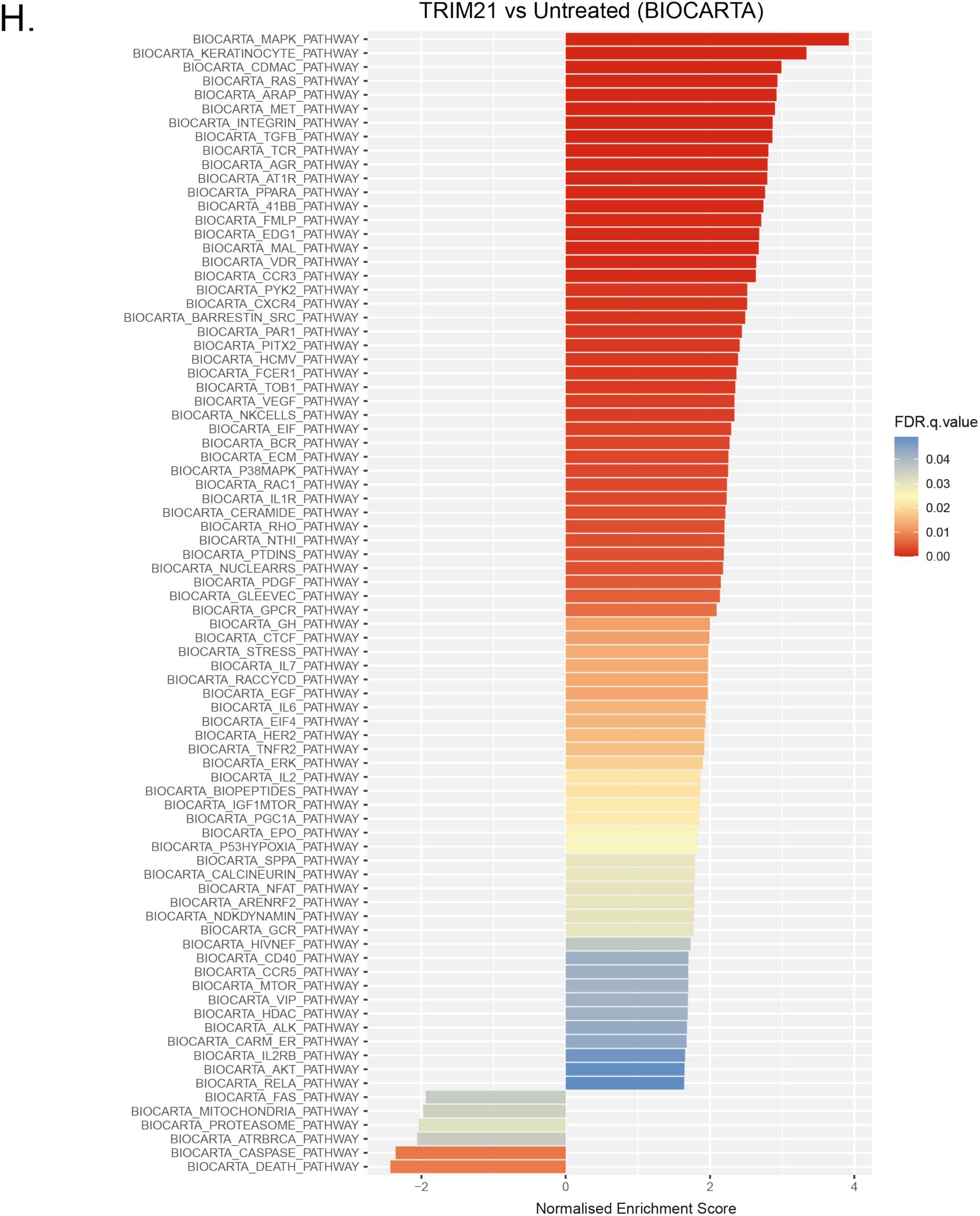
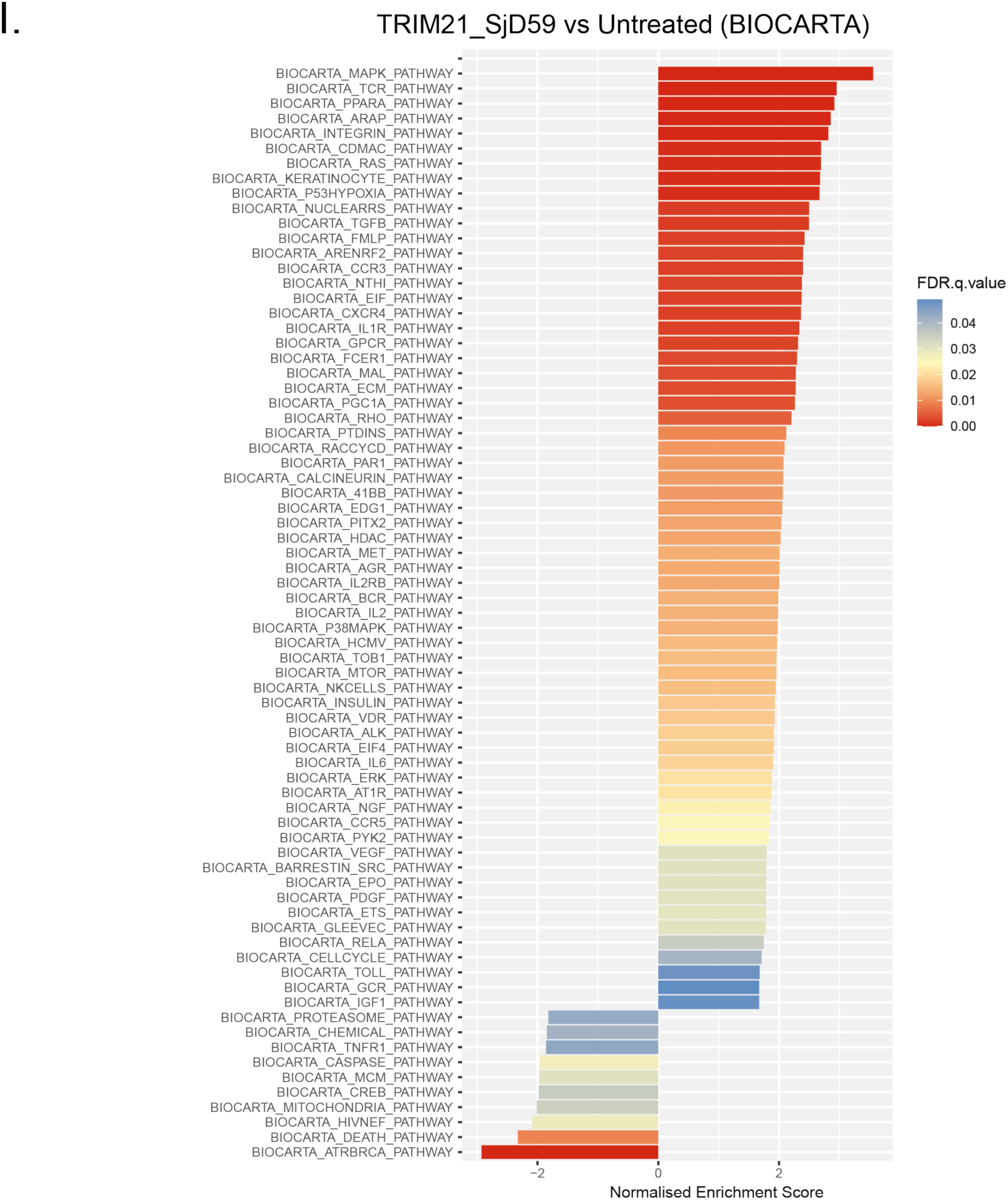
HMDM RNA sequencing after TRIM21 and TRIM21-IC feeding. HMDMs from 3 donors (D8, D10, D11) were fed with 20 μL of GFP-trap eluted TRIM21 (+/- complexed with patient plasma). After 6 hours, RNA was extracted, sequenced, aligned to reference genome and analysed. Significance was calculated using the Wald test (two-tailed), followed by Bonferroni correction. Volcano plots highlight log_2_ fold changes in differentially expressed genes for **(A)** TRIM21_SjD59 vs. TRIM21_HC077 and **(B)** TRIM21_SjD61 vs. TRIM21_HC077 feeding. **(C)** Heatmap showing expression changes in adhesion genes for donors and feeding conditions. **(D-F)** TRIM21 vs. UT and **(G-I)** TRIM21_SjD59 vs. UT gene set enrichment analyses for assigned gene sets.

## Notes

### Competing Interest Statement

The authors have declared no competing interest.

